# Multi-omics integrative approach of senescent endothelial cells and derived extracellular vesicles in a replicative senescence model

**DOI:** 10.1101/2025.11.25.690417

**Authors:** Fátima Milhano Santos, Rafael Ramírez-Carracedo, Sergio Ciordia, Laura Clavaín, Juan Pablo Hernández-Fonseca, Beatriz Martín-Jouve, Guillermo Bodega, Rafael Ramírez, María Piedad Ruiz-Torres, Caridad López-Granero, Marta Ruiz-Ortega, Matilde Alique

**Affiliations:** Cellular and Molecular Biology in Renal and Vascular Pathology Laboratory, Health Research Institute-Fundación Jiménez Díaz University Hospital, Universidad Autónoma de Madrid (IIS-FJD, UAM), 28040 Madrid, Spain; RICORS2040, Madrid, Spain; Departamento Cirugía, Ciencias Médicas y Sociales, Universidad de Alcalá, Alcalá de Henares, 28871, Alcalá de Henares, Madrid, Spain; Functional Proteomics Laboratory, Centro Nacional de Biotecnología, CSIC, Calle Darwin 3, Campus de Cantoblanco, 28049 Madrid, Spain; EGO Genomics, Scientific Park of the University of Salamanca, Adaja Street 4, Building M2, 37185 Villamayor, Spain; Electron Microscopy Unit, Centro Nacional de Biotecnología-CSIC, Campus Universidad Autónoma de Madrid, Madrid, Spain; Biomedicine and Biotechnology Department, Alcala University, 28871 Alcalá de Henares, Madrid, Spain; Departamento de Biología de Sistemas, Universidad de Alcalá, Alcalá de Henares, 28871, Alcalá de Henares, Madrid, Spain; Instituto Ramón y Cajal de Investigación Sanitaria (IRYCIS), 28034 Madrid, Spain; Department of Psychology and Sociology, University of Zaragoza, Teruel, Spain

**Keywords:** Cardiovascular diseases, Endothelial senescence, Extracellular vesicles, miRNA, Multi-omics, Proteomics, Replicative senescence, Transcriptomics, Vascular aging

## Abstract

Aging is a major unmodifiable risk factor for cardiovascular disease (CVD). Replicative endothelial senescence (RES), characterized by permanent cell-cycle arrest and a senescence-associated secretory phenotype (SASP), is a hallmark of vascular aging. However, the molecular mechanisms underlying RES, particularly the role of extracellular vesicles (EVs), remain poorly understood. To this aim, an integrated multi-omics approach (proteomics, mRNA, and miRNA profiling) was applied to characterize senescent and early-passage human umbilical vein endothelial cells (HUVECs) and their secreted EVs. Senescent HUVECs and EVs displayed a canonical senescence profile, including cell-cycle arrest, DDR activation, NF-κB signaling, and SASP induction, alongside marked suppression of RNA metabolism, ribosome biogenesis, and DNA repair. Multi-omics integration has linked endothelial senescence to vascular aging, maladaptive angiogenesis, and ECM remodeling, which are key processes in CVD development. Moreover, senescent EVs propagate senescence by exporting fewer reparative and antioxidant factors while carrying pro-fibrotic, adipogenic, and angiogenic mediators. Multi-omics profiling revealed consistent transcript–protein changes in senescent cells and EVs, defining a robust molecular signature of RES, including endothelial cell-specific markers and post-transcriptional regulators (lncRNAs and miRNAs). Among these, miR-22-3p and miR-126-5p emerged as key modulators in both cells and EVs, with miR-22-3p demonstrating compartment-specific regulation. Integration further uncovered a coordinated regulatory network involving miRNAs (miR-23, miR-335-3p, miR-29 family, miR-590-3p, and miR-126-5p) that were inversely correlated with transcription factors (e.g., REST, KLFs, and ZNFs) and downstream targets, collectively driving endothelial senescence through impaired PI3K–Akt signaling. Although further validation is needed, these findings provide new insights into the molecular mechanisms of RES and open perspectives for modulating secondary senescence and age-related vascular diseases.

## 1. Introduction

Life expectancy has increased markedly in recent decades, although the COVID-19 pandemic briefly reversed this trend. In 2023, the predicted life expectancy at birth in the European Union reached 81.4 years, and the global average is expected to reach 73.3 years by 2024, surpassing pre-pandemic levels.^1–3^ Furthermore, the population aged 60 years and older is anticipated to rise from 1.1 billion in 2023 to 1.4 billion by 2030, underscoring the accelerating global trend toward population aging.^3^ However, healthy life expectancy has not kept pace with overall lifespan growth, largely due to the burden of age-related diseases, such as cardiovascular disease (CVD), neurodegeneration, and chronic kidney disease (CKD).^4^

Aging is a multifactorial process characterized by a decline in cellular function due to cumulative molecular damage, mitochondrial dysfunction, and altered cellular signaling.^5–7^ A hallmark of aging is cellular senescence, a permanent cell-cycle arrest induced by replicative exhaustion, oxidative stress, or DNA damage, in which cells remain metabolically active but lose proliferative capacity.^8–11^ Senescent cells accumulate with age, disrupting tissue homeostasis and regeneration, contributing to inflammation and organ dysfunction.^5,12^ Senescent cells undergo transcriptomic reprogramming that induces a senescence-associated secretory phenotype (SASP), composed of growth factors, proteases, and pro-inflammatory cytokines, which propagate senescence to neighboring cells, amplifying inflammation and tissue damage, a process termed secondary senescence.^13,14^

The secretion of extracellular vesicles (EVs) is one key mechanism of secondary senescence.^10,15,16^ EVs, including exosomes and microvesicles (MVs), are lipid bilayer-enclosed structures that carry bioactive molecules such as proteins, nucleic acids, lipids, and metabolites.^11,17^ Increased scientific interest and recent advances have revealed the high complexity and diversity of these lipid structures^18^, prompting the development of the minimum requirements of the Society of the International Society for Extracellular Vesicles (MISEV2023) guidelines to standardize their classification based on size and biogenesis^19^. By mediating both paracrine and systemic signaling, EVs regulate multiple biological processes, including immune responses, metabolic homeostasis, and cellular communication.^20–22^

Endothelial cells, which form the innermost layer of blood vessels, are essential for vascular function and are among the primary cell types affected by biological aging, making them especially vulnerable to age-related damage.^23,24^ Therefore, the characterization of EVs released by endothelial senescent cells offers valuable insights into age-related diseases, including CVD, serving as a source of potential biomarkers and/or therapeutic targets.^10,11,22,25^ Previous studies from our research group have shown that EVs contribute to the crosstalk between endothelial dysfunction and senescence in cardiovascular complications.^26–30^ EVs secretion has been involved in endothelial dysfunction and vascular damage by promoting sustained inflammation and oxidative stress^30–32^, driving atherosclerosis^33^, and vascular calcification^25,27,28,33^. Our recent findings further support the role of EV-mediated endothelial injury in the development of CKD and CVD. Proteomic analysis of EVs from human umbilical vein endothelial cells (HUVECs) exposed to indoxyl sulfate, a uremic toxin associated with vascular calcification and arterial stiffness, revealed downregulation of extracellular matrix (ECM) integrity proteins and upregulation of fatty acid metabolism pathways, reinforcing the contribution of EVs to endothelial dysfunction and to the CKD–CVD axis.^29^

Aging is the primary risk factor for CVD, contributing to coronary artery disease, hypertension, and atherosclerosis^34^, while the accumulation of senescent cells is strongly linked to increased frailty^35^ and higher CVD prevalence in the elderly^36^. Nevertheless, the mechanisms underlying replicative endothelial senescence (RES), particularly the contribution of EVs to vascular aging, remain poorly understood.^24^ Aging research requires models that capture biological age heterogeneity while predicting outcomes.^37,38^ Among these models, the replicative senescence model, firstly described by Hayflick and Moorhead^8^, provides a robust framework for studying cellular aging. Based on this model, we previously established a RES model using HUVECs, which mirrors the key features of vascular aging, including impaired endothelial function and wound healing.^26^ Senescence is increasingly recognized as a dynamic and heterogeneous state driven by multilayered regulatory networks^39–41^, making multi-omics approaches essential to capture its molecular complexity. Therefore, to elucidate RES mechanisms underlying the vascular aging process, an integrated multi-omics approach (proteomics, transcriptomics, and miRNA profiling) was used to profile senescent HUVECs and their secreted EVs, and compare them with early-passage counterparts. Understanding EV-mediated communication could help identify biomarkers and therapeutic targets to prevent vascular dysfunction and age-related CVD.

## 2. Results

### 2.1. Characterization of the replicative endothelial senescence model and isolation of extracellular vesicles from cultured cells

In our RES model, after 27–38 passages, HUVECs exhibited flattened and enlarged cell bodies with irregularly shaped nuclei, cytoplasmic vacuolization, granular inclusions, and abnormal organelles (Figure S1). These structural alterations were accompanied by a markedly reduced proliferation rate, consistent with previous reports.^26^ The replicative senescence model was further validated by Senescence-Associated β-galactosidase (SA-β-gal) staining, a classical senescence marker, which revealed an increase in SA-β-gal–positive cells in senescent cells compared to early-passage HUVECs (Figure S1), as previously reported.^26^

Following the MISEV2023 guidelines^19^, EVs isolated from early-passage and senescent HUVECs were characterized using several complementary techniques (Figure 1, Supplementary Table 1, and Supplementary Figures S2-6). First, the particle size distribution of the isolated EVs was determined using nanoparticle tracking analysis (NTA) (Supplementary Figure S2 and Table S1). NTA revealed polydisperse EVs with similar size distributions in both early-passage and senescent HUVECs, with no significant differences in the mean or mode of particle size (Figure 1A and Supplementary Figure S2). The presence, morphology, and particle distribution of HUVEC-derived EVs were also assessed using transmission electron microscopy (TEM) (Figures 1B/C and Supplementary Figure S3). TEM confirmed the round morphology and heterogeneous size distribution of EVs under both conditions (Figure 1B). Contrasting with NTA, TEM showed that senescent cells secreted significantly smaller-sized particles (p-value=0.0021) compared to early passage cells (Figure 1C). Overall, TEM analysis consistently revealed smaller EV diameters in both early-passage and senescent HUVECs when compared with NTA. This discrepancy likely reflects the sample dehydration and fixation steps inherent to TEM preparation, which are known to induce EV shrinkage, thereby accentuating the apparent size differences between the vesicle populations. These observations are consistent with previous reports indicating that NTA provides a more reliable technique for estimating EV size (mean and mode).^42^

**Figure 1.**
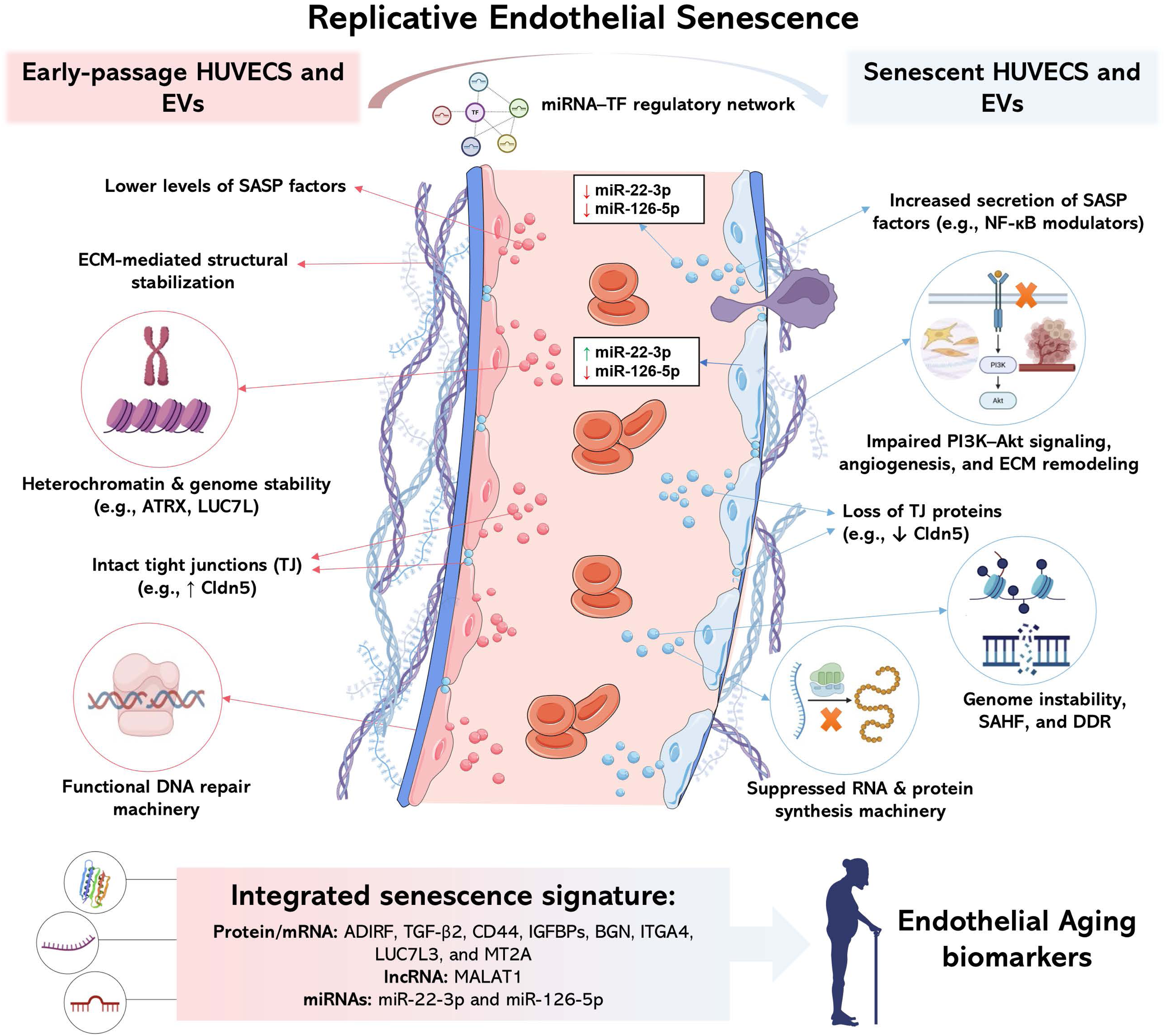
Characterization of extracellular vesicles (EVs) derived from senescent and early-passage endothelial cells was performed using nanoparticle tracking analysis (NTA), transmission electron microscopy (TEM), and flow cytometry (FC). (A) NTA of representative pools from early-passage (pool #1) and senescent (pool #4) EVs showing particle concentration and size distribution; mean and mode sizes are shown. (B) Representative TEM images displaying round morphology and heterogeneous size distribution (scale bars = 1 μm, 200 nm, and 200 nm). (C) Bar graph showing quantitative differences in EV diameter by TEM. (D) Bar graph showing the quantitative differences in the expression of tetraspanins (CD9, CD63, and CD81) were analyzed using FC. (E) FC analysis of EV markers, defining EVs as Tag-It Violet (TIV)+ events. The expression of tetraspanins (CD9, CD63, and CD81) is shown as representative dot plots comparing early-passage and senescent EVs. Statistical analysis: normality was assessed using the Shapiro–Wilk test; parametric and non-parametric data were analyzed using unpaired t-tests or Mann–Whitney tests, respectively (**p < 0.01).

Furthermore, isolated EVs were characterized by flow cytometry (FC) to quantify TIV+ events (TIV+ positive particles) and assess the expression of specific EV markers, including tetraspanins CD9, CD63, and CD8 (Figure 1D/E and Supplementary Figure S4 and Table S1, respectively). Considering the total number of TIV+ events expressing each tetraspanin, senescent EVs showed a higher expression of tetraspanins than EVs derived from early passage cells, but only CD9 was significantly enriched (p-value=0.0016) (Figure 1D).

The presence of canonical EV markers (CD9, CD63, CD81, TSG101, FLOT1/2) in both groups was also confirmed by western blotting (Supplementary Figure S6a) and LC-MS/MS (Supplementary Figures S5 and Supplementary Table 2). Comparison with the Vesiclopedia compendium^43^ confirmed a strong overlap between the EV proteome dataset and previously reported EV proteins (Supplementary Figure S5b). Of the 8,547 proteins deposited in the Vesiclopedia compendium, 3,666 were detected in our EV proteome dataset, including 1,771 endothelial proteins and 3,439 previously reported in exosomes.

Following the MISEV2023 guidelines^19^ and our previous work^26,29^, we confirmed the presence of small EVs and MVs using techniques like NTA and TEM. In addition, specific EV markers, such as tetraspanins, were characterized by proteomics and FC, showing an enrichment of CD9⁺ EVs in senescent conditions. These technical validations ensured the accurate characterization of EV populations before the subsequent multi-omics analysis.

### 2.2. Changes in the proteome of HUVECs and secreted EVs associated with replicative endothelial senescence

A label-free quantitative (LFQ) strategy was applied to compare the proteomes of senescent and early passage HUVECs, as well as the proteomes of their corresponding EVs (Figure 2A). Of a total of 6,315 identified protein groups (89,344 peptide groups, 6,351 proteins; FDR≤1%), 128 were differentially expressed between senescent and early-passage HUVECs (76 overexpressed and 52 underexpressed; adjusted p-value ≤0.05), including 14 exclusively detected in senescent and 32 in early-passage HUVECs (Supplementary Table 2, Figure 2B). Regarding EV proteomic analysis, 4,676 protein groups (48,615 peptides; 4,746 proteins, FDR ≤1%) were identified. Most of these proteins (4,379 protein groups) overlapped with the HUVEC proteome, while 1972 and 367 were exclusively identified in HUVEC and isolated EVs, respectively (Supplementary Table 2 and Figure S7a). Comparing senescent with early-passage EVs, 244 proteins were differentially expressed, including 58 overexpressed proteins (42 unique to senescent EVs) and 186 underexpressed proteins (158 unique to early-passage EVs) (Figure 2C). Principal component analysis (PCA) showed a clear separation between the senescent and early-passage states in both cells (Figure 2D) and EVs (Figure 2E).

**Figure 2.**
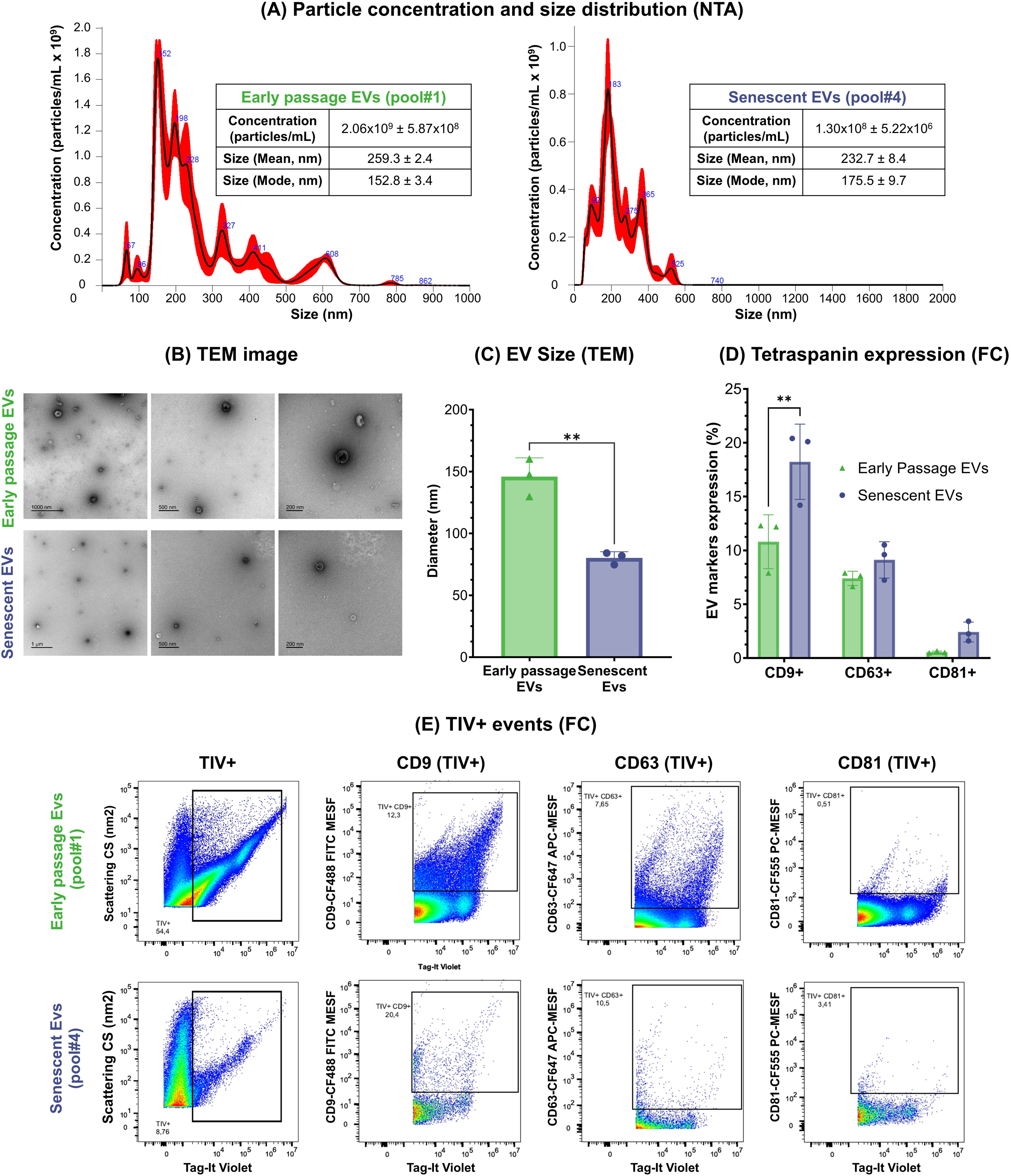
Proteomic profiling of senescent and early-passage endothelial cells and their extracellular vesicles (EVs) in a replicative senescence (RES) model. (A) Experimental workflow of RES model culture, EV isolation, and proteomic analysis by LC–MS/MS. (B-C) Volcano plots showing differentially expressed proteins (DEPs) in senescent compared to early-passage (B) HUVECs and (C) EVs. Significantly regulated proteins (q-value≤0.05) are highlighted, with log2 ratio≥1 indicating overexpressed proteins and log2 ratio≤–1 indicating underexpressed proteins. Non-significant proteins are shown in gray. (D-E) Principal component analysis (PCA) of DEPs showing clear discrimination between senescent and early-passage (D) HUVECs and (E) EV profiles.

Functional enrichment analyses were performed to gain insights into the RES phenotype and its associated biological roles and pathways (Figure 3 and Supplementary Table 3 and Figure S7a and S7b). The top functional terms associated with proteins overexpressed in senescent cells included complement and coagulation cascades, hemostasis, acute-phase response, regulation of insulin-like growth factor (IGF) transport and uptake, ECM organization, and cell adhesion (Figures 3A/C). Many of these processes were also enriched in EVs , albeit less significantly (Figure 3B/D and Supplementary Figure 7a). EVs also showed enrichment in the regulation of inflammation by resistin (RETN) and the production of transforming growth factor beta (TGF-β) (Figure 3B), a cytokine detected only in senescent cells and EVs (Supplementary Table 2). According to the TRRUST analysis, SP1 was identified as a common transcriptional regulator in both cells and EVs, whereas ATF2 and NF-κB specifically regulated the proteins overexpressed in senescent EVs (Supplementary Table 3). Both cells and EVs also showed enrichment in canonical senescence processes, including senescence-associated heterochromatin foci (SAHF), SASP, DNA Damage/telomere stress-induced senescence, and oxidative stress-induced senescence (Figures 3A-D).

**Figure 3.**
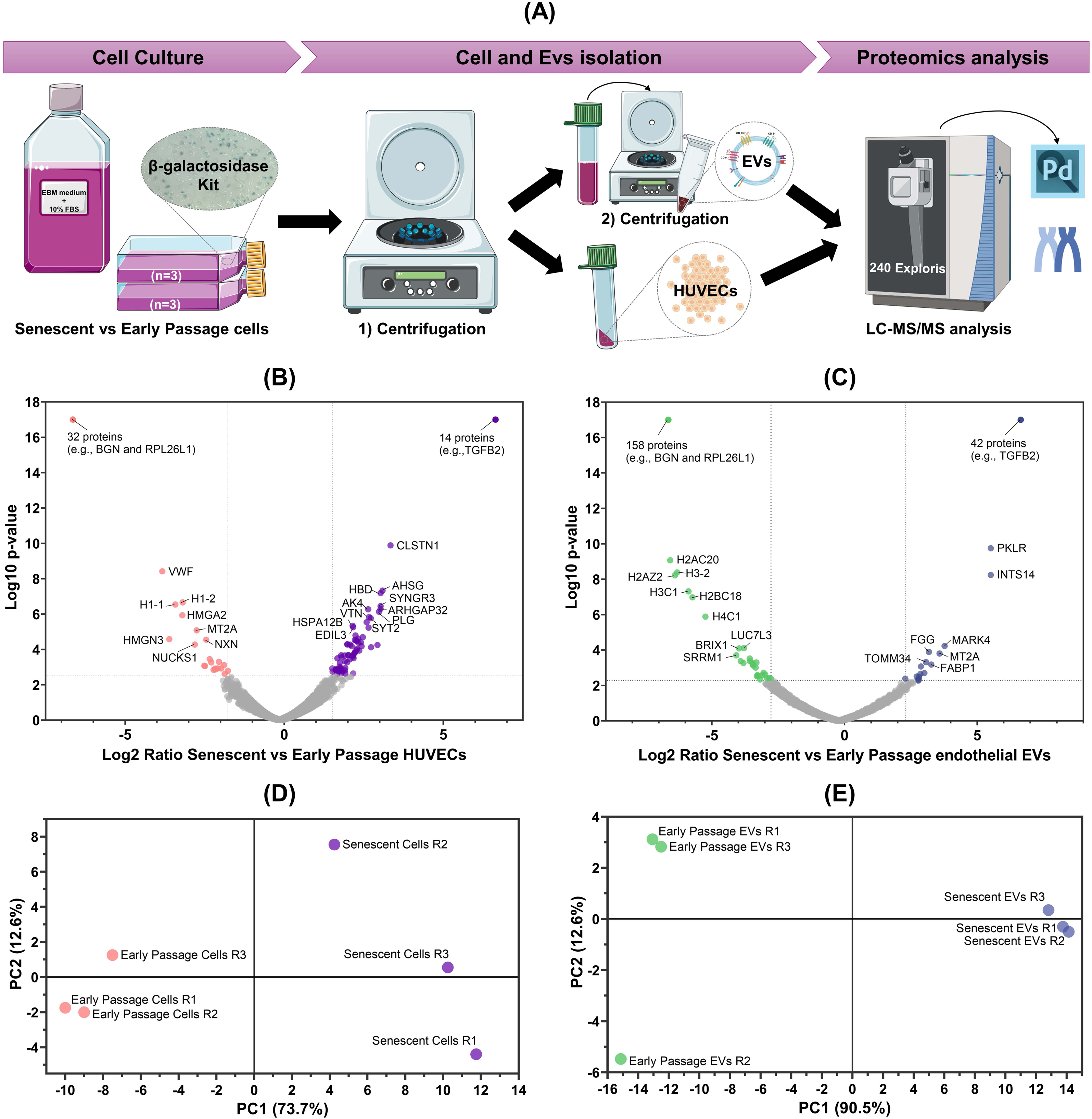
Functional enrichment and protein–protein interaction (PPI) networks of differentially expressed proteins (DEPs) in senescent compared to early-passage endothelial cells and extracellular vesicles (EVs). (A–B) Heatmaps of functional enrichment analysis of DEPs in (A) endothelial cells and (B) extracellular vesicles (EVs), based on Gene Ontology (GO), KEGG, and Reactome pathways using Metascape. (C-D) Protein–protein interaction (PPI) networks of DEPs in (C) endothelial cells and (D) extracellular vesicles (EVs), generated with STRING at high confidence (score = 0.70). The top functional terms are highlighted in different colors within the networks. (E–F) Heatmaps representing disease association analysis of DEPs in (E) endothelial cells and (F) extracellular vesicles (EVs), based on DisGeNET using MetaScape.

In contrast, proteins underexpressed in senescent HUVECs were mainly linked to cholesterol biosynthesis and regulation of canonical Wnt signaling (Figure 3A and Supplementary Table 3), while those in EVs were associated with RNA and DNA metabolism, chromatin organization, and large ribosomal subunit biogenesis (Figure 3B/D).

Endothelial aging and activation of senescence are correlated with the emergence of various age-related diseases. Therefore, we compared our proteomics dataset with the DisGeNET database^44^, a comprehensive repository of gene–disease associations, to assess whether these proteins were linked to known pathologies (Supplementary Table 3). The upregulated proteins in senescent HUVECs and their respective EVs were significantly correlated with inflammation, diabetes, and several CVDs (Figures 3E/F). Many of these proteins are SASP components, including TGF-β2, CD44, alpha-2-macroglobulin, complement factors (C3, C4A), coagulation proteins (e.g., prothrombin), and fibrinolysis regulators (e.g., Tissue-type plasminogen activator [tPA or PLAT]) (Supplementary Table 7).

### 2.3. Changes in the transcriptome of HUVECs and secreted EVs associated with replicative endothelial senescence

To identify transcriptome signatures of endothelial aging in the replicative senescence model, RNA sequencing was performed on senescent and early-passage HUVECs and in their corresponding EVs (Figure 4A and Supplementary Table 4). Comparing senescent with early-passage HUVECs 1,782 differentially expressed genes (DEGs) were identified, including 947 upregulated and 835 downregulated (|log2FC|≥1, q-value≤0.05) (Figure 4B). Given the low transcript yield in EVs, less stringent criteria were applied (|log2FC|≥1, q-value≤0.25), identifying only 22 DEGs in senescent compared with early-passage EVs (3 upregulated, 19 downregulated) (Figure 4C). Using PCA, DEGs distinguished senescent from early-passage phenotypes in both cells (Figure 4D) and EVs (Figure 4DE).

**Figure 4.**
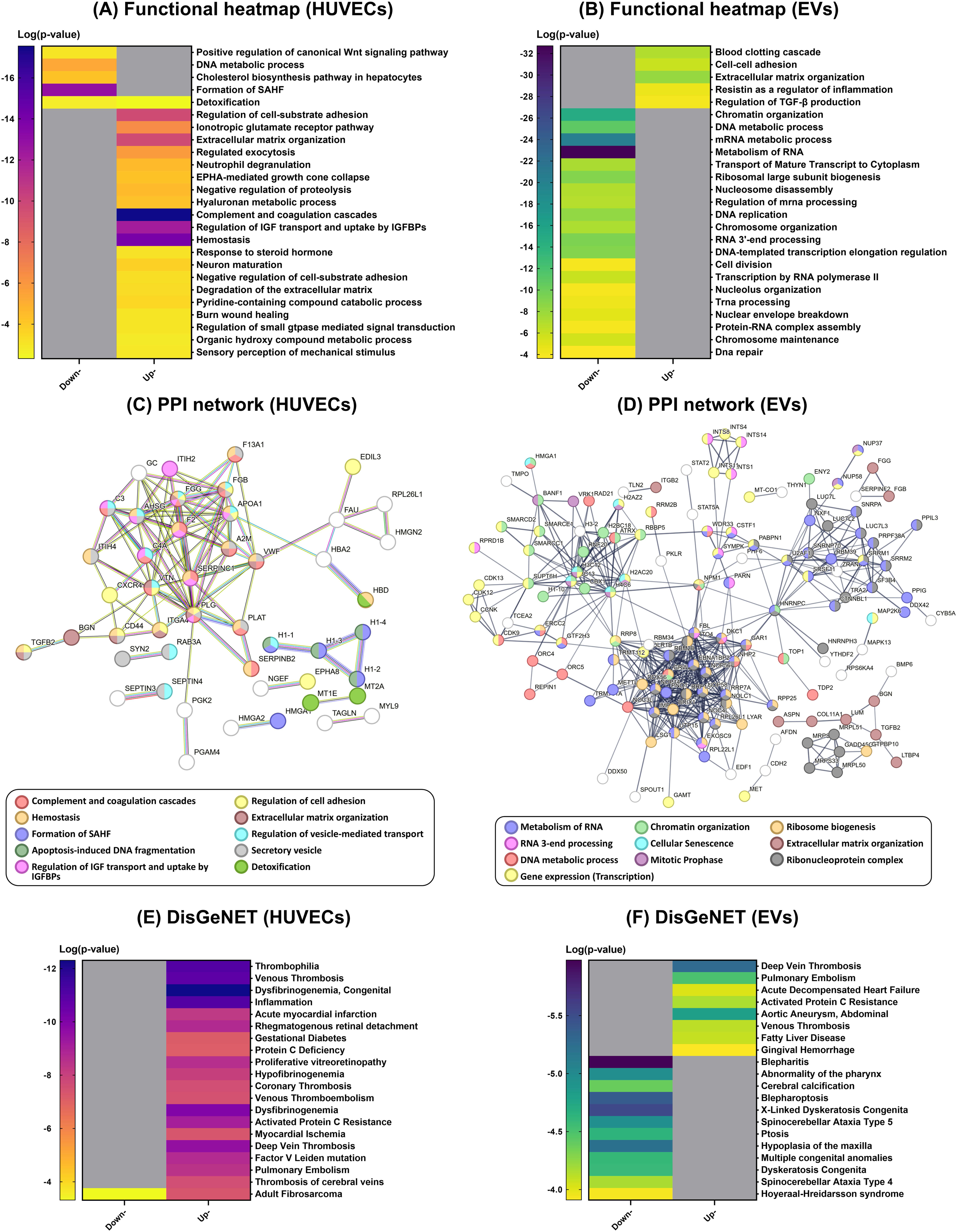
Transcriptomic profiling of senescent and early-passage endothelial cells and their extracellular vesicles (EVs) in a replicative senescence (RES) model. (A) Schematic workflow used for HUVECs and EV isolation and transcriptomics analysis using the NovaSeq X Plus System Illumina platform. (B-C) Volcano plot showing differentially expressed genes in senescent vs. early-passage (B) HUVECs and (C) EVs. Significantly regulated genes in senescent cells (|log2 ratio|≥1, q-value<0.05) and EVs ((|log2 ratio|≥1, q-value<0.25) are highlighted, with log2 ratio≥1 indicating overexpressed proteins and log2 ratio≤–1 indicating underexpressed proteins. Non-significant proteins are shown in gray. (D-E) Principal component analysis (PCA) of DEGs showing clear discrimination between senescent and early-passage (D) HUVECs and (E) EV profiles.

Functional enrichment analysis (Metascape, ORA, and GSEA) was used to assign biological roles and pathways (Figure 5, Supplementary Table 5, and Figures S8). Consistent with the proteomic results, the overexpressed genes in senescent cells were enriched in blood coagulation, platelet activation, and ECM organization (Clusters 1-2, Figure 5B). Furthermore, both DEGs and DEPs in the RES model were associated with vascular processes, such as blood vessel diameter maintenance and regulation of blood circulation (Supplementary Figure S8c). Downregulated genes were primarily associated with the cell cycle, inflammatory responses, and small-molecule biosynthesis, as well as Wnt signaling, DNA replication and repair, and mitosis (Clusters 3-4, Figure 5C). Comparison with senescent EVs further revealed that several downregulated genes and proteins converged on translation- and transcription-related processes, such as mRNA splicing, nucleocytoplasmic transport, and transcriptional regulation (Supplementary Figure S8c). Interestingly, a substantial number of DEGs encoded transcription factors (TFs), including those with zinc-coordinating, helix–turn–helix, and basic DNA-binding domains (Supplementary Table 5).

**Figure 5.**
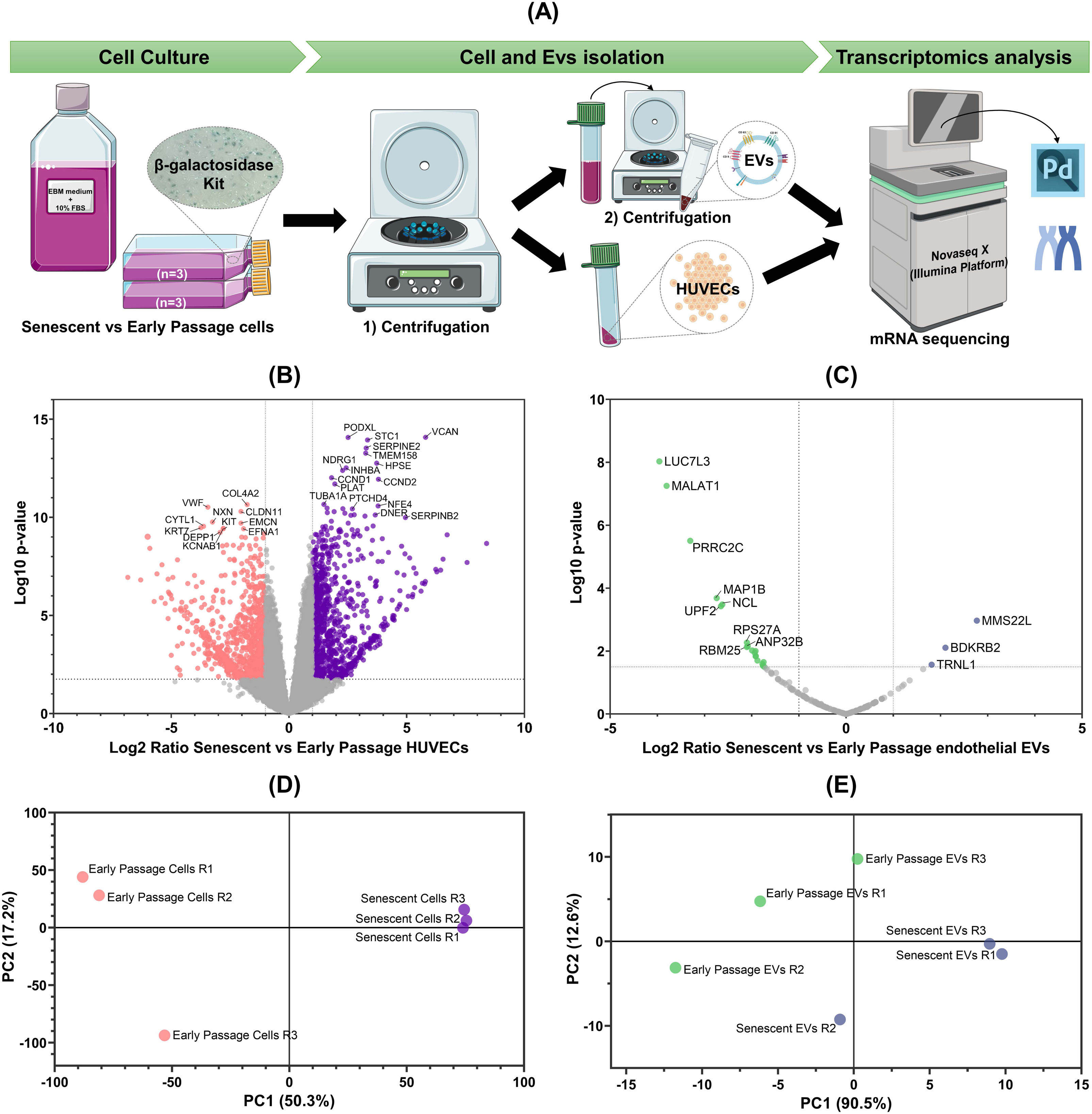
Functional enrichment and protein–protein interaction (PPI) networks of differentially expressed genes (DEGs) in senescent compared to early-passage endothelial cells. (A) Heatmap of functional enrichment analysis of DEGs in endothelial cells based on Gene Ontology (GO), KEGG, and Reactome pathways using Metascape. (B-C) Functional integration of proteomic and transcriptomic datasets using over-representation analysis (ORA). (B) Functional reduction heatmap of upregulated mRNAs and proteins in senescent cells compared to early passage cells. Cluster 1 is related to blood coagulation, wound healing, and platelet activation, while Cluster 2 is mainly associated with extracellular matrix organization. (C) Functional reduction heatmap of downregulated mRNAs and proteins in senescent HUVECs compared to early passage cells. Cluster 3 corresponds to processes related to DNA replication and repair, canonical Wnt signaling, and response to toxic substances, whereas cluster 4 represents cell cycle processes, including mitotic chromosome condensation and metaphase plate congression. (D-E) PPI network of DEGs associated with migration, proliferation, and adhesion generated with STRING (confidence 0.70). Node colors correspond to enriched GO Biological Process terms shown in (E) functional dot plot. Functional dot plots based on KEGG and Reactome pathways are shown below, with dot colors indicating the false discovery rate (FDR) and bubble size indicating the gene count. (F-G) PPI networks of DEGs associated with the cell cycle, p53 signaling, and senescence were generated with STRING (confidence 0.70). Node colors correspond to the enriched functional terms shown in (G) the dot plot. (H) Gene Set Enrichment Analysis (GSEA) plots showing hallmark gene sets enriched in early-passage cells.

In addition, STRING was used to generate protein–protein interaction (PPI) networks, revealing two PPI networks linking DEGs to multiple pathways (Figures 5D/F). The first PPI network was enriched in enzyme-linked receptor signaling and processes regulating migration, proliferation, and adhesion, as well as key pathways including phosphoinositide-3-kinase (PI3K)/Akt, MAPK, Rap1, and AGE-RAGE signaling (Figures 5D-E). Within this PPI network, several DEGs were associated with neovascularization and vascular endothelial growth factor (VEGF) signaling, including TGF-β2, Platelet-derived growth factor (PDGF), Fibroblast growth factor (FGF), angiopoietins (ANGPT), and receptors for these growth factors, including FGFR1. The second PPI network (Figure 5F) was enriched for p53 signaling, cell senescence, cell cycle regulation, DNA damage response (DDR), and Wnt signaling pathways (Figure 5G). Both networks were regulated by p53 and STAT3, which were found to be downregulated and upregulated, respectively, in senescent cells, along with other putative upstream regulators identified by TRRUST, such as E2F1, RELA, NF-κB, SMAD3, and HDAC1 (Figures 5D/F, Supplementary Table 5). In addition, hallmark gene sets, including E2F and MYC targets and G2/M checkpoint genes, were enriched in early-passage cells (FDR<25%), reflecting active cell cycle progression (Supplementary Table 5 and Figures 5I). Their specific association with early-passage cells suggests repression in late passages, which is consistent with senescence-associated cell cycle arrest. Notably, transcriptional repressors, such as E2F7, were upregulated in senescent cells, reinforcing the role of E2F family members in RES.

In contrast, senescent EVs were enriched in fewer processes, including RNA metabolism, nonsense-mediated decay, regeneration, and vesicle-mediated transport (Supplementary Table 6 and Figure 8b). tRNA-Leu (TRNL1), which is upregulated in senescent cells, encodes a tRNA ligase linked to mitochondrial diseases and CVD (DisGeNET). We also detected alterations in the VEGF signaling pathway. Specifically, the bradykinin receptor B2 (BDKRB2) was upregulated in senescent EVs, whereas nucleolin (NCL) was downregulated in both senescent cells and their EVs.

### 2.4. Molecular signature of endothelial replicative senescence through integrated proteomic and transcriptomic profiling of HUVECs and EVs

Establishing the molecular profile of HUVECs during RES and of their EV cargo is critical for understanding endothelial aging and senescence and its association with multiple diseases. In addition to elucidating RES mechanisms, characterizing senescent EVs could provide new insights into their role in secondary senescence and identify novel targets for senotherapeutic intervention.^13,14^ Nevertheless, this process could be inherently complex, since EV cargo can differ markedly from their parent cells at qualitative and quantitative levels, as previously reported.^29^ Accordingly, we performed an integrative analysis of HUVEC and EV datasets, which provided complementary information and specific compartment biomarkers and specific compartment biomarkers.

By comparing these datasets, we identified nine proteins and three mRNAs that overlapped between HUVECs and their EVs (Figure 6A/B and Supplementary Table 7). TGF-β2 and adipogenesis regulatory factor (ADIRF) were exclusively detected, at both the protein and gene levels, in senescent cells and at the protein level in senescent EVs. In contrast, biglycan (BGN) was restricted to early-passage cells, observed at both the protein and gene levels, whereas its downregulation in EVs was evident only at the protein level. Fibrinogen chains were upregulated in both senescent cells and EVs, whereas high mobility group proteins (HMGs) were consistently downregulated. MT2A was the only protein whose expression levels decreased in senescent cells but increased in EVs derived from these cells (Figure 6A). At the transcript level, TRNL1 was upregulated in both cells and EVs, whereas microtubule-associated protein 1B and metastasis-associated lung adenocarcinoma transcript 1 (MALAT1), a well-known long non-coding RNA (lncRNA) that regulates phenotypic switching in endothelial cells^45^, were upregulated in cells but downregulated in EVs (Figure 6B).

**Figure 6.**
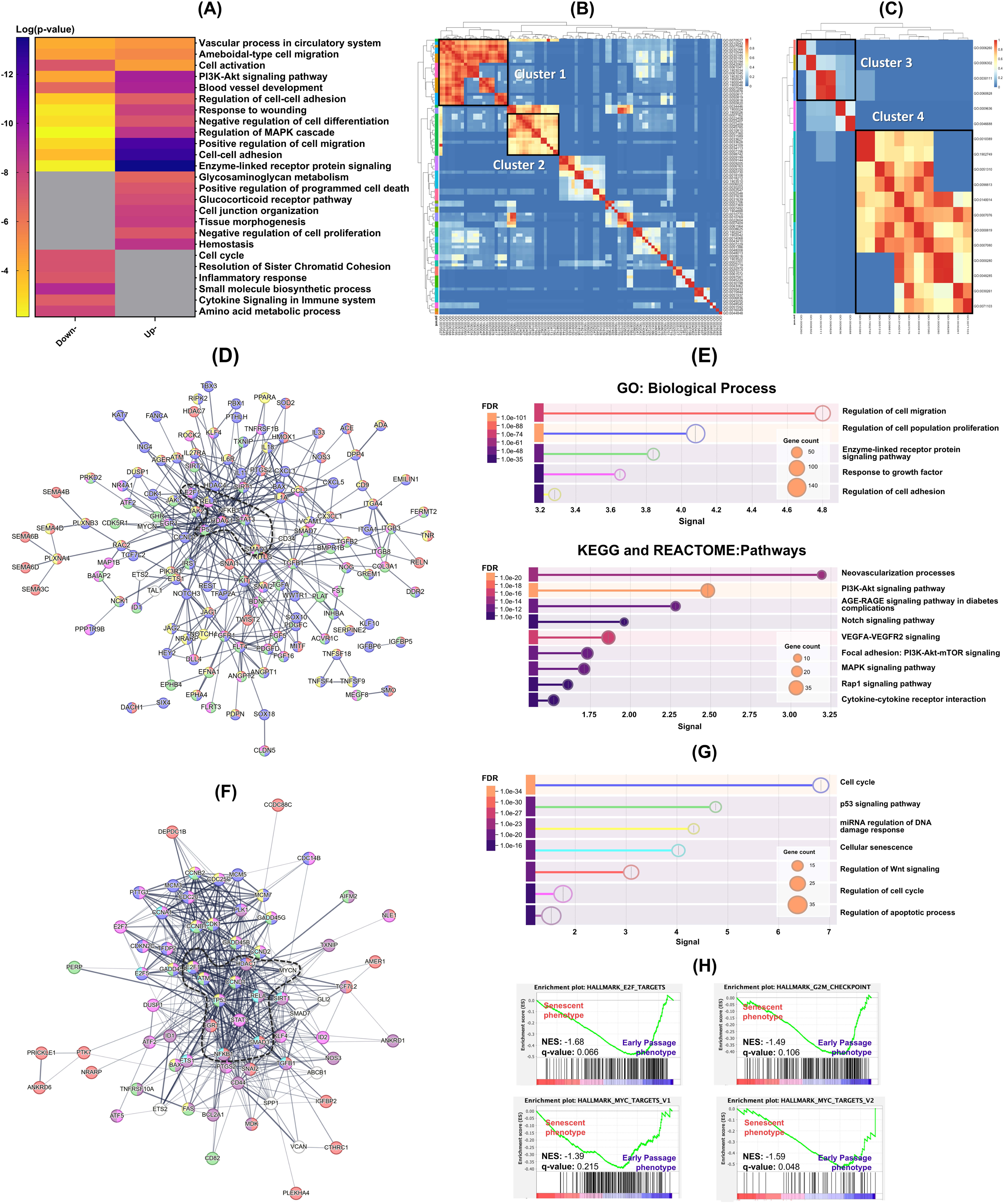
Correlation of transcriptomic and proteomic datasets in replicative endothelial senescence. (A–B) Venn diagrams comparing differentially expressed (A) proteins and (B) genes in senescent and early-passage HUVECs and their extracellular vesicles (EVs). (C–D) Venn diagrams showing the overlap between proteomic and transcriptomic datasets in (C) HUVECs and (D) endothelial EVs. (E) Distribution of correlation coefficients between transcriptomic and proteomic data in HUVECs (left, purple) and EVs (right, green). (F-G) Global mRNA–protein correlation matrix in (F) senescent HUVECs and (G) EVs. Positive correlations are shown in orange and negative correlations are shown in blue.

To establish a molecular signature of RES at both the gene and protein levels, we integrated proteomic and transcriptomic data from cells and EVs (Figure 6C-G and Supplementary Table 7). Although several authors have reported a lack of correlation between mRNA and protein levels,^46^ a good positive correlation was observed in this study, despite the low degree of overlap, as shown by the Spearman’s rank correlation coefficient (Rho) (Figure 6C-E).

In the cells, 31 genes/proteins overlapped between the two datasets (Figure 6C). Among them, 28 genes were positively correlated with protein levels; 18 were up-regulated and 10 were down-regulated in senescent cells. Interestingly, heat shock 70 kDa protein 12B and CD44 were consistently upregulated across both datasets (Figure 6F). Down-regulated biomolecules, including BGN, von Willebrand factor, Protein MGARP, and Inactive tyrosine-protein kinase 7 (PTK7), displayed very high correlations (rho>0.9), suggesting their potential as RES biomarkers at both the gene and protein levels.

Chronophin, C-type lectin domain family 11 member A, and insulin-like growth factor-binding protein 2 (IGFBP2) were the only overlapping molecules that showed no correlation between the transcriptomic and proteomic datasets. IGFBP2 was detected only at the protein level in senescent cells, whereas its mRNA was downregulated, unlike other IGF-binding proteins (IGFBPs). Other DEPs, although not significantly regulated at the gene level, tended to increase or decrease, displaying a high correlation coefficient (rho>0.9) (Figure 6F and Supplementary Table 7). In senescent EVs, Luc7-like protein 3 (LUC7L3) showed a coordinated decrease at both the transcript and protein levels (Figure 6G), whereas these changes were not observed in cells, suggesting that LUC7L3 is a potential molecular biomarker of endothelial secondary senescence.

Finally, comparison with high-throughput transcriptomic senescence signatures, including CellAge,^47^ SeneQuest,^48^ and SenMayo^49^ revealed a substantial overlap with our dataset in both senescent HUVECs and their respective EVs (Figure 7 and Supplementary Table 7). Many of the molecules described above were also present in these datasets, reinforcing their potential as biomarkers of RES.

**Figure 7.**
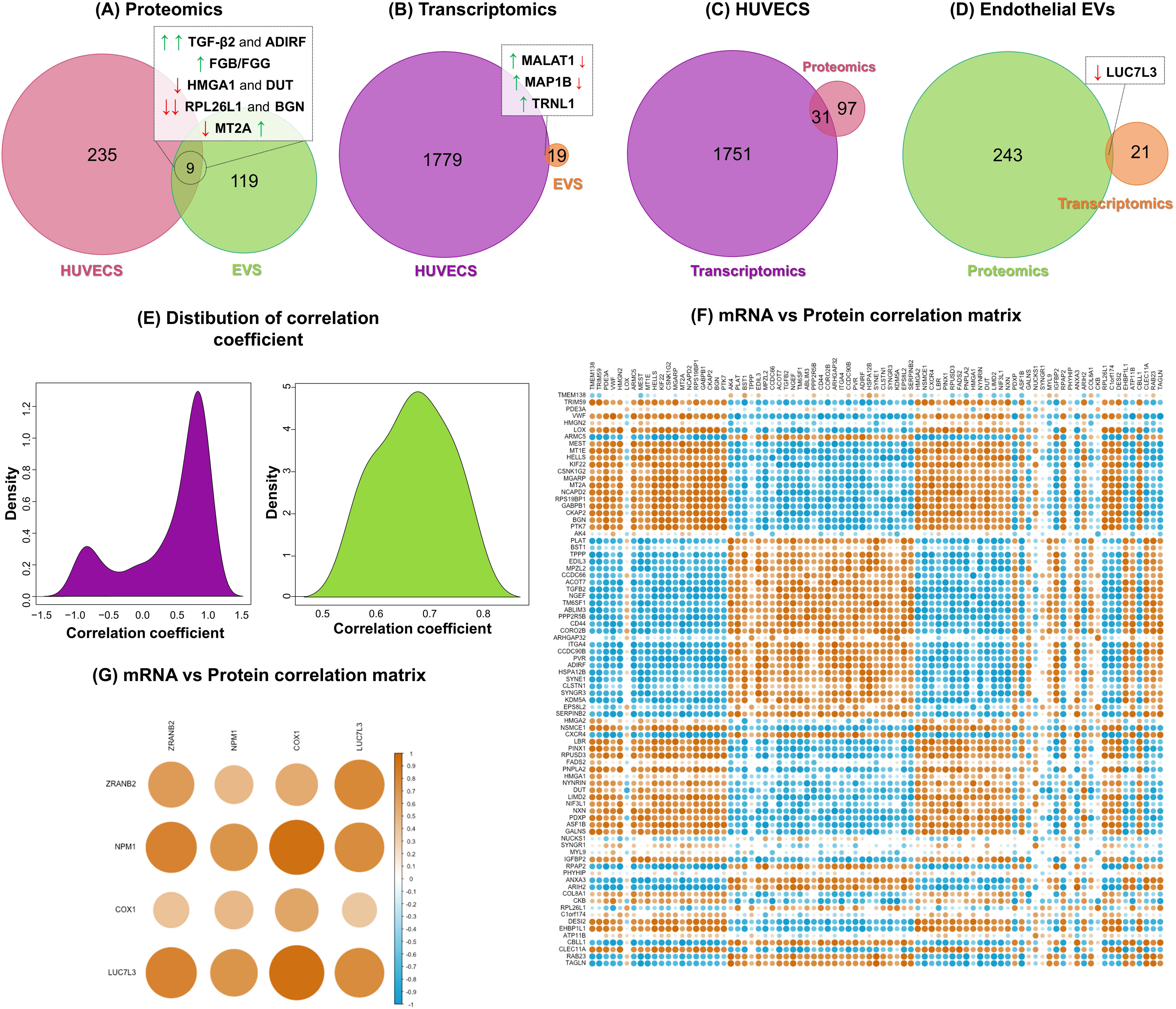
Venn diagrams comparing differentially expressed proteins/genes in (A) senescent HUVECs and their (B) extracellular vesicles with high-throughput transcriptomic signature gene sets, including CellAge^47^, SeneQuest^48^, and SenMayo^49^.

### 2.5. miR-22-3p and miR-126-5p are putative regulators of replicative endothelial senescence in HUVECs and secreted EVs

Finally, to assess whether post-transcriptional mechanisms contribute to RES, we conducted a miRNA profiling of senescent and early-passage HUVECs, as well as their respective EVs (Supplementary Table 8).

In cells, 140 miRNAs were found to be differentially regulated (|log2FC|>1; q-value≤0.05), with 91 upregulated and 49 downregulated miRNAs (Figure 8A). In contrast, EVs showed low transcript yield, with only 10 differentially expressed miRNAs, of which 4 were upregulated and 6 were downregulated (|log2FC|≥1, q-value≤0.25) (Figure 8B). Of the EV-associated miRNAs, only miR-22-3p and 126-5p were also found differentially expressed in the senescent cells (Figure 8C and Supplementary Table 8). While miR-22-3p was found to be upregulated in senescent cells but downregulated in EVs, miR-126-5p was found to be downregulated in both senescent cells and EVs, consistent with our previous results.^26^ In contrast, miR-29a-3p, miR-22-3p, and miR-4488 were exclusively deregulated in senescent EVs (Figure 8C/D).

Functional analysis of miRNAs differentially expressed in cells and EVs revealed an enrichment in angiogenesis, EC migration and proliferation, and vascular development (Figure 8E/F and Supplementary Table 8 and Figure S9/S10). In EVs, miR-22-3p and miR-126-5p, together with EV-upregulated miR-15b-5p and miR-29a-3p, have also been implicated in the regulation of these processes. While VEGF, a central angiogenic mediator^50^, was not significantly downregulated at the transcript level (showing only a tendency), several miRNAs involved in the regulation of VEGF production, hypoxia response, and cellular response to VEGF were found to be differentially expressed (Figure 8F and Supplementary Table 8). Most of these miRNAs were downregulated, with some exceptions in senescent cells (e.g., miR-34a-5p) and EVs (e.g., miR-15b-5p).

Other biological processes regulated by differential miRNAs in both cells and EVs were also consistent with a senescent phenotype, including the regulation of the cell cycle, translation, cell-death pathways, and cytokine production (Figure 8F/G). Indeed, both upregulated (miR-22 and miR-34a-5p) and down-regulated miRNAs (e.g., miR-21-5p, miR-590, miR-17, and miR-20) in HUVECs are regulators of cellular senescence, including replicative senescence. Similarly, most of these miRNAs were correlated with cell aging and its regulation. Interestingly, upregulated (miR-22 and miR-34a-5p) and downregulated miRNAs (miR-17 and miR-590) were associated with positive and negative regulation of cellular aging, respectively. Furthermore, differentially regulated miRNAs stand out as regulators of key pathways of the NF-κB signaling (e.g., miR-590, miR-15b, miR-21), TGF-β signaling (e.g., miR-15b, miR-21), PI3K signaling (miR-20a-5p, miR-126, and miR-21), and the intrinsic apoptotic signaling (e.g., miR-21-5p).

### 2.6. miRNAs correlate with proteomics and transcriptomics changes

To study the influence of miRNAs on RES, we performed a comprehensive molecular integration of miRNA, transcriptomic, and proteomic data from senescent HUVECs. These analyses revealed significant correlations, identifying 75 differentially regulated miRNAs in senescent cells that correlated with 149 genes and 6 proteins. Among these, 29 miRNAs were associated with a single gene/protein, but most miRNAs exhibited multiple interactions, forming regulatory clusters of interconnected miRNAs and genes/proteins (Figure 9). These findings suggest the existence of a complex regulatory network that drives the biological processes and pathways of RES. Curiously, according to disease knowledge databases, many of the regulated molecules have been associated with several types of cancer, including breast, skin, liver, and kidney (Supplementary Figure 11).

**Figure 8.**
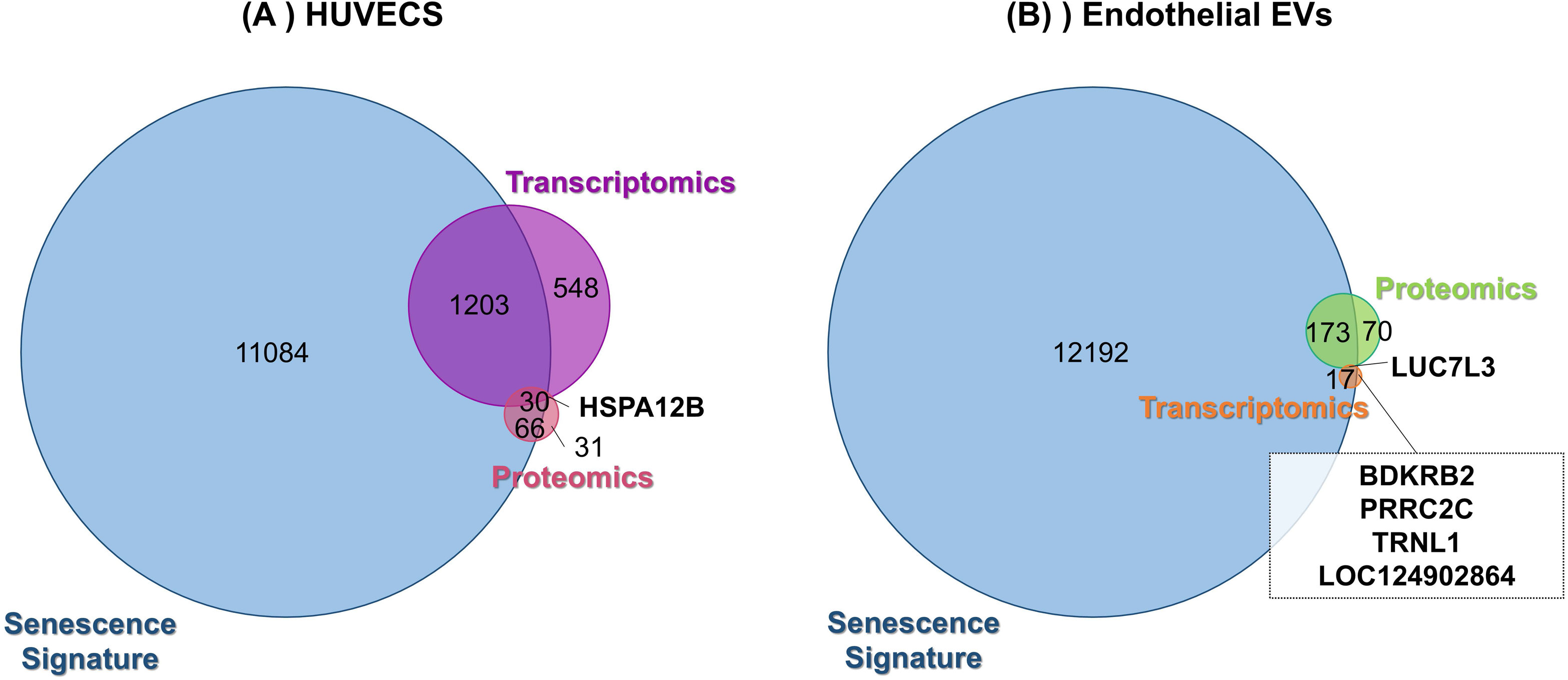
miRNA profiling of senescent and early-passage endothelial cells and their extracellular vesicles (EVs) in a replicative senescence (RES) model. (A-B) Volcano plot showing differentially expressed genes in senescent vs early-passage (A) HUVECs and (B) EVs. Significantly regulated genes in senescent cells (|log2 ratio|≥1, q-value< 0.05) and EVs ((|log2 ratio|≥1, q-value<0.25) are highlighted, with log2 ratio≥1 indicating overexpressed proteins and log2 ratio≤–1 indicating underexpressed proteins. Non-significant proteins are shown in gray. (C) Venn diagram showing the overlap of differentially expressed miRNAs between senescent HUVECs and EVs. (D) Heatmap of miRNAs found to be differentially expressed in senescent compared to early-passage EVs. (F-G) Functional enrichment of predicted targets of differentially expressed miRNAs in senescent (F) HUVECs and (G) EVs. Bubble plots display enriched biological processes and pathways (upregulated miRNA above and downregulated below), with the bubble size representing the number of genes and color indicating different functional enrichment terms according to gene ontology.

**Figure 9.**
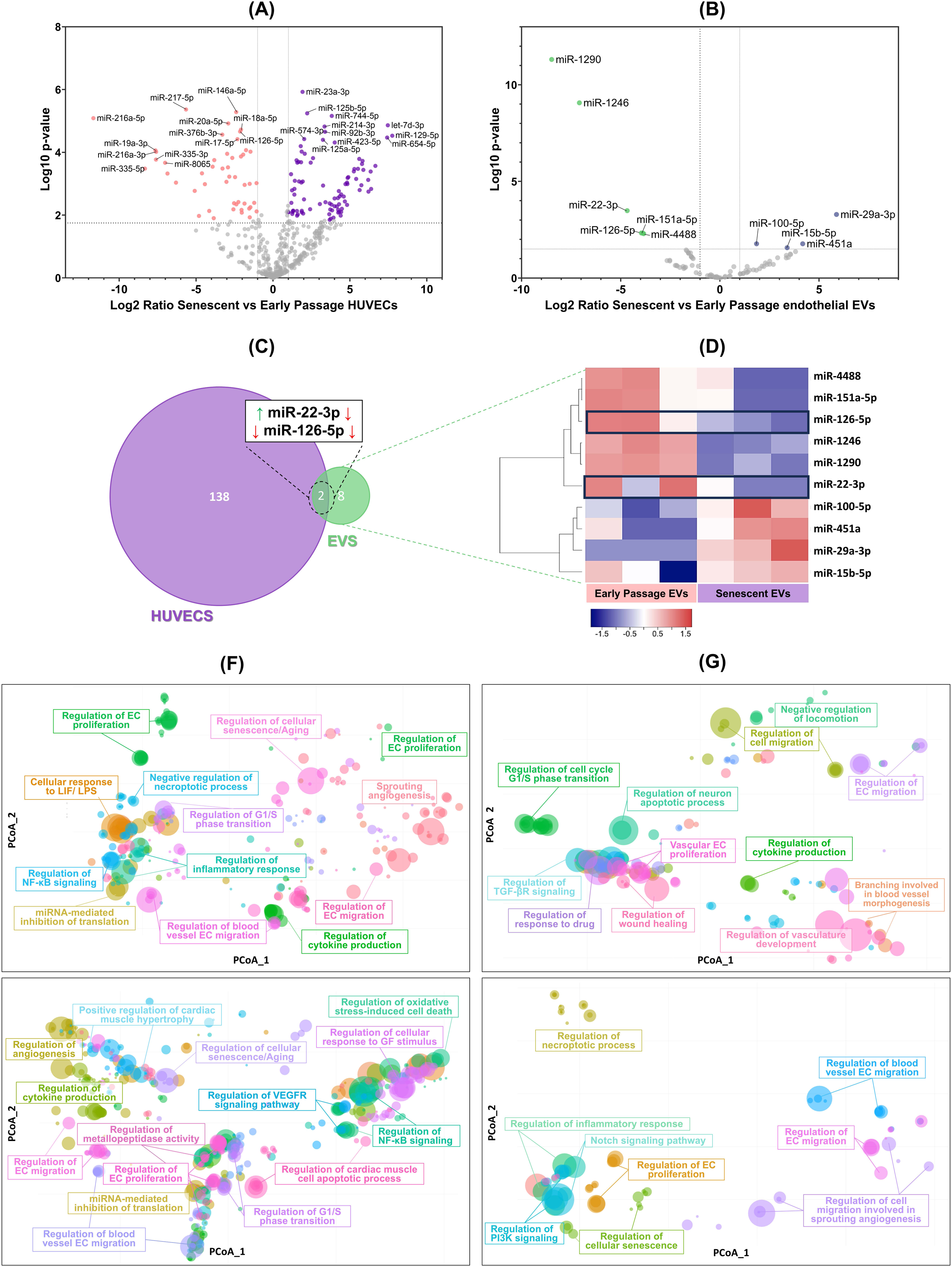
miRNA–target correlation networks in replicative endothelial senescence (RES). (A–D) Clusters showing significant correlations between differentially expressed miRNAs and their putative targets (genes and proteins) in senescent HUVECs. A force-directed (OpenCL) layout was used to generate networks in Cytoscape (Metscape) by integrating miRNA, transcriptomic, and proteomic data with the KEGG database. Edges denote significant miRNA–gene/protein correlations based on p-value and node shapes represent miRNAs (purple polygons), genes (squares), and proteins (squares). Nodes are color-coded according to functional categories, including regulation of cell migration/motility, angiogenesis, PI3K–Akt signaling, focal adhesion, extracellular matrix remodeling, metabolism and transcriptional regulation.

Cluster A (Figure 9A) emerged as a significant molecular network, comprising 21 miRNAs downregulated in senescent cells that correlated with 78 genes and a protein, transmembrane 6 superfamily member 1 (TM6SF1). These miRNAs included miR-335-3p, which held a central position in the cluster and was correlated with more than 20 genes (Supplementary Table 9). Other miRNAs include miR-590, miR-17 family (miR-17-5p, -19, -20a, -106a), and miR-29 family. Functional analysis of miRNA targets indicated that they play roles in cell motility and migration, focal adhesion, transmembrane transport, angiogenesis, and ECM/cytoskeleton organization (Figure 9 and Supplementary Table 9). In cluster A (Figure 9A), a regulatory network revealed a PI3K-Akt–centered cluster involving insulin signaling (e.g., Insulin receptor substrate 1), angiogenic mediators (ANGPT1 and FGFR1), G1/S transition regulators (CCND1/2), integrins (ITGA4), and ECM proteins (Reelin, COL4A1). Senescent cells also exhibited reduced levels of PI3K subunits (PIK3CG, PIK3R1) and Akt-regulating miRNAs (miR-126, miR-21, miR-20a). In addition, KIT, a key regulator of EC survival and angiogenesis, was potentially downregulated by miR-221-3 (correlation coefficient=-1) (Supplementary Table 9). Notably, molecular analysis has identified both pro-angiogenic/proliferative and anti-angiogenic/anti-proliferative miRNAs as potential regulators of senescence, influencing both the promoters and inhibitors of these processes. For instance, miR-590-3p has been correlated with both angiogenic proteins (ANGPT1) and negative regulators of VEGF-induced angiogenesis, including Rho GTPase-activating protein 42 (ARHGAP42) and Rho-associated protein kinase 2 (ROCK2) (Figure 9A and Supplementary Table 9).

These results also emphasize that several miRNAs can regulate critical TFs, including DNA-binding protein inhibitors, zinc finger (ZNF), Krüppel-like factors (KLF), and homeobox genes (HOXC8 and SIX4) (Figure 7). These TFs were regulated by various miRNAs in clusters A, B, and C (Figure 7). In cluster A, RE1-Silencing TF (REST), which is upregulated in senescent cells, was correlated with miR-29b-3p, miR-106a, and miR-335-3p. In clusters B and C, miR-23 isoforms correlated with several TFs (ZNF493, ZNF626, and ZNF506), whereas miRNA-23a-3p correlated with high mobility group box 2 (HMGB2) and Metallothionein 2A (MT2A), both of which are associated with senescence (Figure 9B/C).

Finally, the same comprehensive molecular integration of miRNA, transcriptomic, and proteomic data was performed in EV studies. In contrast to the cells, only two miRNAs, miR-29a-3p and miR-126-5p, showed significant correlations with Glutathione Peroxidase 7 (GPX7) and Neurocan (NCAN) in senescent EVs (Supplementary Table 9). GPX7 was detected only in early-passage EVs, whereas NCAN was detected only in senescent EVs.

## 3. Discussion

Endothelial senescence has emerged as a promising therapeutic target for CVD, highlighting the importance of elucidating the molecular mechanisms underlying vascular aging and dysfunction.^23^ In this study, an integrated transcriptomic and proteomic multi-omics approach was applied to a RES model based on the Hayflick and Moorhead model^8^, which mirrors key aspects of vascular aging such as impaired wound healing, endothelial dysfunction, enlarged morphology, and elevated SA-β-Gal activity, as previously described.^26^ Our main multi-omics findings revealed a canonical senescence signature in this RES model in HUVECs and their EVs, marked by cell cycle arrest, DDR, NF-κB signaling, and SAPS induction, as well as a pronounced suppression of RNA metabolism and translation, and ribosome biogenesis. We also identified consistent transcript-protein changes in senescent cells and EVs, mechanistically and functionally linked to senescence, vascular deterioration, and CVD, revealing EC-specific senescence markers (such as ADIRF, TGF-β2, CD44, IGFBPs, BGN, ITGA4, and MT2A) and post-transcriptional regulators, including LUC7L3, MALAT1, and miRNAs (e.g., miR-22-3p, miR-126-5p). Importantly, several mediators of endothelial dysfunction, including angiogenic factors and ECM components, were identified in our omics experiments, reinforcing the importance of endothelial senescence in CVD development.

Multi-omics profiling confirmed a canonical senescence signature in ECs and EVs, revealing a strong enrichment of the cell cycle, DDR, and SAPS, which overlapped with reference gene sets such as those described in CellAge^47^, SeneQuest^48^, and SenMayo^49^. Late-passage ECs and their EVs displayed SASP enrichment in acute phase proteins, coagulation and complement factors, pro-fibrotic components, adhesion molecules, angiogenic mediators, and ECM-remodeling proteins, which are strongly linked to CVD, vascular dysfunction, fibrotic, and coagulation disorders (DisGeNET). The presence of these molecules in EVs is particularly relevant, as several studies have shown that EV-mediated secretion of SASP components promotes secondary senescence.^16^ Our data also support this hypothesis, as several molecules implicated in NF-κB signaling were exclusively detected in senescent EVs, including NF-κB essential modulator (NEMO) and FAS (TNFRSF6). NEMO is a key upstream activator of NF-κB signaling, a central driver of the SASP and chronic inflammation linked to age-related diseases and CVD. ^51,52^ Both NEMO and FAS act as key mediators of stress-induced signaling in the senescent ECs. While NEMO mediates IL-1 and TNF receptor signaling and NF-κB activation in response to DNA damage,^51^ FAS has been found to increase the susceptibility of senescent ECs to TNF-α-induced apoptosis.^53^

The hallmarks of aging encompass genomic instability, telomere attrition, and epigenetic alterations^54^, as well as the recently emerged dysregulation of RNA processing and metabolism^55^ and the transcriptional repression of DNA repair genes^56^. These age-related changes are associated with the loss of protective factors that predispose cells to replication stress, persistent DDR, and senescence, thereby disrupting stress-responsive gene expression, which is essential for maintaining cellular adaptability under adverse conditions^55^. Our analysis of the senescent EV profile revealed a pronounced suppression of RNA metabolism, translation, chromatin organization, and ribosome biogenesis. Accordingly, several transcriptional cyclin-dependent kinases (CDK9, CDK12, and CDK13), essential for splicing, RNA processing, genome stability, and DDR,^57,58^ together with DNA repair regulators (e.g., RAD21, ERCC2), were exclusively detected in EVs from early-passage cells. In parallel, senescent HUVECs showed coordinated repression of DNA-repair machinery, including Fanconi proteins, damage-inducible factors, chromatin regulators, and altered E2F family levels, which have been shown to induce secondary DNA damage and premature senescence.^56^ We also observed a concerted down-regulation of histones, histone chaperones, replication-licensing factors, and HMGBs in senescent cells and their EVs, indicating chromatin remodeling that leads to the inhibition of transcriptional activity. In addition, senescent EVs show downregulation of protective mechanisms, particularly components of the nonsense-mediated decay pathway, a well-characterized RNA surveillance mechanism that prevents the accumulation of aberrant transcripts, which is inhibited in senescence^59^ and cardiomyopathy^60^. Collectively, these findings suggest that early passage EVs can propagate repair-supporting signals, whereas senescent EVs lose these reparative properties and export fewer repair and RNA-processing molecules. Further supporting this hypothesis, our results indicated that senescent EVs also lack key transcriptional regulators, such as ATRX and splicing factors, such as LUC7L3, which are critical for DDR, heterochromatin maintenance, and genome stability^61,62^, suggesting that they may propagate genome instability. Indeed, ATRX downregulation has also been observed in uremia-induced endothelial dysfunction and senescence^29^.

Impairment of endothelial function is a hallmark of endothelial senescence, a central driver of vascular aging and CVD, through loss of vasoprotective capacity and a shift toward a pro-inflammatory and pro-thrombotic phenotype.^24,63,64^ Multi-omics profiling revealed consistent transcript–protein changes in senescent cells, many of which are mirrored by EVs, identifying potential biomarkers mechanistically linked to senescence, endothelial dysfunction, and/or CVD. For instance, both senescent cells and their EVs are enriched in adipogenic mediators such as ADIRF, a key regulator of adipogenesis.^65^ Senescent EVs also contain RETN, a pro-inflammatory adipokine linked to insulin resistance, obesity, and CVD.^66–69^ Higher RETN levels predict CVD events in older adults, independently of traditional risk factors^69^ and ischemic stroke risk in postmenopausal women,^68^ emphasizing its role in both CVD and aging. Consistent with data on uremic-induced ECs senescence, in which EV fatty acid metabolism changes co-occur with cellular adipogenesis and inflammation,^29^ these findings reinforce senescent EVs as vectors of adipogenic and inflammatory mediators that amplify endothelial dysfunction, promote atherogenesis, and increase cardiovascular risk. Nevertheless, senescence-driven endothelial dysfunction in this RES model converged on two major pathological axes: maladaptive angiogenesis and ECM remodeling.

Endothelial barrier breakdown is an early hallmark of vascular aging and a key driver of CVDs, such as hypertension and atherosclerosis.^70^ Barrier integrity relies on tight junction (TJ) proteins; however, this delicate balance can be compromised by numerous stimuli, including SASP-derived pro-inflammatory cytokines, growth factors, and ECM-remodeling enzymes, increasing permeability, leukocyte recruitment, and angiogenesis.^70,71^ In our study, we have observed a loss of claudins, particularly the endothelium-specific TJ protein Cldn5, in both senescent ECs and EVs, suggesting not only a barrier compromise but also their potential propagation via secondary senescence. The compromised endothelial barrier can trigger angiogenesis, a process essential for EC proliferation and wound healing, consisting of the sprouting of new capillaries. Angiogenesis declines with aging and has been associated with ECs senescence ^26^, contributing to increased CVD risk^72^. Increasing pieces of evidence suggest that age-related alterations in the vascular ECM also represent one of the mechanisms of impaired angiogenesis.^72^ Our data support a senescence-driven rewiring of angiogenic programs in EC senescent cells, as well as changes in ECM remodeling.

Although VEGF levels remained unchanged in our model, senescent ECs displayed widespread repression of angiogenic modulators, including Mgarp^73^, PTK7, and multiple growth factors^50,74^ (PDGF and FGF), together with key pro-angiogenic miRNAs (miR-17 family, miR-216a-3p, miR-29)^75,76^ and repressors (e.g., miR-21-5p^77^). IGF signaling, another central pathway in endothelial physiology and angiogenesis regulation^78,79^, was also found to be enriched in the present study. Several IGFBPs, including IGFBP5 and IGFBP7, consistent with their roles as SASP components and paracrine mediators,^40,80–83^ were overexpressed in senescent ECs and in EVs. In parallel, miR-320a-3p, a proposed cardiomyopathy biomarker that mediates anti-angiogenic effects by IGF1 targeting in cardiomyocytes,^84,85^ was upregulated in senescent ECs. The upregulation of pappalysins (PAPPA and PAPPA2), proteases that release IGFs and modulate IGFBP–ECM interactions^78,84^, further implicates IGF signaling in senescence-related angiogenic dysfunction and ECM remodeling.

Changes in angiogenic modulators have also been observed in EVs, supporting the hypothesis that angiogenic rewiring is mediated by EV-mediated signaling. EVs from senescent cells transport higher levels of BDKRB2 and angiogenesis-related miRNAs (miR-15b-5p and miR-29a-3p). These miRNAs have emerged as attractive therapeutic targets, as previous studies have linked them to aging and ischemic cardiomyopathy through their role in the VEGF-modulatory network^86^. In contrast, BDKRB2 mediates bradykinin signaling, a key vascular homeostasis pathway that regulates vasodilation, vascular permeability, and inflammation. Its clinical relevance is underscored by the widespread use of bradykinin-pathway modulators in CVD therapy.^87,88^ Conversely, senescent cells export lower levels of NCL, a multifunctional protein that acts as a co-receptor for several growth factors and a validated anti-angiogenic target^89^, consistent with impaired angiogenesis and defective vascular repair. Nevertheless, since NCL is also a multifunctional regulator of rRNA maturation, ribosome assembly, and mRNA metabolism,^90^ its loss may reflect coupled defects in RNA metabolism and angiogenic signaling. In senescent EVs, we also observed lower levels of miR-22-3p and lncRNA MALAT1, both of which displayed divergent regulation between senescent cells and their EVs, indicating compartment-specific roles in the regulation of angiogenesis. MALAT1 is a member of the “Angio-LncRs”, a group that regulates pro-angiogenic transcriptional programs and modulates endothelial proliferation and senescence.^24,45,91^ In its turn, miR-22-3p can contribute to CVD, such as myocardial infarction, hypertrophy, and heart failure, largely by promoting fibroblast migration, hypertrophy, and suppression of cardiac autophagy.^92–94^ This miRNA has been linked to senescence, inflammation, and angiogenesis; however, while previous studies reported miR-22-3p induction with aging and its enrichment in EVs from aged mice,^11,93,95,96^ our data show that senescent EVs transport lower levels of miR-22-3p. Therapeutically, miR-22 inhibition has been shown to restore autophagy and improve post-infarction cardiac remodeling and function in aged mice,^97^ while MALAT1 silencing enhances endothelial cell migration and reduces angiogenesis in vivo, suggesting that its inhibition may offer therapeutic benefits in neovascular diseases.^45^ Considering this, the selective downregulation of miR-22-3p and MALAT1 in EVs may represent a protective mechanism to limit the paracrine spread of pro-senescent signals. However, additional studies are required to elucidate their compartment-specific functions in endothelial aging and determine whether their reduced export contributes to the modulation of secondary senescence.

Another common feature of senescent HUVECs is the alteration of cytoskeletal dynamics and ECM remodeling, reinforcing recent findings that matrisome modifications are major drivers of aging.^98^ Under physiological conditions, ECM turnover is tightly regulated, maintaining vascular structure and function, but this balance is disrupted with aging and senescence,^99,100^ leading to excessive ECM deposition, fibrosis, and eventually to CVD^101–103^. We have found that several ECM components, such as CD44, ITGA4, TGF-β2, and fibrinolytic system components (tPA and Plasminogen activator inhibitor 2) were consistently upregulated in senescent HUVECs, reflecting a pro-fibrotic phenotype switch. TGF-β2, which is markedly increased in both senescent cells and EVs, is a pleiotropic cytokine that governs vascular remodeling, cellular senescence, endothelial–mesenchymal transition, and fibrosis, which are key pathophysiological features of CVD.^104–106^ TGF-β2 is particularly linked to cardiovascular development and pathological remodeling,^106^ highlighting it as a promising therapeutic target. In contrast, the leucine-rich proteoglycan BGN was uniquely detected in early-passage cells and EVs. BGN is essential for ECM assembly and collagen organization, but it also promotes VEGF-driven angiogenesis.^107,108^ The role of BGN in CVD is complex and context-dependent, as its deficiency can reduce fibrotic scarring and preserve cardiac function,^107^ whereas its activation can conversely protect cardiac cells under stress conditions.^109,110^ This duality underscores the need for tight regulation of BGN expression, positioning it as both a potential biomarker and therapeutic target in RES and CVD.

Our data also revealed a complex coordinated miRNA-TF regulatory network in senescent cells, supporting previous evidence that increased chromatin accessibility is not merely a consequence but also a driver of senescence orchestrated by TFs.^111^ In this context, our correlation analyses suggest that upregulation of specific miRNAs may contribute to the downregulation of key TFs. For example, within clusters B and C, the miR-23 isoforms showed negative correlations with several transcriptional regulators (SIX4, ZNF493, ZNF626, and ZNF506). In particular, ZNF506 is a key regulator of the early dynamic signaling in the DDR pathway and its deficiency has been associated with genomic instability.^112^ Reinforcing the role of miR-23-TF regulation in RES, miR-23a-3p has also been correlated with HMGB2 and MT2A, both of which are associated with senescence. Consistent with our findings, the loss of HMGB2 has been identified in other studies as an early event in replicative senescence, where it plays a crucial role in controlling SAHF formation, SASP regulation, and transcriptional remodeling.^113,114^ In addition, HMGB2 has been previously identified as a direct target of miR-23a-3p in endothelial senescence, suggesting that selective miRNA inhibition could represent a potential senomorphic strategy.^115^ MT2A is a cysteine-rich metallothionein involved in metal-ion homeostasis and antioxidant defense.^116^ It has been proposed as a therapeutic target in inflammatory diseases and cancer^116^, as well as being linked to cardiomyopathy^116^ and longevity^117,118^. Interestingly, although its downregulation in senescent cells appears to be mediated by miR-23a-3p, MT2A was enriched in senescent EVs, indicating compartment-specific regulation. Although their therapeutic potential in vascular aging and CVD remains largely unexplored, further studies are required to clarify the specific roles of MT2A in secondary senescence and its value as a potential target for endothelial dysfunction.

Additionally, another coordinated miRNA–TF correlation network emerged in our analysis, composed of several downregulated miRNAs showing strong negative correlations with multiple TFs, including REST, Krüppel-like factors (KLF11, KLF10), and zinc finger (ZBTB4, ZBTB7A, ZNF480).

Within this network, miR-335-3p appeared to be a central regulatory node, correlating with more than 20 genes, including REST, ZNF480, ZBTB7A, and NPAS3. The role of miR-335-3p in aging and senescence is well established, with previous studies reporting its upregulation in these contexts.^119–121^ However, in our study, miR-335-3p was found to be downregulated and associated with increased TF expression, suggesting a tissue- and context-dependent function, consistent with its dual roles described in cancer.^122,123^ In addition to miR-335-3p, other miRNAs previously linked to senescence,^124–126^ aging,^125,127^, and CVD^75,92,94^, including miR-590-3p and miR-29/miR-17 family members, will be discussed below, were also found to be downregulated and negatively correlated with multiple TFs. These findings suggest that coordinated miRNA downregulation contributes to the derepression of TFs and target genes that modulate essential endothelial processes, thereby promoting endothelial dysfunction, as discussed below.

Consistent with this, several of these downstream targets, found in senescent cells, have been previously implicated with fibrosis, vascular dysfunction, inflammation, and CVD, including angiogenic mediators (e.g, ANGPT1^128^), cytoskeleton-associated genes (e.g., ROCK2),^129,130^ integrins^131^ (e.g., ITGA4), metalloproteinases (e.g., ADAMTS6 and PAPPA), and ECM glycoproteins (e.g., Reelin^132^). Interestingly, many of these genes converged on the PI3K–Akt signaling axis, a central regulator of endothelial survival and angiogenesis.^133^ Given its pivotal role in vascular homeostasis, impaired PI3K–Akt signaling has been linked to atherosclerosis, ischemia/reperfusion injury, hypertrophy, and heart failure, underscoring its clinical relevance.^134,135^ In our study, KIT, an upstream PI3K–Akt effector involved in cardiac repair and survival,^136^ was inversely correlated with miR-221-3p in senescent cells, suggesting that its vascular repair capacity could be impaired by miRNA regulation. Collectively, these observations indicate that a complex miRNA regulatory network underlies the transcriptional and functional alterations observed in our RES model, as discussed below.

One of the miRNAs that correlated with TFs in our analysis was miR-126-5p, a key endothelial regulator of vascular integrity, proliferation, vessel maturation, and angiogenesis.^137,138^ In our study, miR-126-5p was downregulated in senescent HUVECs nd negatively correlated with HOXC8, a conserved homeobox TF involved in cell proliferation, migration, angiogenesis, and epithelial–mesenchymal transition.^139^ Additionally, miR-126-5p was correlated with CDC14B, a protein phosphatase involved in mitotic exit and DDR,^140^ reinforcing its role in the senescence program. This is consistent with our previous findings showing miR-126-5p downregulation in RES and identifying HIF-1α as a miR-126 target involved in protective and reparative endothelial functions, supporting its potential therapeutic benefit in age-related vascular disease.^26^ In this study, we further observed that senescent EVs contained reduced miR-126-5p levels, a pattern consistent with impaired angiogenic and reparative capacity and indicative of EV-driven propagation of endothelial dysfunction. Moreover, miR-126-5p levels in EVs were correlated with NCAN, a hyalectan uniquely detected in the senescent EV population, indicating a potential link between angiogenesis and ECM remodeling during secondary-senescence. Unlike other widely expressed ECM hyalectans (e.g., decorin or versican), NCAN is a specific component of the central nervous system ECM essential for brain development.^141,142^ Although its precise role in RES remains unclear, a previous study showed that HUVECs co-cultured with neural stem cells were the main source of NCAN and other neuropil-associated proteins in vasculature-like structures, underscoring their contribution to the neurovascular microenvironment and angiogenesis-mediated recovery.^142^

Although not directly correlated with TFs in our dataset, miR-590-3p is particularly relevant because it links the fibrotic and angiogenic pathways. Its loss has been associated with vascular dysfunction, whereas EV-mediated delivery has been shown to improve myocardial infarction outcomes.^143^ Furthermore, the role of miR-590-3p as an anti-fibrotic regulator of ECM remodeling makes it a promising therapeutic candidate for restoring cardiac function following myocardial infarction.^144^ Our findings further support a central role for miR-590-3p in endothelial function and ECM remodeling. It has correlated with both ANGPT1, a vascular protective molecule that promotes vessel maturation and stability and EC survival,^128^ and negative regulators of VEGF-induced angiogenesis, including ARHGAP42, a regulator of vascular tone associated with hypertension risk,^145^ and the actin cytoskeleton modulator, ROCK2. ROCK2, also targeted by the anti-fibrotic miR-335-3p and miR-30e-3p in our study, regulates actin cytoskeleton dynamics, ECM remodeling, endothelial–mesenchymal transition, and vascular permeability. ROCK2 activation downstream of TGF-β2 and CCN2 contributes to fibrosis, endothelial activation, and vascular remodeling,^146,147^ and, therefore, has been implicated in hypertension, hypertrophy, and ischemia/reperfusion injury^148^. Therefore, ROCK2 inhibition emerges as a promising cardioprotective strategy, since regulators such as miR-335-3p can attenuate TGF-β1–induced fibrosis and may offer fewer side effects than current ROCK inhibitors.^148,149^ Furthermore, the established role of ROCK2 in senescence also positions its targeting as a potential senomorphic therapeutic approach.^150^

Another critical member of this regulatory layer is the miR-29 family, a key miRNA for maintaining ECM integrity by targeting collagens, fibrillins, and elastin. Therefore, their downregulation has been implicated in endothelial dysfunction,^151^ myocardial infarction, and ischemia–reperfusion injury.^152,153^ However, the persistent repression of collagen and elastin by miR-29 in aging tissues, including murine aortas, has been shown to weaken vascular integrity and promote aneurysms, hypertrophy, and cardiac insufficiency.^93,154^ Therefore, miR-29 levels must be tightly regulated, as both insufficient expression and excessive suppression of ECM proteins can respectively drive fibrosis or compromise vascular wall stability, each contributing to CVD. In our study, miR-29c-3p and miR-29b-3p were found to be downregulated in senescent cells. In contrast, EVs from senescent cells displayed an anti-fibrotic phenotype, carrying higher levels of miR-29a-3p and reduced pro-fibrotic onco-miRs, such as miR-1290 and miR-1246, which are associated with endothelial–mesenchymal transition in restenotic arteries^155^ and endothelial barrier breakdown,^156^ respectively. We also observed correlations between these miRNAs in both senescent HUVECs and EVs. miR-29c-3p was negatively correlated with ADAMTS6, a metalloproteinase involved in ECM remodeling and recently implicated in cardiovascular development,^157^ supporting the involvement of the miR-29 family in the ECM remodeling alterations observed in our senescent HUVECs. In senescent EVs, miR-29a-3p levels were inversely correlated with the antioxidant enzyme GPX7, an antioxidant enzyme absent in late-passage EVs, suggesting a role in propagating redox imbalance and oxidative injury. Reinforcing this, previous studies showed that GPX7 upregulation mitigates vascular aging and cellular senescence,^158^ whereas reduced GPX activity promotes lipid peroxidation, thereby contributing to endothelial dysfunction, atherogenesis, and arrhythmia^159^. Furthermore, we recently reported that GPX activity in endothelial and platelet MVs from CKD patients inversely correlated with tissue factor, a pro-coagulant mediator linked to increased cardiovascular risk.^30^

Finally, the repression of miR-29b-3p, together with miR-335-3p and miR-106, intersects with transcriptional control through REST, a master repressor of neuronal genes in non-neuronal tissues.^160^ REST plays multifaceted roles in aging and senescence, supporting neuronal longevity and protection, and its dysregulation drives senescence and neurodegeneration.^160–162^ It also influences cardiac and vascular physiology by modulating ion channels and smooth muscle proliferation.^160^ Recently, REST was shown to target ECM proteins such as CD44, underscoring its role as a modulator of ECM remodeling, epithelial–mesenchymal transition, and phenotypic reprogramming.^163,164^ Paradoxically, REST upregulation in our senescent HUVECs was accompanied by increased expression of its targets, including neuronal-like markers and ECM genes such as CD44,^163,164^ whereas C-terminal domain small phosphatase 1, an essential stabilizer of REST activity,^165^ was detected only in early-passage EVs. This paradox suggests that REST dysregulation may contribute to phenotypic drift and its potential EV-mediated propagation, but may also represent a compensatory attempt to restore EC identity. Indeed, recent evidence has shown that REST can be recruited by ETV2, a master regulator of EC specification, to preserve endothelial identity and restrict alternative phenotypes.^166^ Collectively, these findings emphasize REST modulation as a potential therapeutic strategy for vascular aging and CVD, warranting further investigation in the context of RES.

## 4. Conclusion

Using an integrated multi-omics strategy for the characterization of senescent HUVECs and EVs, we established an integrative molecular framework that links molecular endothelial senescence to age-related diseases such as CVD. Senescent HUVECs and EVs exhibited a canonical senescence signature together with a pronounced suppression of RNA metabolism, ribosome biogenesis, and DNA-repair pathways, aligning with newly emerging hallmarks of cellular aging. Omics analyses have also revealed angiogenic and ECM mediators of endothelial dysfunction, underscoring the role of senescence in CVD. Senescent EVs further amplify this dysfunction by losing reparative factors and carrying pro-fibrotic and angiogenic mediators.

Multi-omics profiling revealed consistent transcript–protein changes in senescent cells and EVs, helping to define a robust molecular signature of RES, encompassing EC-specific senescence markers (e.g., ADIRF, TGF-β2, CD44, IGFBPs, BGN, ITGA4, and MT2A) and post-transcriptional regulators, including LUC7L3, MALAT1, and miRNAs (miR-22-3p, miR-126-5p). In particular, miR-22-3p and miR-126-5p emerged as key modulators of replicative senescence in both cells and EVs, with miR-22-3p exhibiting compartment-specific regulations. Finally, integrative proteomic and transcriptomic (mRNA and miRNA) analyses revealed a tightly coordinated regulatory network involving multiple TFs and miRNAs that orchestrate endothelial dysfunction. Several miRNAs (miR-23, miR-335-3p, miR-29 family, miR-590-3p, miR-126-5p) were negatively correlated with multiple TFs (REST, KLFs, and ZNFs) and other downstream targets, collectively driving endothelial senescence through impaired PI3K/Akt signaling, maladaptive angiogenesis, ECM remodeling, and broader transcriptional dysregulation. Although further studies are required to validate the functional consequences of these regulatory interactions *in vivo*, our findings provide new mechanistic insights into endothelial senescence and uncover potential avenues for modulating secondary senescence and age-related vascular diseases.

## 5. Experimental Section/Methods

### 5.1. Human umbilical vein endothelial cells culture

HUVECs were provided by Lonza (CC-2517, lot number 323352) from a pooled primary cell frozen at passage 1 in a cryopreservation medium containing endothelial growth medium (EGM) with 10% heat-inactivated fetal bovine serum (FBS) (Sigma). Cultures were grown at 37°C in a 5% CO2 atmosphere at 95% humidity in EGM consisting of endothelial basal media (EBM; Lonza CC-3121) supplemented with a growth bullet kit (Lonza, CC-4133) containing Bovine Brain Extract, ascorbic acid, hydrocortisone, epidermal growth factor, gentamicin/amphotericin B, and supplemented with 10% heat-inactivated FBS were used.

### 5.2. Replicative senescence model

First-passage cryopreserved HUVECs were cultured and serially passaged until reaching senescence, following the replicative senescence protocol previously described by our group.^26^ The rate of population doubling (PD) between passages was calculated using the formula: PD = [ln {number of cells harvested} − ln {number of cells seeded})/ln2]. Cells studied within 2–8 passages (early passage; PD < 20) were regarded as early passage HUVECs, whereas those passaged 28–38 times (PD > 96) were regarded as senescent HUVECs. SA-β-gal staining was performed using the Senescence-Galactosidase Staining Kit (Abcam) following the manufacturer’s instructions to evaluate the percentage of senescent cells.

### 5.3. Isolation, quantification, and characterization of extracellular vesicles

EVs were isolated from the culture media of early passage and senescent HUVECs, pooled into 3 samples per condition, and diluted with filtered (0.2 μm) PBS (Sigma), as previously described.^27^ Briefly, cell culture supernatants were subjected to serial centrifugation in a Centrifuge 5427R (Eppendorf, Hamburg, Germany): 15 min at 1,400 x g to remove cells and cellular debris, followed by centrifugation for 30 min at 20,000 x g to concentrate the EVs. EVs were immediately processed or stored at -20°C in filtered (0.2 μm) PBS 1x (Sigma). Finally, the isolated EVs were divided into three pools per condition: 1) EVs from early endothelial passage cells (pools #1, 2, and 3) and 2) EVs from senescent HUVECs (pools #4, 5, and 6).

EVs from endothelial cell cultures were characterized based on their surface markers (FC), particle size (NTA), pan-EV-specific markers (FC and WB), and morphological analysis by TEM following MISEV2023,^19^ with particular attention to guidelines for cell culture-derived EVs.^167^ GraphPad Prism 8.0.2 (GraphPad Software, La Jolla, CA, USA) was used for data visualization and statistical analysis. For TEM and NTA data, normality was assessed using the Shapiro–Wilk test, followed by unpaired t-tests. Data were analyzed using two-way ANOVA to evaluate the effects of cell condition (senescent vs. control) and tetraspanin markers (CD9⁺, CD63⁺, and CD81⁺) on EV populations.

The size distribution and enumeration/concentration of EVs (quantity of isolated EVs) were calculated using NTA with a NanoSight NS300 system (Malvern Panalytical). EV samples were quantified in terms of size, number of particles, and concentration through three independent measurements using real-time visualization and individual particle tracking. The samples were diluted and 1:10-1:20 in particle-free PBS according to the manufacturer’s recommendations for image capture by NTA analysis. The images of the EVs were captured in 10 s movies, repeated three times for each sample, and tracked particles were automatically calculated (particle size) and graphed according to Brownian motion and the diffusion coefficient (Dt). The results are displayed as a frequency size distribution graph and output to a spreadsheet. The results obtained were in “finite track-length adjusted” (FTLA) size per concentration graph and averaged FTLA size per concentration graph with the standard error of the mean. Data were analyzed using the NTA 3.2 Dev Build 3.2.16 software (Malvern Panalytical).

EV pools from early passage and senescent HUVECs (n=3) were resuspended in filtered PBS. For negative staining, glow discharge was performed in duplicate on carbon-coated collodion 400 mesh nickel grids (Gilder). Fifteen microliters of each sample was adsorbed onto the grid for 2 min. The grids were then washed with two drops of Milli-Q water, stained with 20 µL of 2% aqueous uranyl acetate (Electron Microscopy Sciences) for 1 min, and dried at room temperature. The grids were visualized using a JEOL JEM 1400 flash electron microscope (operating at 100 kV). Micrographs were taken using Digital Micrograph software (GATAN) OneView digital camera at various magnifications.

To measure the EVs by TEM, GATAN version 3.51.3720.0 was used, taking the exterior and interior measurement diameters of the vesicles (n=20) for each sample (n=3 per condition). Measurements were performed on vesicles in which both diameters were easily visible. Data are expressed as mean ± standard deviation and were analyzed using an unpaired t-test. P values less than 0.02 were considered statistically significant.

EVs were diluted in filtered PBS (0.1 μm; Sigma) and incubated with 25 μM Tag-it Violet (TIV+) (BioLegend) for 90 min at 37°C in the dark for membrane labeling to discriminate the background. Tetraspanin analysis (pan-EV makers), including anti-CD9-FITC (Beckman Coulter), anti-CD81-PE (Beckman Coulter), anti-CD63-PE (Beckman Coulter), and antibody mix of anti-CD9-488/anti-CD81-555/anti-CD63-647 (Leprechaun Kit-Unchained labs), were performed for EVs that were incubated for one hour at room temperature in the dark. In addition, a negative control was included in which the EV samples were treated with 0.1% Triton X-100 (Sigma) for 15 min at RT. Samples were analyzed using a CytoFLEX S (Beckman Coulter) previously calibrated with NIST polystyrene (PS) sphere assembly for scattering calibration: 80-100-150-200-300-400 nm (Thermo Fisher) and SPHEROTM Rainbow Calibration Particles, 4 peaks, 0.4-0.6 µm (RCP-05-5; Spherotech). The FCM Pass software was used to estimate the EV size based on the refractive index for size calibration. The flow cytometer CytoFLEX S was routinely calibrated following the International Consortium of MIFlowCyt-EV^168^ guidelines and tested daily for quality control.

#### 5.3.4. Western blot analysis

Pooled EV samples from early and senescent HUVECs were lysed with RIPA buffer (Santa Cruz Biotechnology) and quantified using a BCA Protein Assay Kit (Thermo Scientific, 23225), as described previously. The absorbance was measured using a plate reader (FLUOstar Omega, BMG Labtech). Briefly, equal amounts of protein (12 μg protein/lane) were diluted with Laemmli sample buffer under reducing conditions (with β-mercaptoethanol) for TSG101 protein and under non-reducing conditions for CD9/CD63 protein. Proteins were separated at 25 mA on Mini-Protean precast gels 4-20% (Bio-Rad). Samples were then transferred onto nitrocellulose membranes with a Trans-Blot Turbo Transfer System (BioRad), blocked with PBS 1X and 0.1%Tween 20 (PBS-T) containing 5% BSA, and then incubated overnight at 4°C with shaking, with primary antibodies diluted in the same buffer (1/1000). The primary antibodies used were the Exosome Panel (ab275018; Abcam; TSG101 and CD9). The following day, the membranes were washed three times with PBS-Tween and incubated with Anti-Rabbit IgG-HRP-linked secondary antibody for 1 h, followed by three additional washes with PBS-T. Bands were visualized using the SuperSignal West Pico PLUS Chemiluminescent substrate, and digital images were captured using an Amersham Imager 600.

### 5.4. Proteomics analysis

Senescent and early passage HUVECs (n=3 per biological replicate) and secreted EVs were lysed with a lysis buffer containing 5% sodium dodecyl sulfate (SDS), 25 mM triethylammonium bicarbonate (TEAB), 5 mM tris(2-carboxyethyl) phosphine (TCEP), and 10 mM chloroacetamide (CAA)), supplemented with Pierce™ DNase (25 kU, 88701, ThermoFisher). Lysates were sonicated using an ultrasonic processor UP50H (Hielscher Ultrasonics) for 1 min on ice (0.5 cycles, 100% amplitude) and centrifuged at 18,000 × g for 10 min. The supernatants were incubated at 60°C for 30 min to reduce alkylated cysteine residues. The protein concentrations of the supernatants were determined using Pierce™ 660 nm Protein Assay Reagent was supplemented with Ionic Detergent Compatibility Reagent (Thermo Scientific) according to the manufacturer’s instructions. Protein digestion on S-Trap columns (Protifi) was performed according to the manufacturer’s instructions, with minor changes.^169,170^ Briefly, 20 µg of each sample was digested overnight at 37°C at a trypsin-to-protein ratio of 1:15. Tryptic peptides were cleaned up using Stage-Tips prepared from Octadecyl C18 solid-phase extraction disks (Empore™, 66883-U), as previously described.^170^ The eluted peptides were dried in a speed vacuum and quantified by fluorimetry (QuBit) according to the manufacturer’s instructions.

For nano-liquid chromatography coupled with Electrospray Ionization Tandem Mass Spectrometry (nanoLC-ESI-MS/MS) analysis, 1 µg of each sample was individually analyzed using an Ultimate 3000 nano HPLC system (Thermo Fisher Scientific) coupled online to an Orbitrap Exploris 240 mass spectrometer (Thermo Fisher Scientific). Each sample (1 µg in 5 µL of sample resuspended in mobile phase A: 0.1% formic acid (FA)) was loaded onto a 50 cm × 75 μm Easy-spray PepMap C18 analytical column at 45°C. Tryptic peptides were separated at a flow rate of 250 nL/min using a 120-minute gradient ranging from 4% to 95% mobile phase B (80% acetonitrile (ACN) in 0.1% FA). To avoid carry-over, two 40-minute blank samples (mobile phase A) were systematically run between the samples. Data acquisition was performed using a data-dependent top-20 method in the full-scan positive mode, scanning 375–1200 m/z. MS1 scans were acquired at an Orbitrap resolution of 60,000 at m/z 200, with a normalized automatic gain control (AGC) target of 300%, a radio frequency (RF) lens of 80%, and automatic maximum injection time (IT). The top 20 most intense ions from each MS1 scan were selected and fragmented with a high-energy collisional dissociation (HCD) of 30%. The resolution of the HCD spectra was set to 15,000 at m/z 200, with a normalized AGC target of 50% and an automatic maximum IT. The precursor isolation window was 1 m/z and a dynamic exclusion of 45 s was applied. Precursor ions with single, unassigned, or six or higher charge states from the fragmentation selection were excluded.

Raw data files were processed using the Proteome Discoverer 2.5.0.400 software (Thermo Scientific, Bremen, Germany), and database search was carried out using four search engines (Mascot (v2.8.0), MsAmanda (v2.4.0), MsFragger (v3.1.1), and Sequest HT) against a Homo Sapiens UniProtKB reviewed database (19th February 2021, 20,378 sequences) containing the most common laboratory contaminants (cRAP database with 69 sequences). The search parameters were set as follows: cysteine carbamidomethylation (+57.021464 Da), methionine oxidation (M) (+15.994915 Da), N-term acetylation (+42.010565 Da), and Gln→pyro-Glu (-17.026549 Da) as variable modifications. Precursor mass tolerances were set at 10 ppm and the fragment mass tolerance at 0.02 Da and trypsin/P was selected as a protease with a maximum of 2 missed cleavage sites. The False Discovery Rate (FDR) was calculated using the processing node Percolator (maximum delta Cn 0.05; decoy database search target) and the validation of proteins, peptides, and peptide spectral matches (PSMs) with a False Discovery Rate (FDR) ≤1%. Precursor ion quantitation was also performed in Proteome Discoverer using the “Minora” feature in the processing method and the “Feature Mapper” and “Precursor Ions Quantifier” nodes in the consensus step. The Feature Mapper node in the consensus method was used to create features from unique peptide-specific peaks by applying chromatographic retention time alignment with a maximum shift of 10 min and a signal-to-noise threshold of 5. The following parameters were set in the “Precursor Ions Quantifier” node: unique+razor peptides were used for quantitation, precursor abundance was based on intensity, and the normalization mode was based on the total peptide amount. Protein abundance was calculated by summing the sample abundance of the connected peptide groups. Protein abundance was calculated by summing the sample abundance of the connected peptide groups. Pairwise-based ratios were calculated, and a t-test background-based was performed as a statistical test. To determine the differentially expressed proteins (DEPs) between senescent and early passage HUVECs and isolated EVs, the following filters were considered: quantitation by protein group (only master proteins were selected) with a protein FDR confidence of high, normalized protein abundance was detected in at least three samples per group, the percentage of coefficient of variation of abundance values should have a value in each sample group, and the p-value adjusted using Benjamini–Hochberg correction was set at ≤0.05. A volcano plot and Principal Component Analysis (PCA) were also performed using Proteome Discover. DEPs were analyzed using Metascape^172^ and STRING 11.5^173^ for functional enrichment based on gene ontology (GO), KEGG and REACTOME pathways/reactions, and protein-protein interactions.

### 5.5. Transcriptomics analysis

The samples analyzed in this study were senescent and early passage HUVECs (n=3 per biological replicate) and secreted EVs (n=3 per biological replicate). Total RNA was extracted from the cell pellets using the MagMAX™ mirVana™ Total RNA Isolation Kit (Thermo Fisher Scientific, Waltham, MA, USA) according to the manufacturer’s instructions. EVs were processed using the Total Exosome RNA and Protein Isolation Kit (Invitrogen), following the manufacturer’s recommendations. Dual-indexed mRNA libraries were prepared using the QuantSeq 3’ mRNA-Seq V2 Library Prep Kit with UDI (Lexogen GmbH, Vienna, Austria), according to the user guide. According to the manufacturer’s recommendations, 250 ng of starting material was used for cellular RNA analysis, while a low-input protocol (<10ng total) was used for EVs. Library quality was evaluated using the QiaXcel Advanced System (Qiagen), while the quantity was measured using a Quantus Fluorometer (Promega Corporation). Sequencing was performed using a NovaSeq X Plus System (Illumina, San Diego, CA, USA). The read length for the paired-end run was 2 × 150 bp and the target coverage was 5 M reads for each library. For miRNA, libraries of small RNA were constructed using the NEXTFlex Small RNA-Seq Kit v4 (Illumina Compatible) (PerkinElmer, Waltham, MA, USA), according to the manufacturer’s guidelines. Size selection was performed by targeting fragments that were 150–180 bp in size using BluePippin (Sage Science, Beverly, Massachusetts, USA). Size-selected libraries were then PCR-amplified with indexed sequencing primers and finally sequenced on a NovaSeq X Plus System (Illumina, San Diego, California, USA) for 150 × 2 cycles in high-output mode with a target coverage of 20 M reads for each library.

*R* (version 4.1.2) was used to perform differential enrichment analyses. For mRNA-Seq and miRNA-Seq data, count values were imported and processed using *edgeR* (version 3.36.0).^174^ Expression values were normalized using the trimmed mean of M values (TMM) method and lowly-expressed genes (< 1 counts per million) were filtered out. Differentially expressed genes were identified using linear models (*Limma-Voom*) (version 3.50.3)^175^, and *P* values were adjusted for multiple comparisons by applying the Benjamini-Hochberg correction. For proteomics, the data were log-transformed before plotting and differential enrichment analyses. Heatmaps and volcano plots were generated using the *heatmap3* (version 1.1.9)^176^, *EnhancedVolcano* (version 1.12.0) (https://github.com/ kevinblighe/EnhancedVolcano), and *Glimma* (version 2.4.0)^177^ packages. Gene functional annotation was performed using *clusterProfiler* (version 4.2.2)^178^ and *rrvgo* (version 1.6.0)^179^ packages. Correlation matrix plots were generated using *corrplot* (version 0.92). miRNA targets were obtained from miRDB (version 6.0)^180^. Functional reduction heatmaps were generated using *rrvgo*.

## Abbreviations

ADIRF: Adipogenesis Regulatory Factor
ANGPT: Angiopoietin
BGN: Biglycan
BDKRB2: Bradykinin Receptor B2
CVD: Cardiovascular Diseases
CKD: Chronic Kidney Disease
COL: Collagen
C3: Complement C3
C4A: Complement C4A
CDK: Cyclin-Dependent Kinase
DEGs: Differentially Expressed Genes
DEPs: Differentially Expressed Proteins
EndMT: Endothelial–Mesenchymal Transition
ECM: Extracellular Matrix
EVs: Extracellular Vesicles
FGF: Fibroblast growth factor
FGFR1: Fibroblast growth factor receptor 1
FC: Flow Cytometry
GPX7: Glutathione Peroxidase 7
HMGB2: High Mobility Group Box 2
HMGs: High Mobility Group Proteins
HUVECs: Human Umbilical Vein Endothelial Cells
PTK7: Inactive Tyrosine-Protein Kinase 7
IGF: Insulin-like Growth Factor
IGFBP: Insulin-like Growth Factor-Binding Protein
ITGA4: Integrin alpha-4
KLF: Krüppel-like Factors
lncRNA: Long non-coding RNA
MT2A: Metallothionein 2A
MALAT1: Metastasis-Associated Lung Adenocarcinoma Transcript 1
MVs: Microvesicles
NTA: Nanoparticle Tracking Analysis
NCAN: Neurocan
NCL: Nucleolin
PAPPA: Pappalysin
PDGF: Platelet-derived growth factor
PPI: Protein–Protein Interaction
REST: RE1-Silencing Transcription Factor
RES: Replicative Endothelial Senescence
RETN: Resistin
ARHGAP42: Rho GTPase-activating Protein 42
ROCK2: Rho-associated Protein Kinase
SAHF: Senescence-associated Heterochromatin Foci
SASP: Senescence-Associated Secretory Phenotype
SA-β-gal: Senescence-Associated β-Galactosidase
tPA or PLAT: Tissue-type plasminogen activator
TF: Transcription factor
TGF-β: Transforming Growth Factor Beta
TEM: Transmission Electron Microscopy
VEGF: Vascular Endothelial Growth Factor
ZNF: Zinc Finger Factors

## Acknowledgements

Extracellular Vesicle characterization and analysis were performed at the Extracellular Vesicles Unit (UVEx) of the Hospital Nacional Parapléjicos (SESCAM). In addition, we acknowledge Dr. Luis F. Lorenzo from EGO Genomics for their assistance with bioinformatics analysis and Andrea Figuer for her technical support.

## Data Availability Statement

The datasets generated and analyzed during this study will be deposited in the Gene Expression Omnibus (GEO) and will be made publicly available prior to publication in a peer-reviewed journal.

Received: ((will be filled in by the editorial staff)) Revised: ((will be filled in by the editorial staff)) Published online: ((will be filled in by the editorial staff))

## Supporting Information

Supporting Information is available from the Wiley Online Library or from the author.

Supplementary Figures S1 – Supplementary Figures.

## Supplementary Tables

**Supplementary Table 1**. Extracellular vesicles (EVs) derived from senescent and early-passage cells were characterized using nanoparticle tracking analysis (NTA), transmission electron microscopy (TEM), and flow cytometry (FC).

**Supplementary Table 2**. Proteins identified (ID), quantified (QUANT), and differentially expressed (DEPs) in senescent and early-passage endothelial cells and secreted extracellular vesicles (EVs).

**Supplementary Table 3**. Functional analysis of differentially expressed proteins (DEPs) in senescent and early-passage endothelial cells and secreted extracellular vesicles (EVs).

**Supplementary Table 4**. Raw RNA-seq read counts and differentially expressed genes (DEGs) in senescent and early-passage endothelial cells and secreted extracellular vesicles (EVs).

**Supplementary Table 5**. Functional analysis of differentially expressed genes (DEGs) in senescent and early-passage endothelial cells.

**Supplementary Table 6**. Functional analysis of differentially expressed genes (DEGs) in senescent and early-passage extracellular vesicles (EVs).

**Supplementary Table 7**. Molecular integration of proteomics and transcriptomics analysis of senescent and early-passage endothelial cells and their extracellular vesicles (EVs).

**Supplementary Table 8**. Differentially expressed miRNAs in senescent and early-passage endothelial cells and their extracellular vesicles (EVs), with associated functional analysis.

**Supplementary Table 9**. Putative and experimental correlations between miRNAs and differentially expressed genes/proteins in senescent and early-passage endothelial cells and their extracellular vesicles (EVs).

---

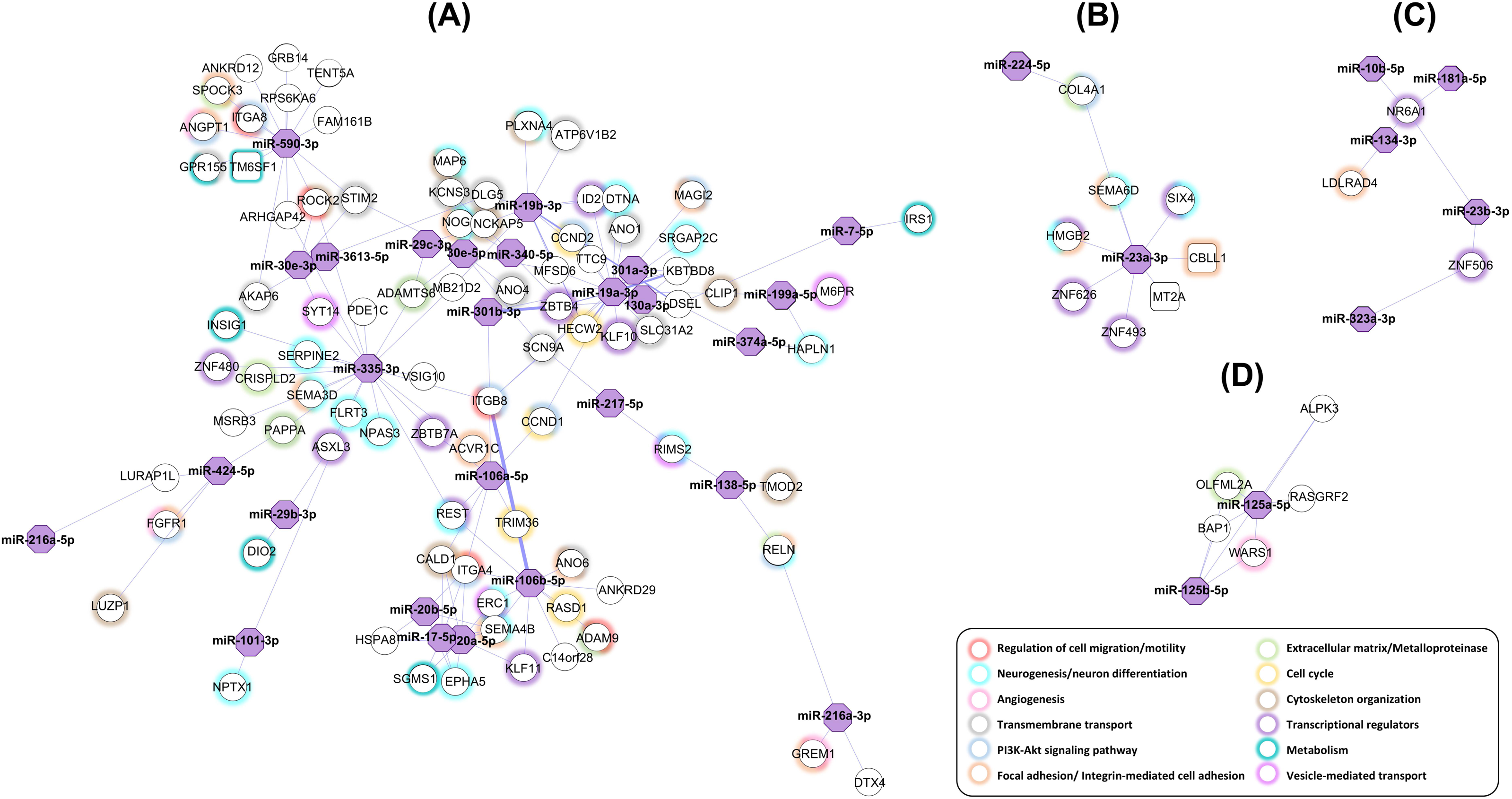

