## Supplemental figures for "Multi-omics integrative approach of senescent endothelial cells and derived extracellular vesicles in a replicative senescence model"

### Supplementary Figures

**Supplementary Figure S1: Replicative human senescent endothelial cell model.**

**A)** Representative pictures of early passage (left) and senescent (right) human endothelial cells showing changes in their morphology (total magnification 50 µm). **B)** Human early passage and senescent endothelial cell β-galactosidase staining. 10x objective (total magnification, 100×). Representative images of early passage (left) and senescent (right) endothelial cells (HUVECs) are shown. β-Galactosidase staining shows senescent cells in blue, validating the replicative senescence model. HUVEC Passage 7: PD<20 and HUVEC Passage 29: PD>96.

**A)** **Early passage endothelial cells** **Senescent endothelial cells**

**B)**

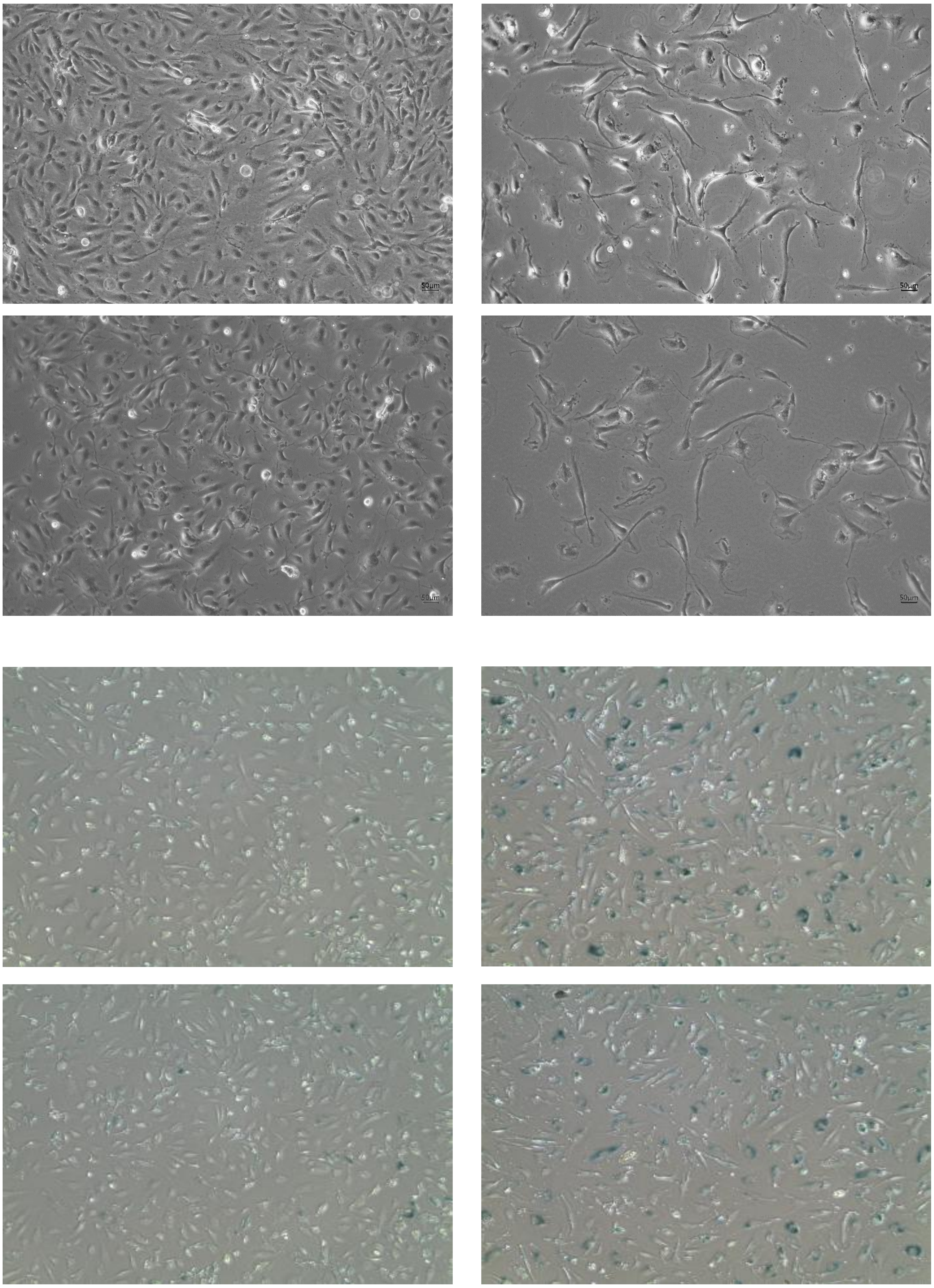

**Supplementary Figure S2: Characterization of the extracellular vesicles (I) by NTA.** Extracellular vesicle (EVs) concentration and size (mean and mode) from early passage (A: pool#2 and C: pool#3) and senescent (B: pool#5 and D: pool#6) human endothelial cells were measured using nanoparticle tracking analysis (NTA).

#### A) Early passage EVs (pool#2)

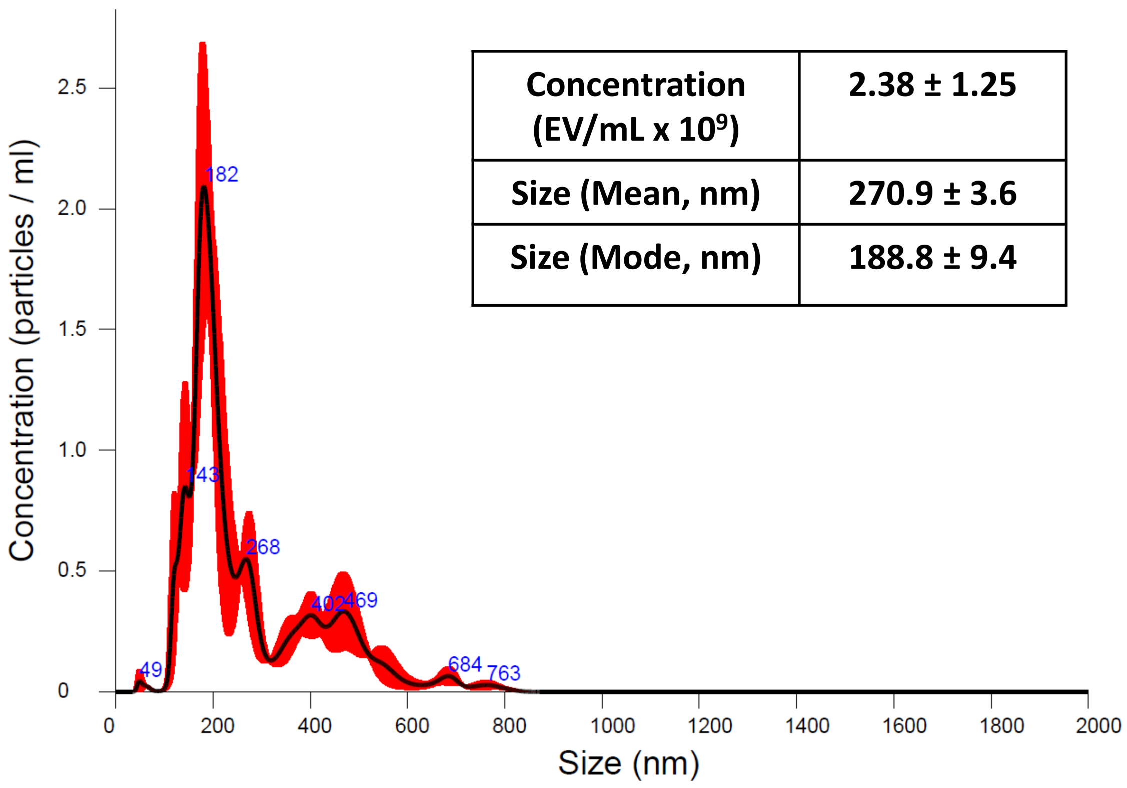


| Concentration (EV/mL x $10^9$) | 2.38 ± 1.25 |
|---|---|
| Size (Mean, nm) | 270.9 ± 3.6 |
| Size (Mode, nm) | 188.8 ± 9.4 |

#### B) Senescent EVs (pool#5)

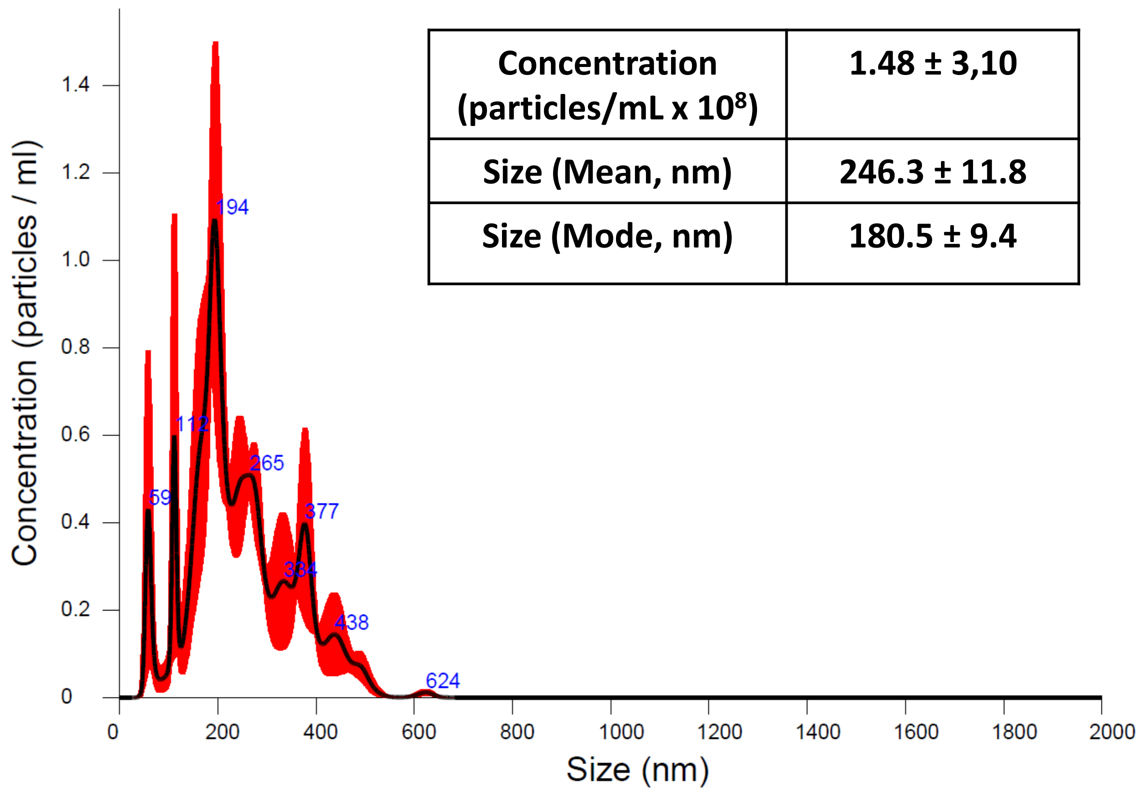


| Concentration (particles/mL x $10^8$) | 1.48 ± 3,10 |
|---|---|
| Size (Mean, nm) | 246.3 ± 11.8 |
| Size (Mode, nm) | 180.5 ± 9.4 |

#### C) Early passage EVs (pool#3)

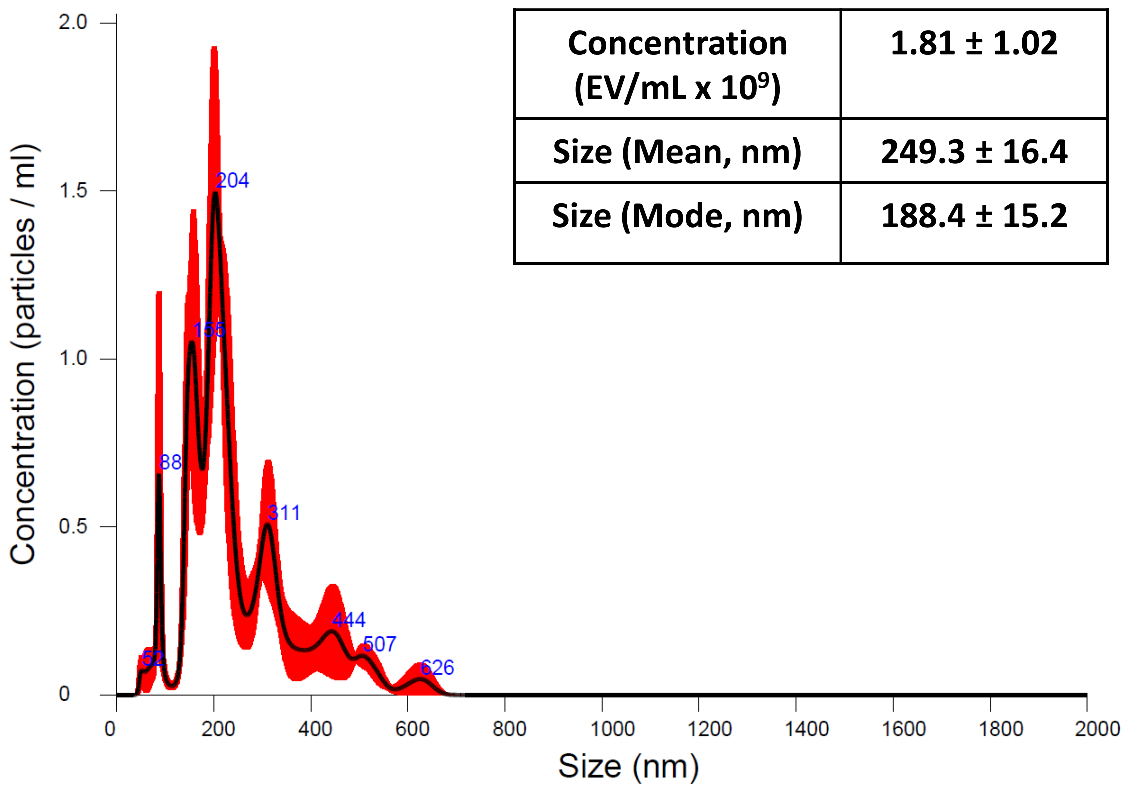


| Concentration (EV/mL x $10^9$) | 1.81 ± 1.02 |
|---|---|
| Size (Mean, nm) | 249.3 ± 16.4 |
| Size (Mode, nm) | 188.4 ± 15.2 |

#### D) Senescent EVs (pool#6)

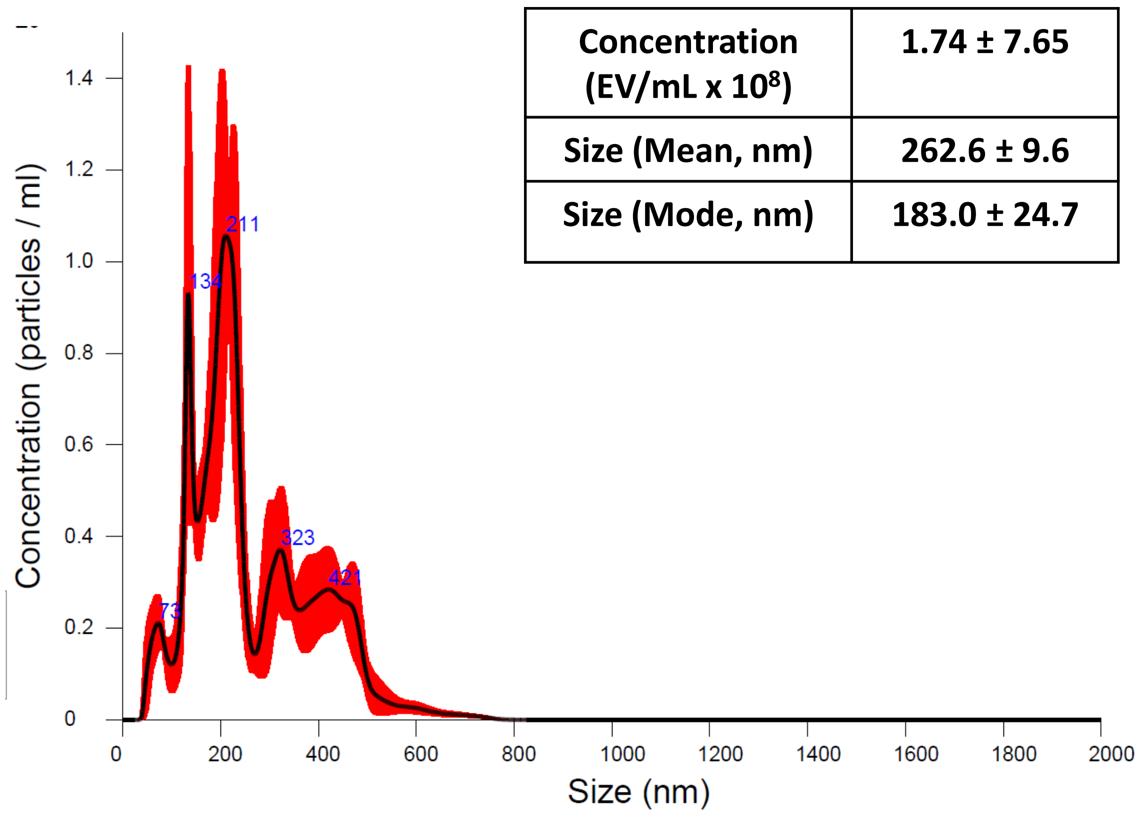


| Concentration (EV/mL x $10^8$) | 1.74 ± 7.65 |
|---|---|
| Size (Mean, nm) | 262.6 ± 9.6 |
| Size (Mode, nm) | 183.0 ± 24.7 |

**Supplementary Figure S3: Characterization of the extracellular vesicles (II) by TEM.**

**A) Early passage EVs**

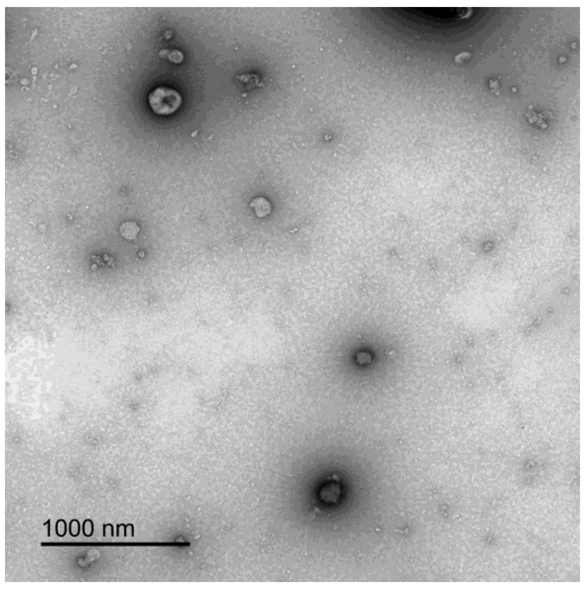


**(pool#1)**

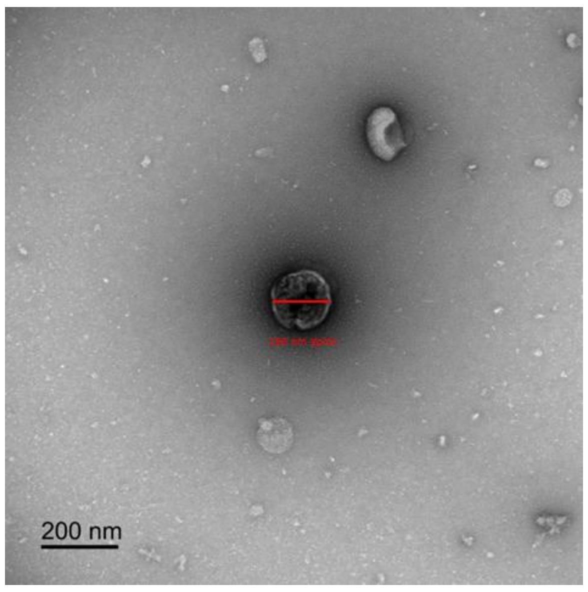


**(pool#2)**

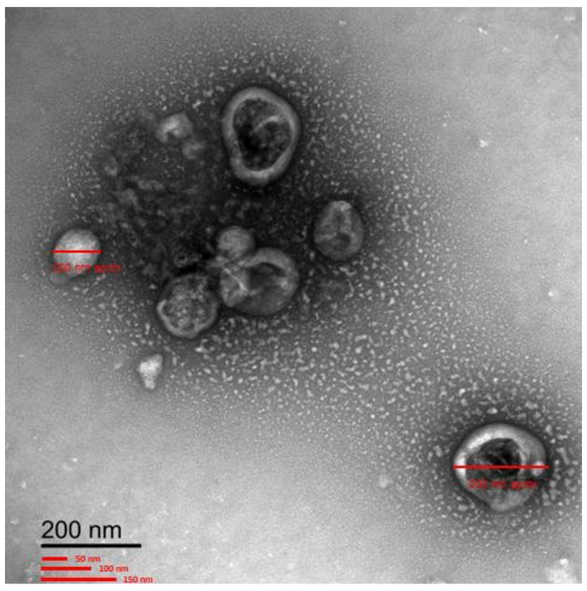


**(pool#3)**

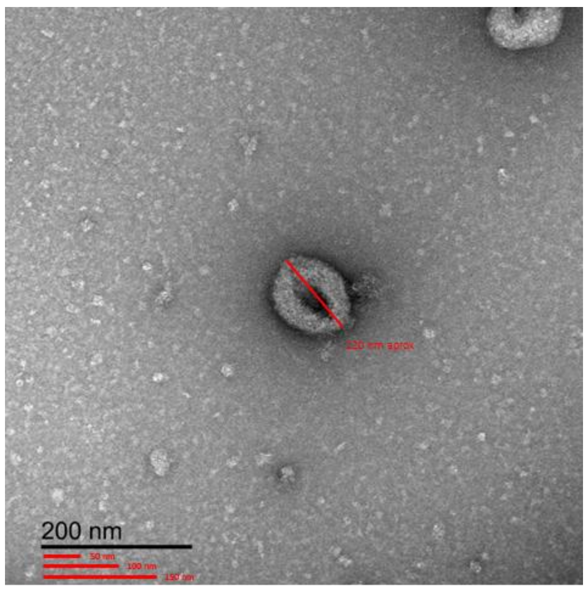


**B) Senescent EVs**

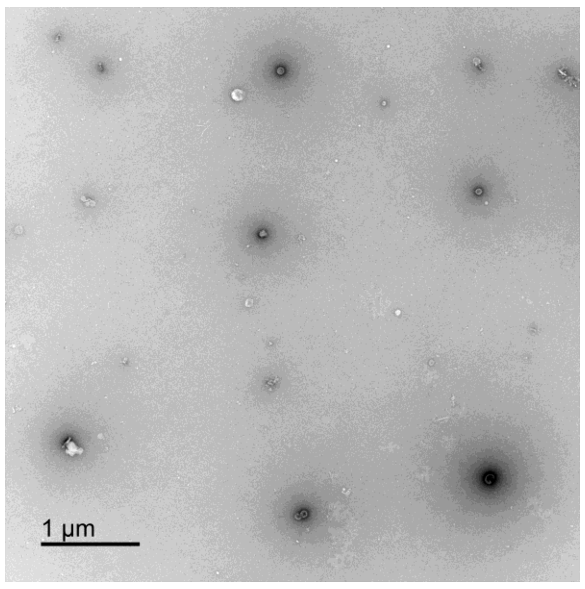


**(pool#4)**

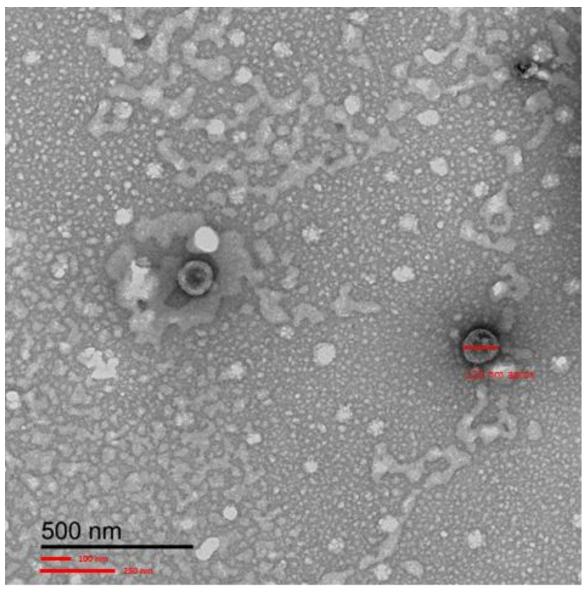


**(pool#5)**

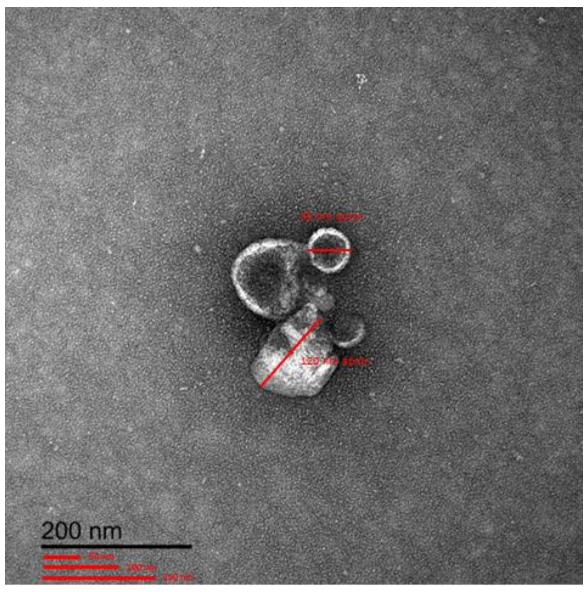


**(pool#6)**

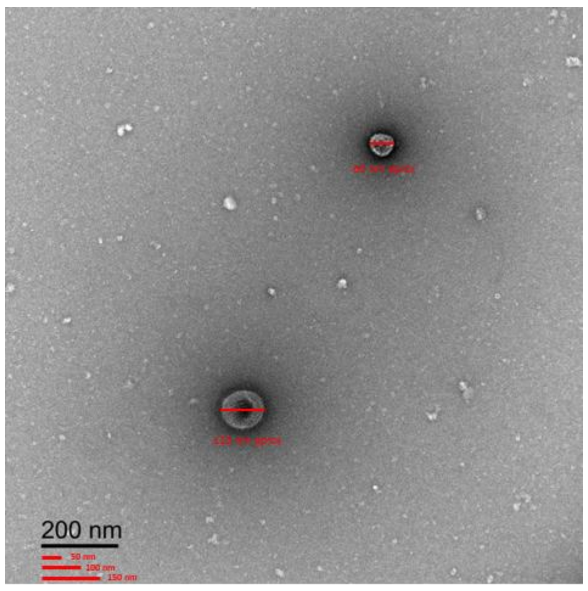

**Supplementary Figure S4a: Characterization of the extracellular vesicles (III) by Flow cytometry.**

EVs from early passage (pools #2 and 3) and senescent (pools #5 and 6) human endothelial cells were characterized by flow cytometry (FC), defining the EV as the events Tag-It Violet (TIV+) for assessing the expression of specific pan-EV marker (tetraspanins: CD9+, CD63+, and CD81).

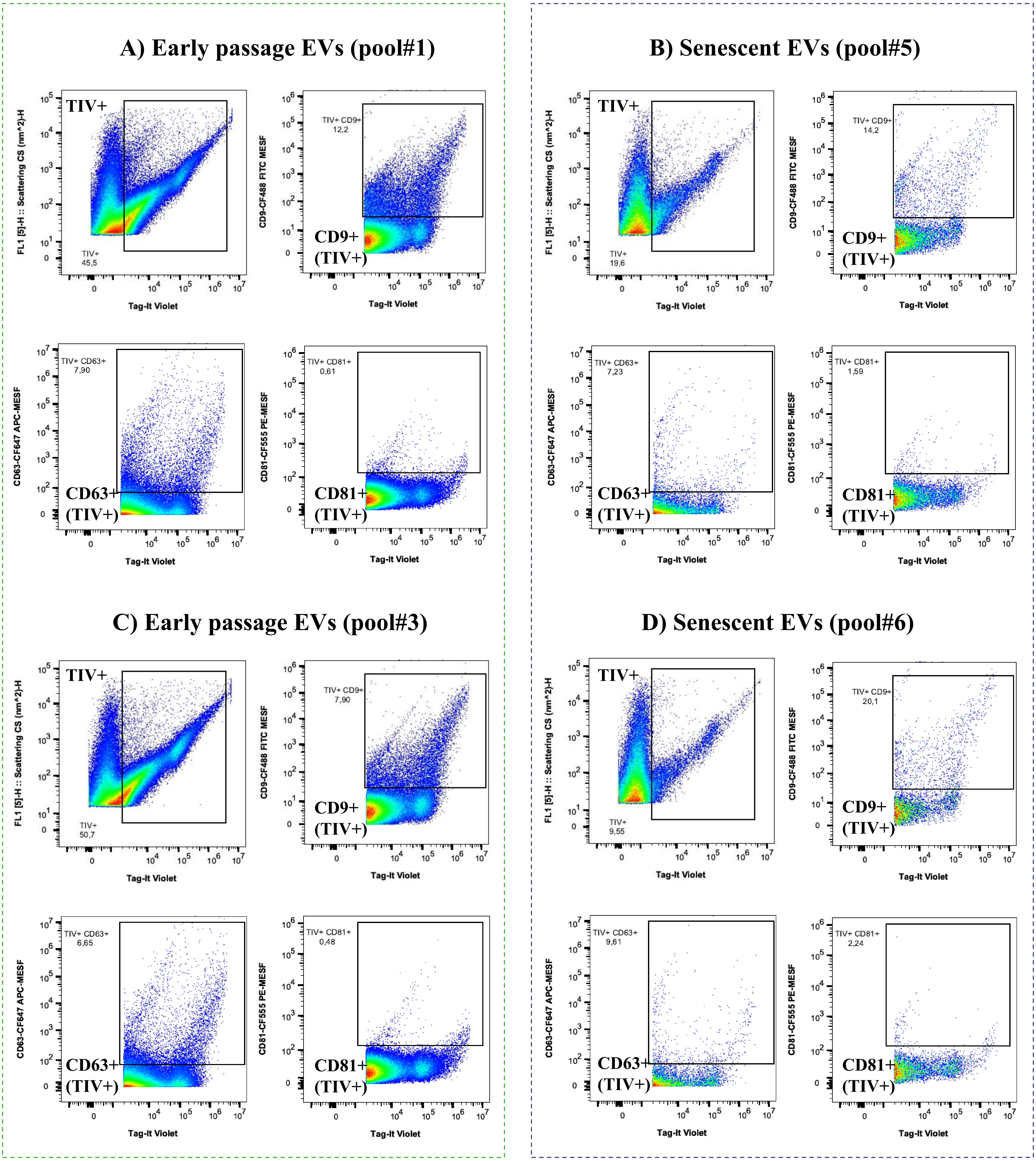

**Supplementary Figure S4b: Characterization of the extracellular vesicles (III) by Flow cytometry.**

Percentage of expression of specific pan-EV marker (tetraspanins: CD9+, CD63+, and CD81) in EVs from early passage and senescent human endothelial cells

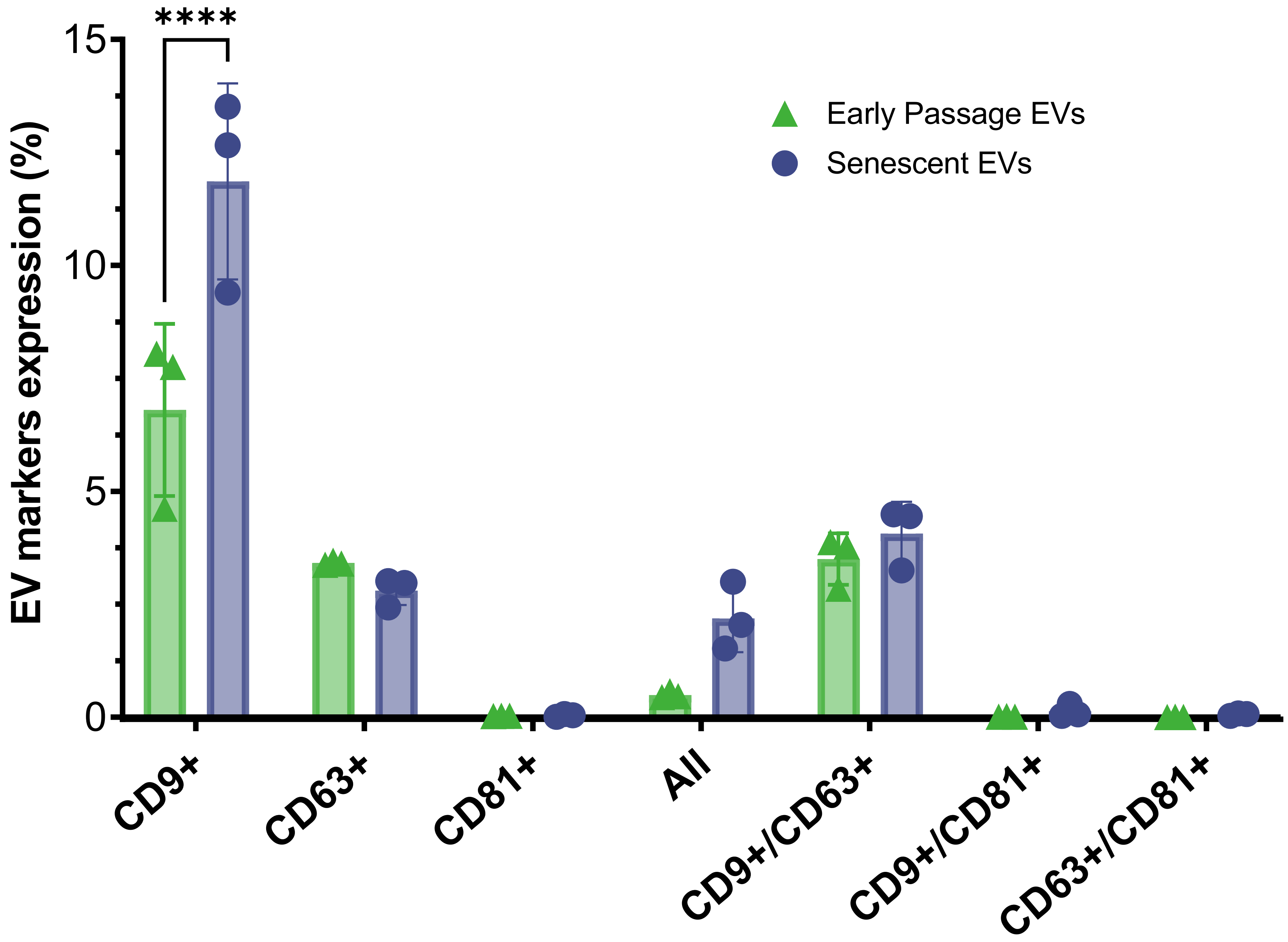

**Supplementary Figure S5a: Characterization of the extracellular vesicles (IV). Measurement of CD9+, CD63+, and CD81+ and other markers by LC-MS/MS.**

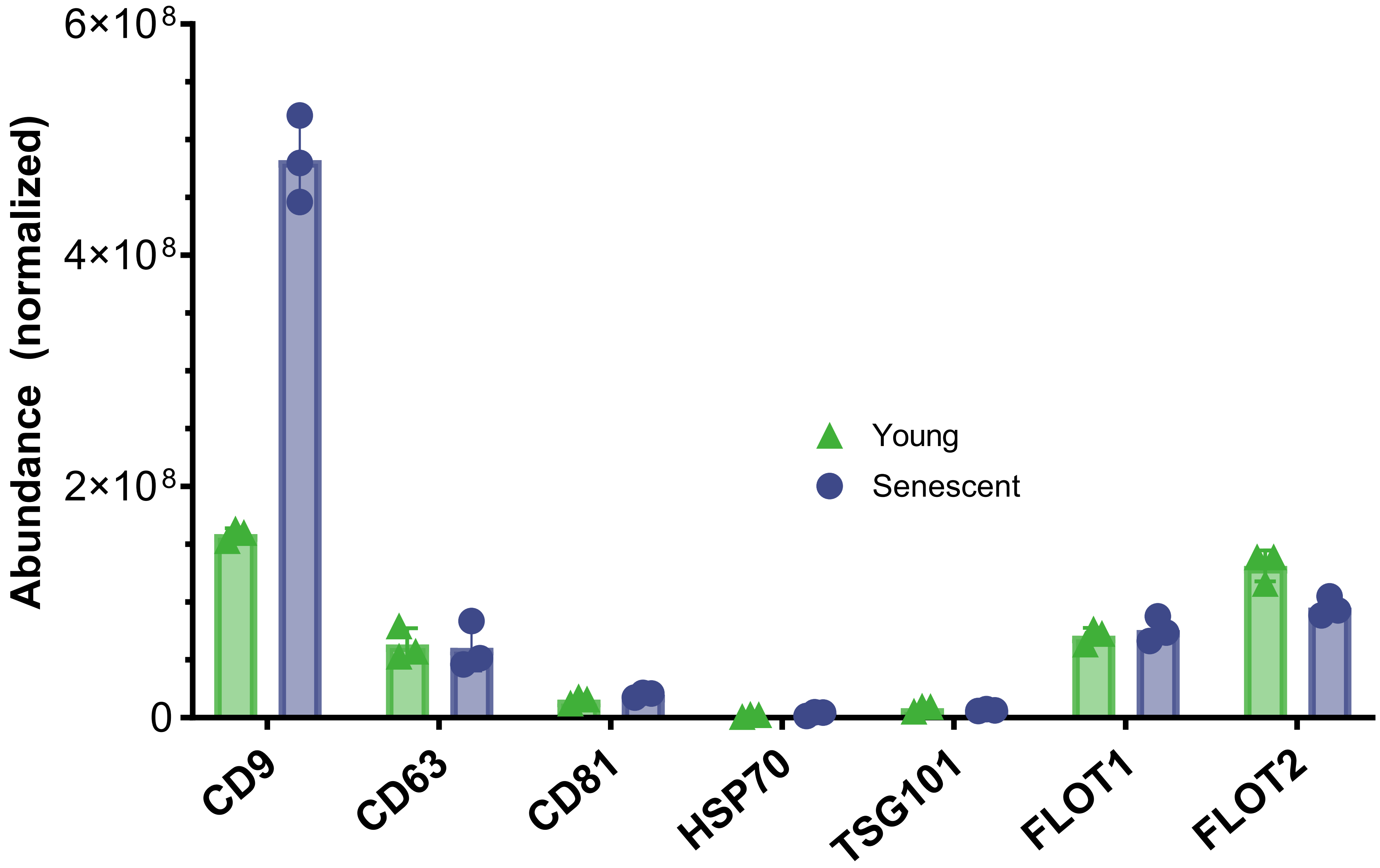

**Supplementary Figure S5b: Characterization of the extracellular vesicles (IV).** Measurement of CD9+, CD63+, and CD81+ and other markers by LC-MS/MS. (A) Functional analysis for cellular component (gene ontology). (B) Venn diagrams comparing proteins identified in extracellular vesicles with those included in the Vesiclopedia compendium.

#### (A) Cellular component (Gene Ontology)

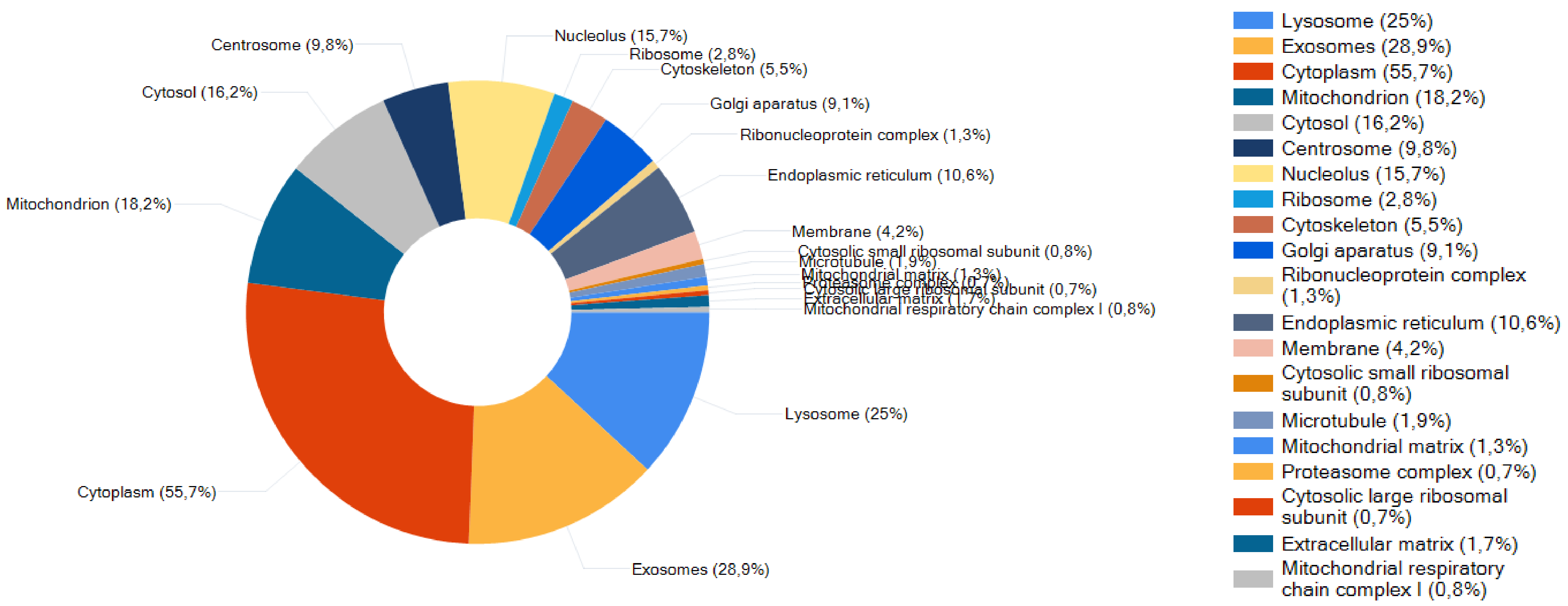


#### (B) Comparison with Vesiclepedia data

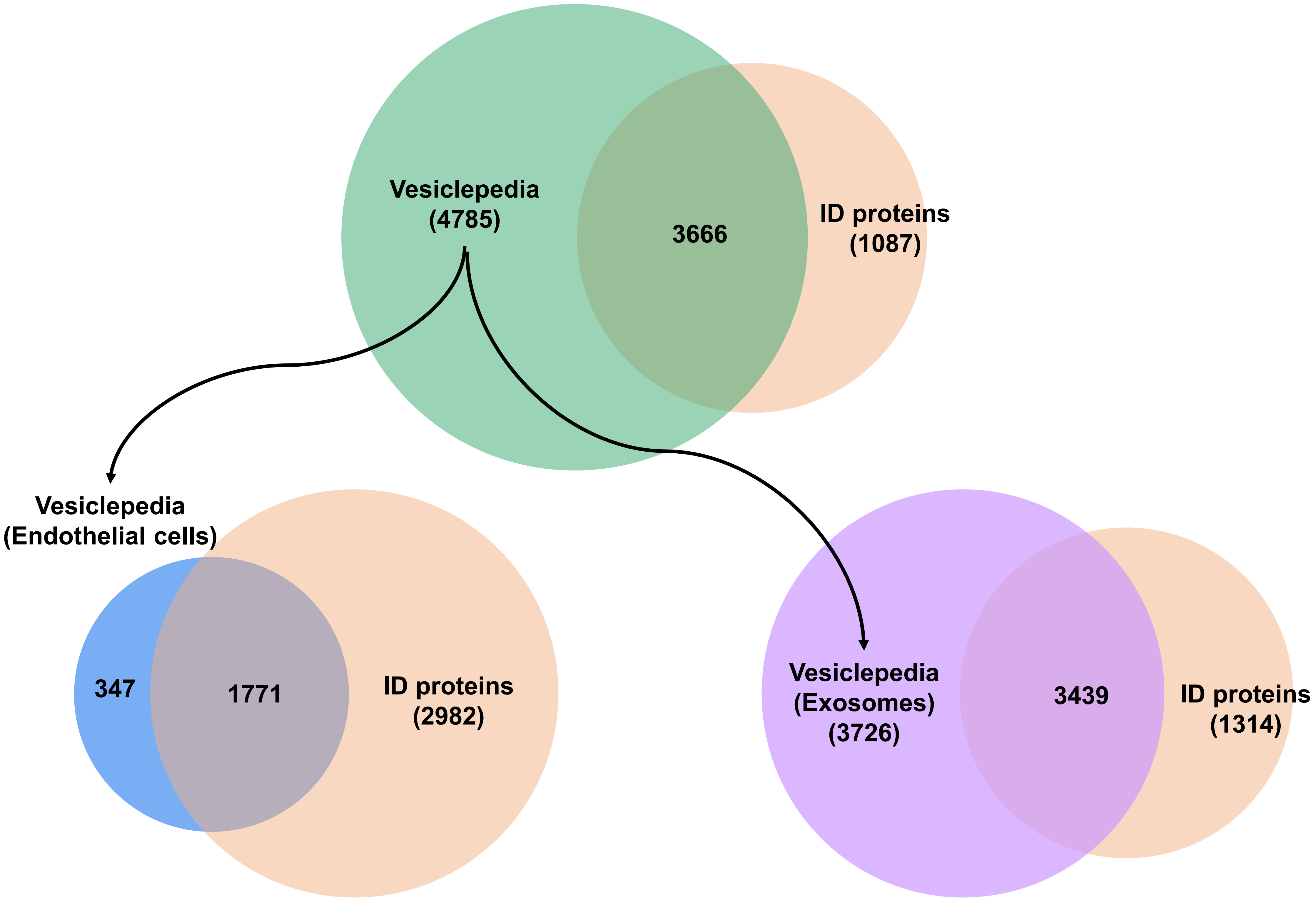

**Supplementary Figure S6a: Characterization of the extracellular vesicles (IV) by WB.**
Analysis of common EV markers CD9+, CD63+, and Tumor susceptibility gene 101 protein (TSG101) by western blot.

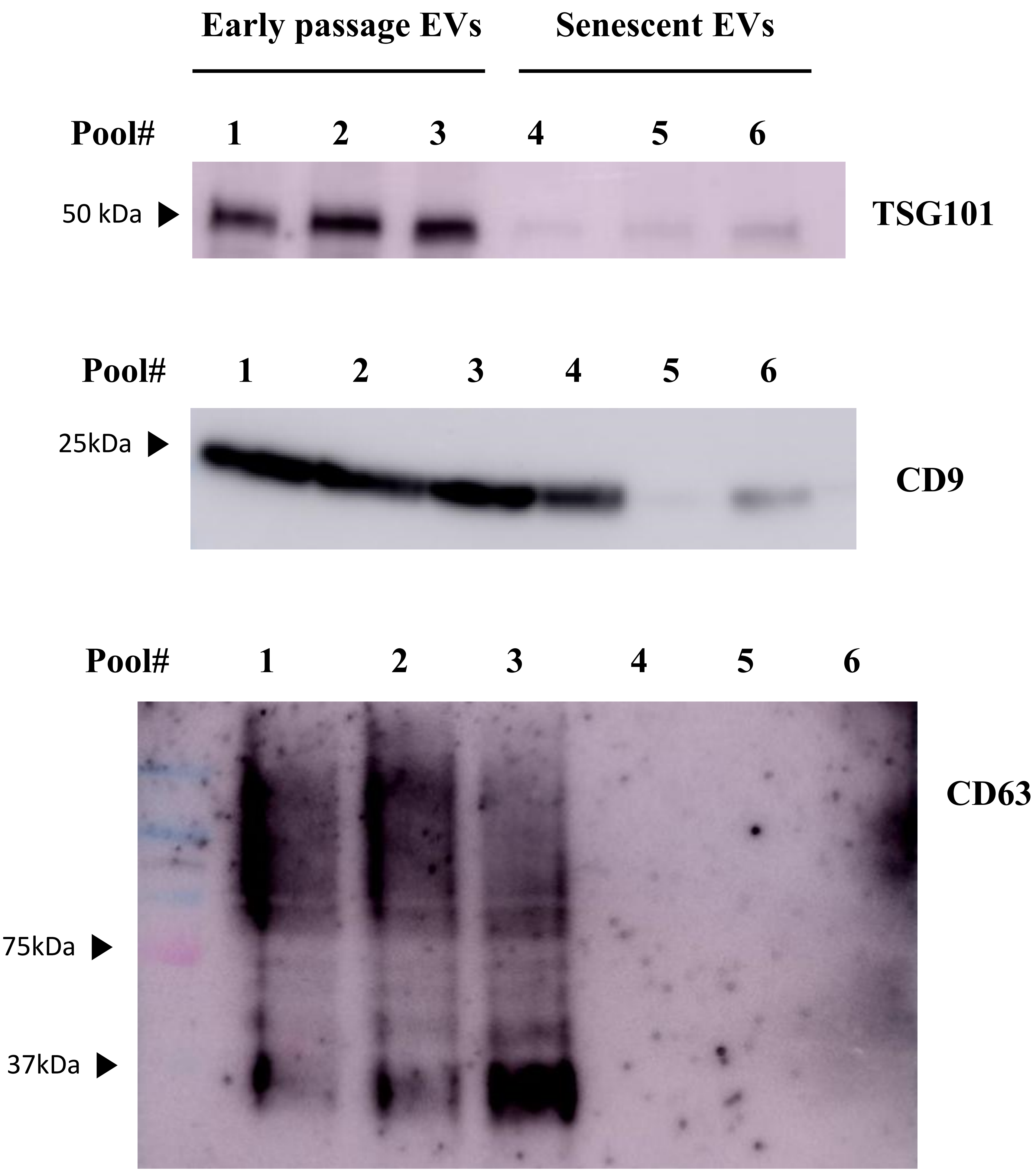

**Supplementary Figure S7a: Senescent cells and EVs characterization by proteomics analysis.** (A) Venn and (B) Circos diagrams comparing the proteins identified and found differentially expressed in senescent cells and extracellular vesicles (EVs).

#### A) Venn Diagrams

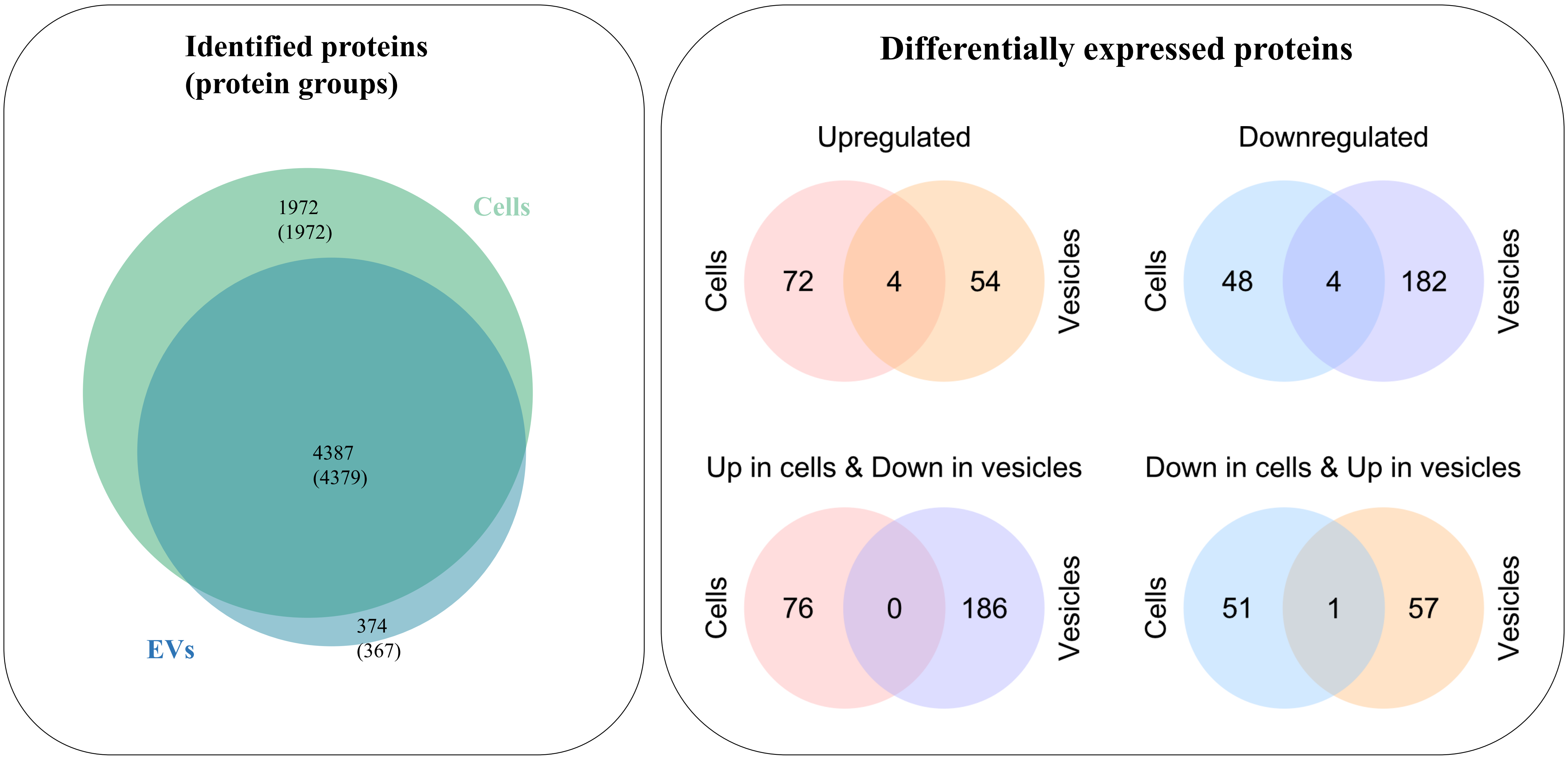


#### B) Circos Diagrams

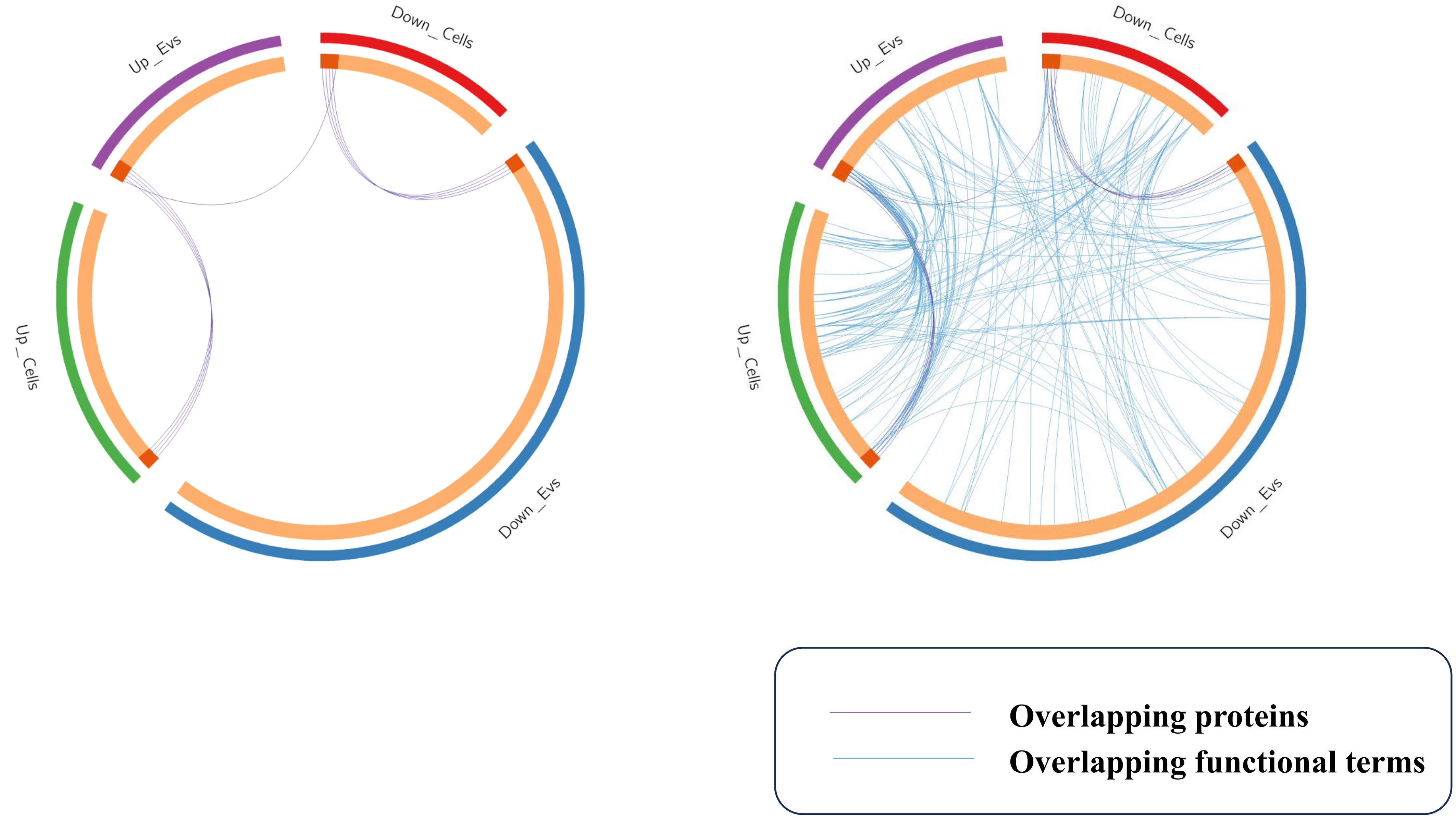

**Supplementary Figure S7b: Senescent cells and EVs characterization by proteomics analysis.** Functional enrichment analysis of proteins found A) over-expressed and B) down-expressed in senescent compared to early passage HUVECs

#### A) Over-expressed proteins (Bubble plot and treemap)

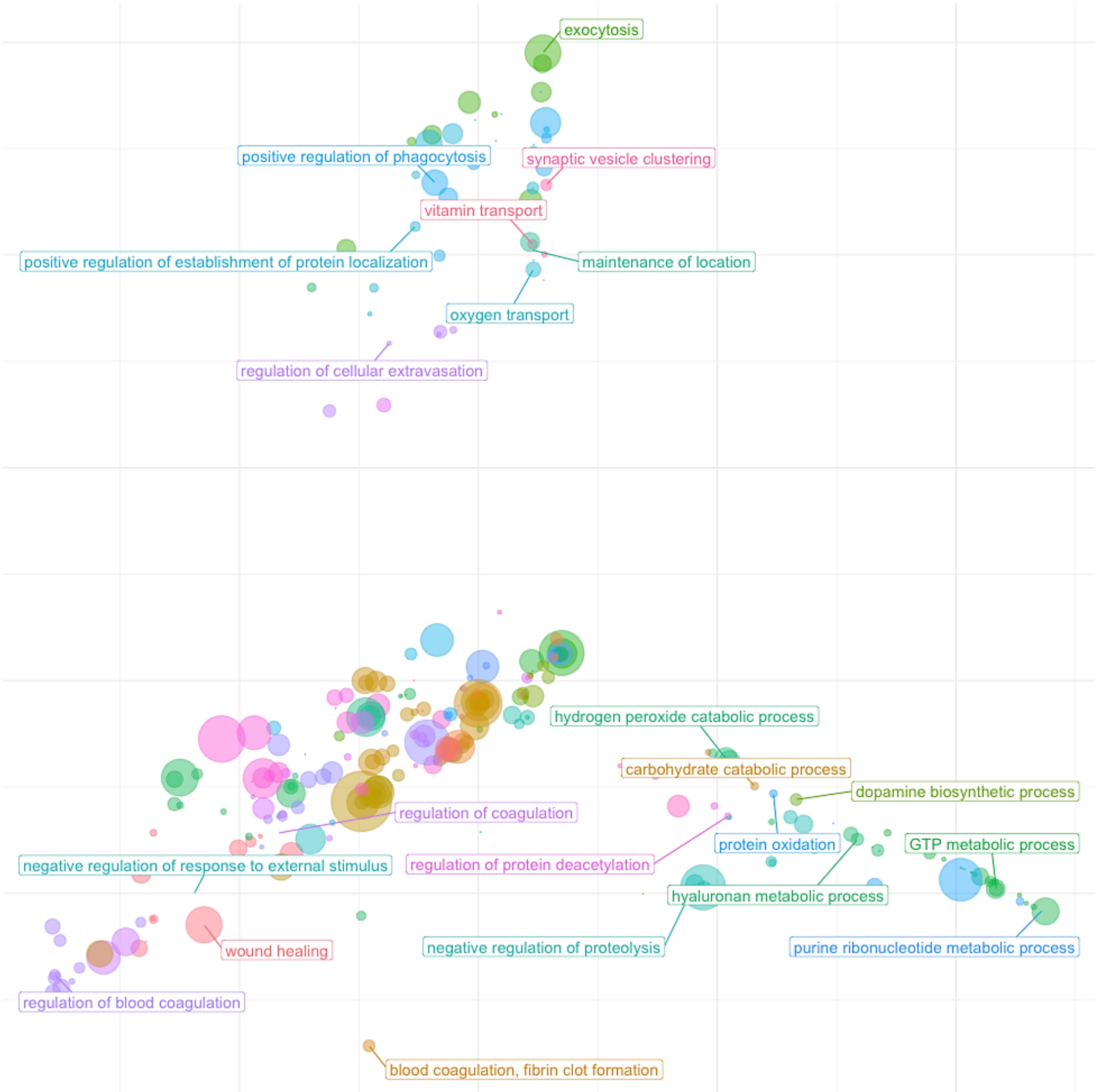


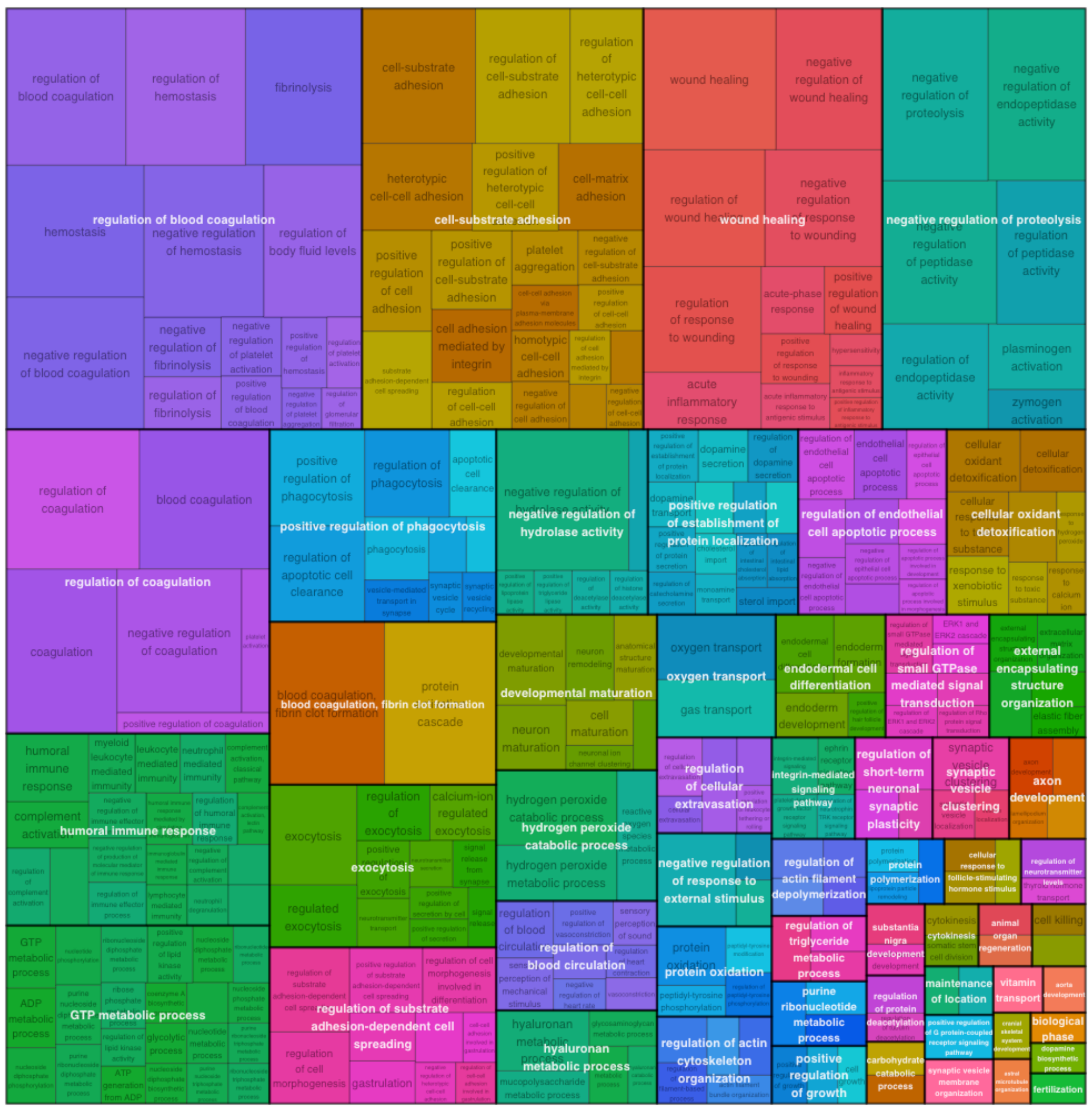


#### B) Down-expressed proteins (Bubble plot and treemap)

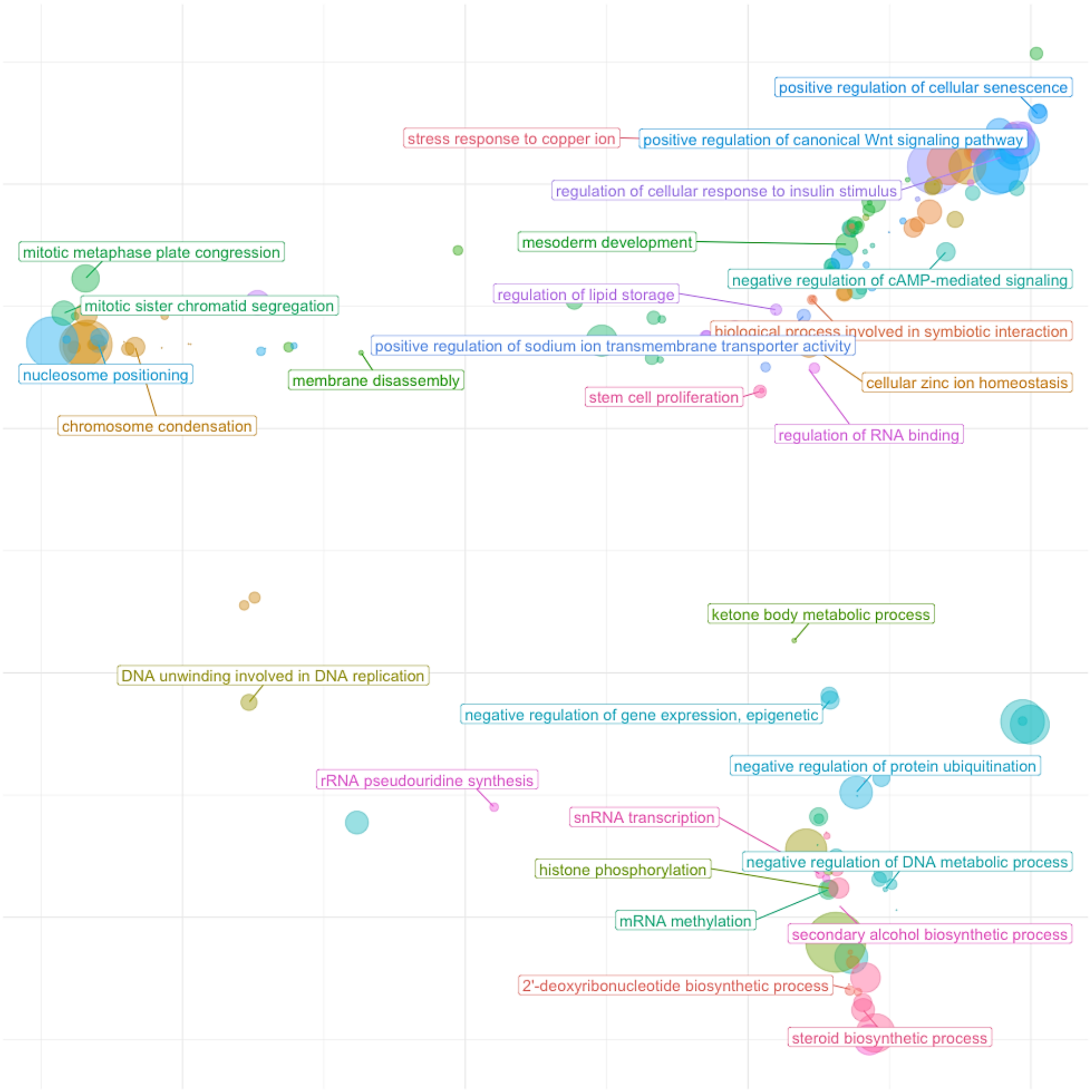


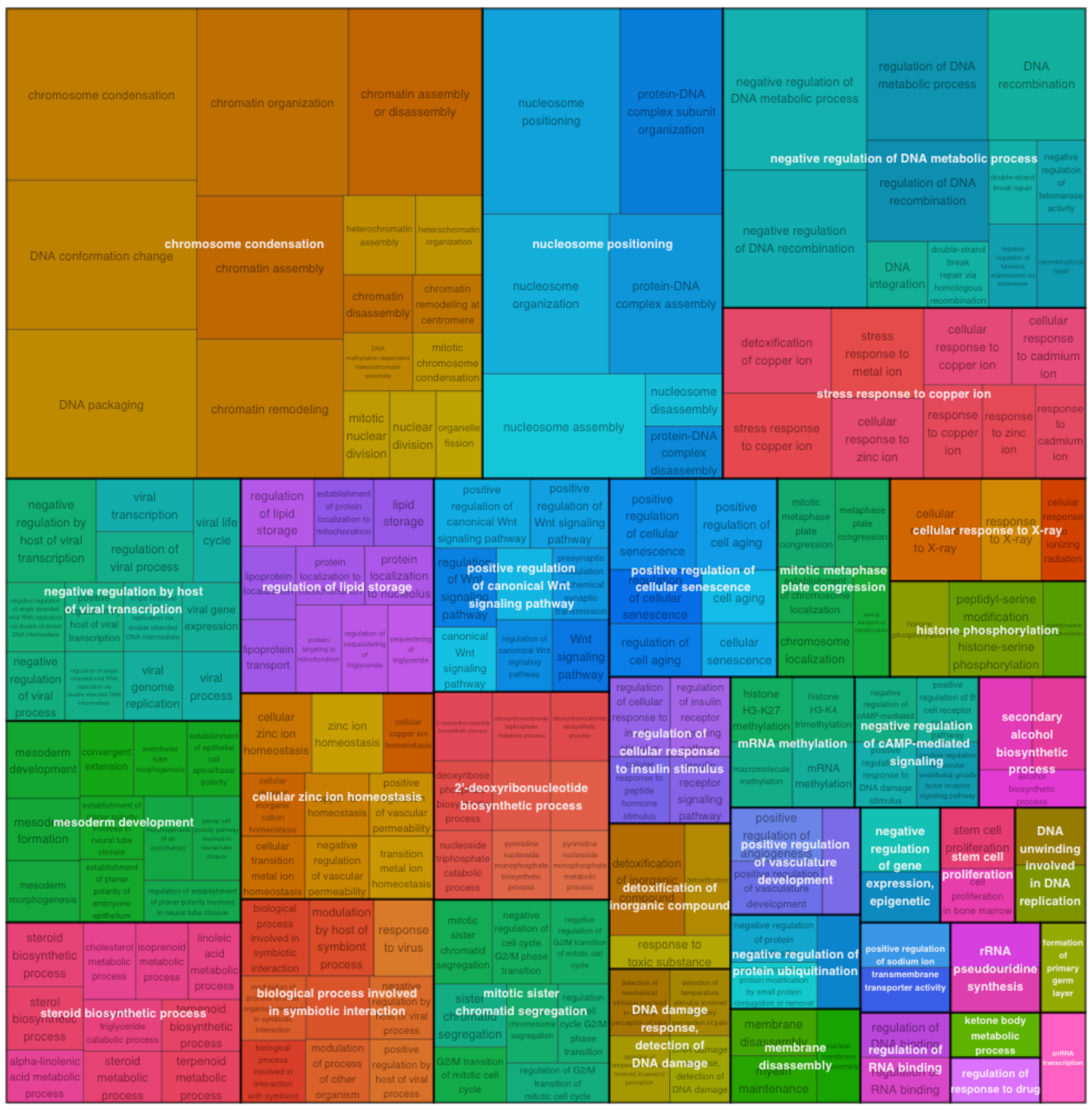

**Supplementary Figure S7c: Senescent cells and EVs characterization by proteomics analysis.** Functional enrichment analysis of proteins found A) over-expressed and B) down-expressed in senescent compared to early passage extracellular vesicles

#### A) Over-expressed proteins (Bubble plot and treemap)

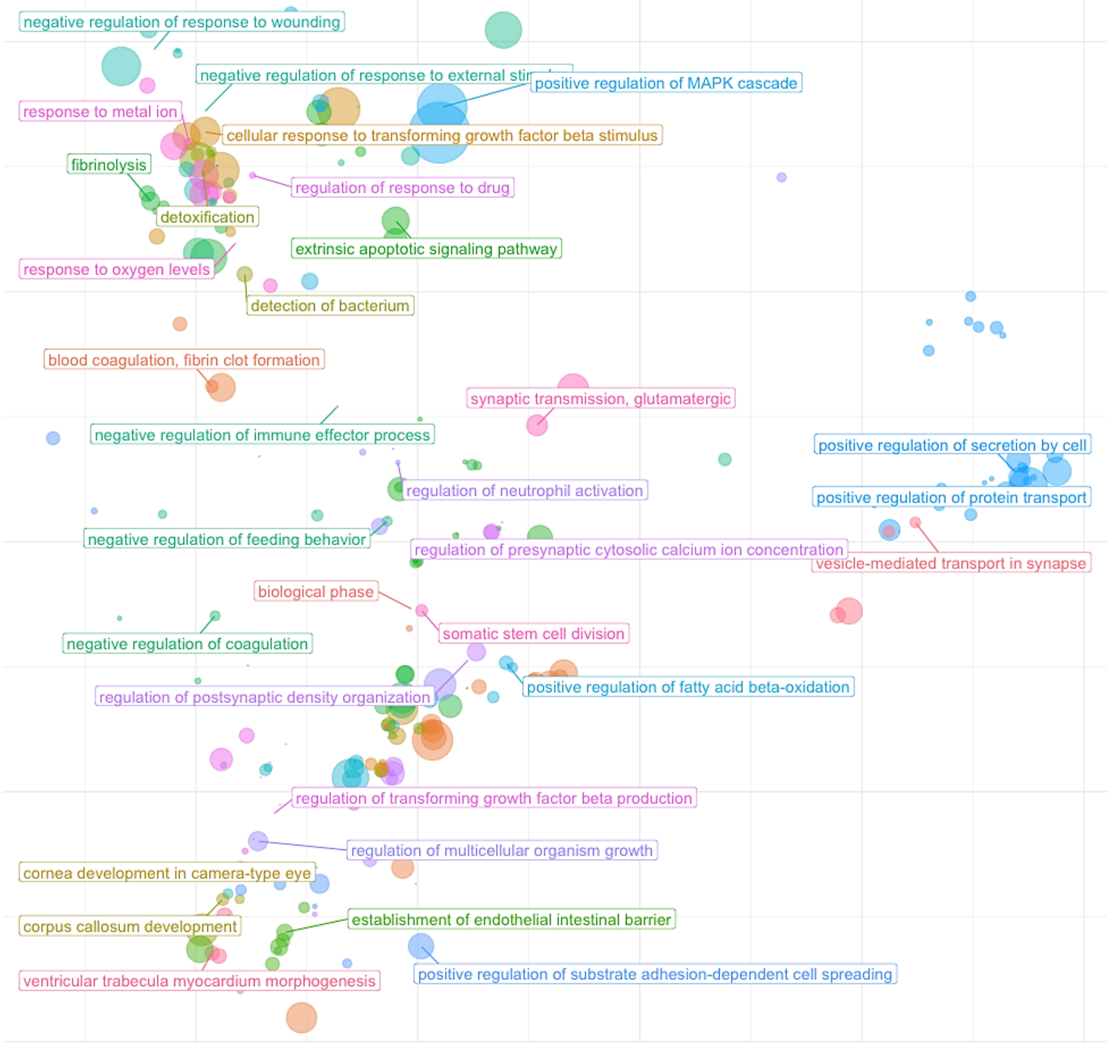


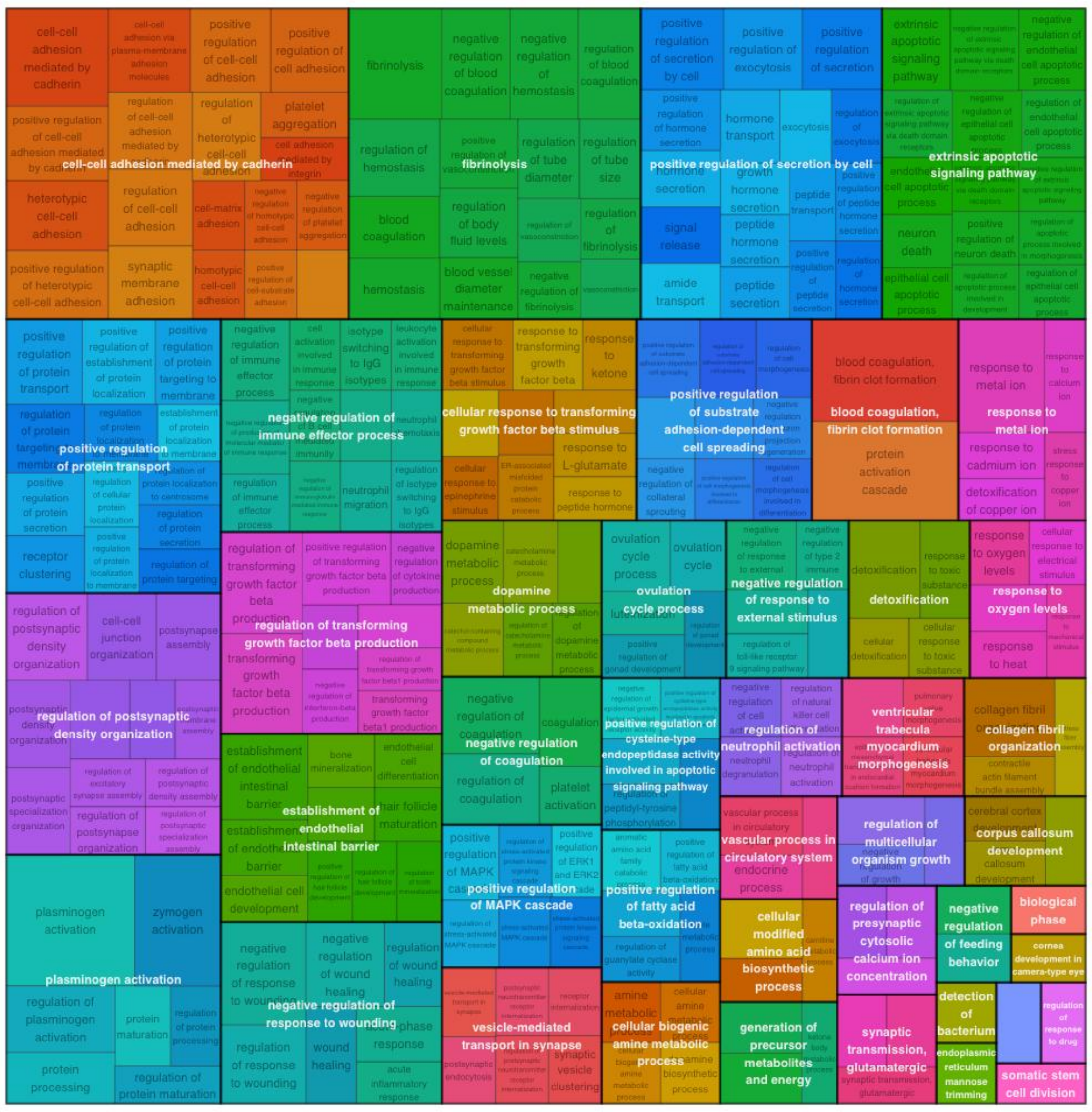


#### B) Down-expressed proteins (Bubble plot and treemap)

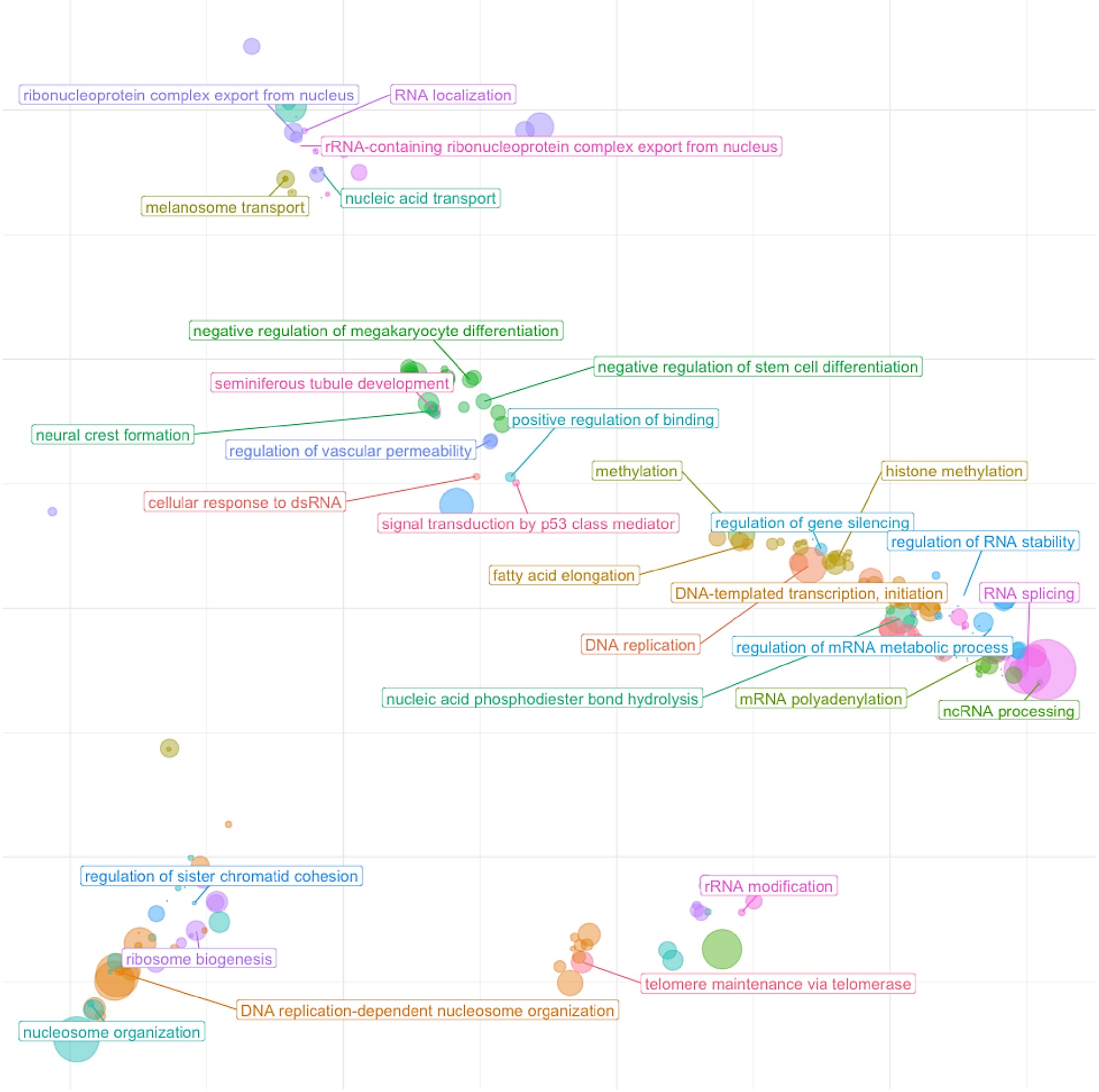


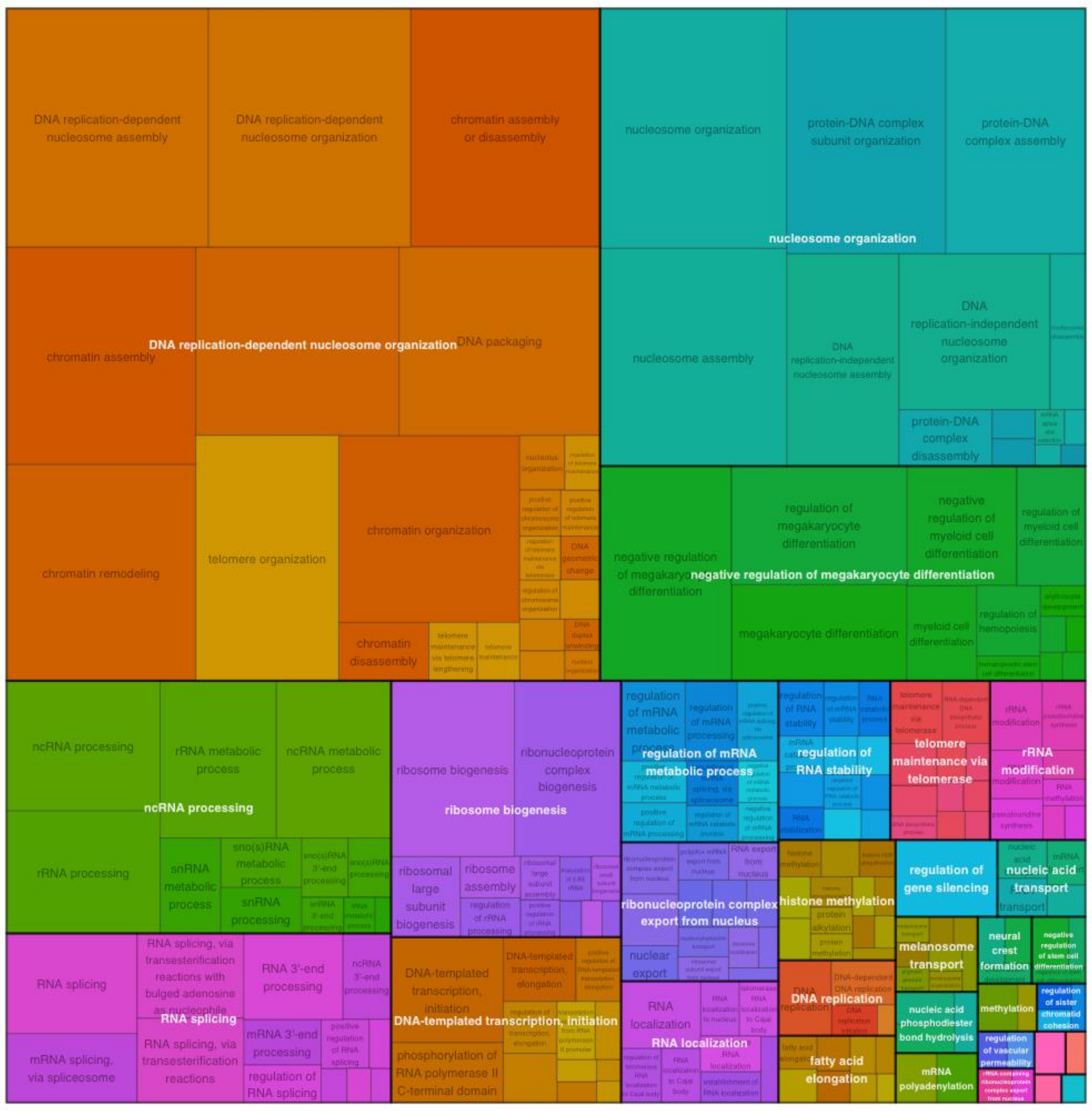

**Supplementary Figure S8a: Senescent cells and EVs characterization by transcriptomics analysis.** Functional enrichment analysis of mRNA found A) up-regulated and B) down-regulated in senescent compared to early passage HUVECs

#### A) Up-regulated mRNA
#### (Bubble plot and treemap)

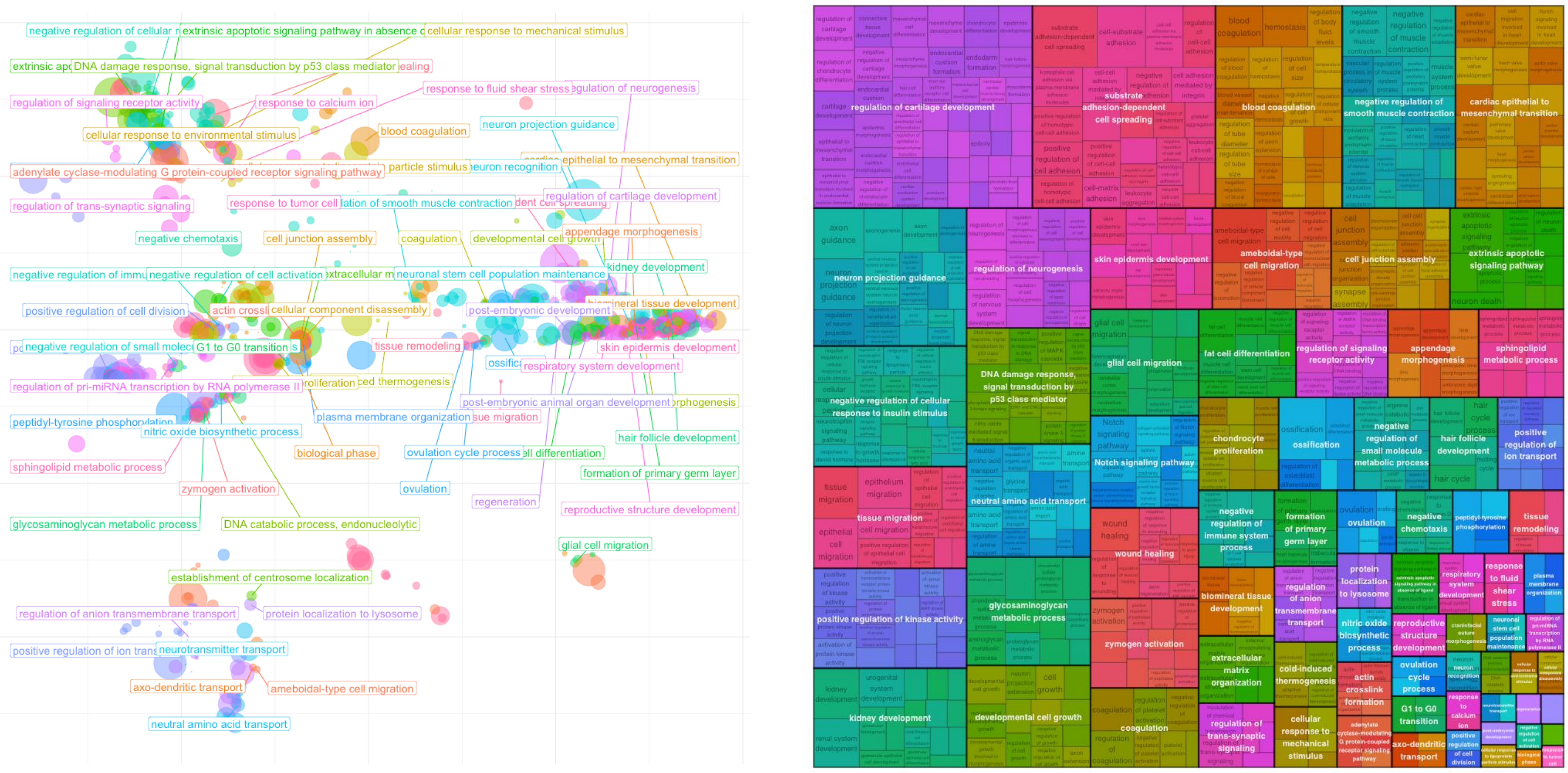


#### B) Down-regulated mRNA
#### (Bubble plot and treemap)

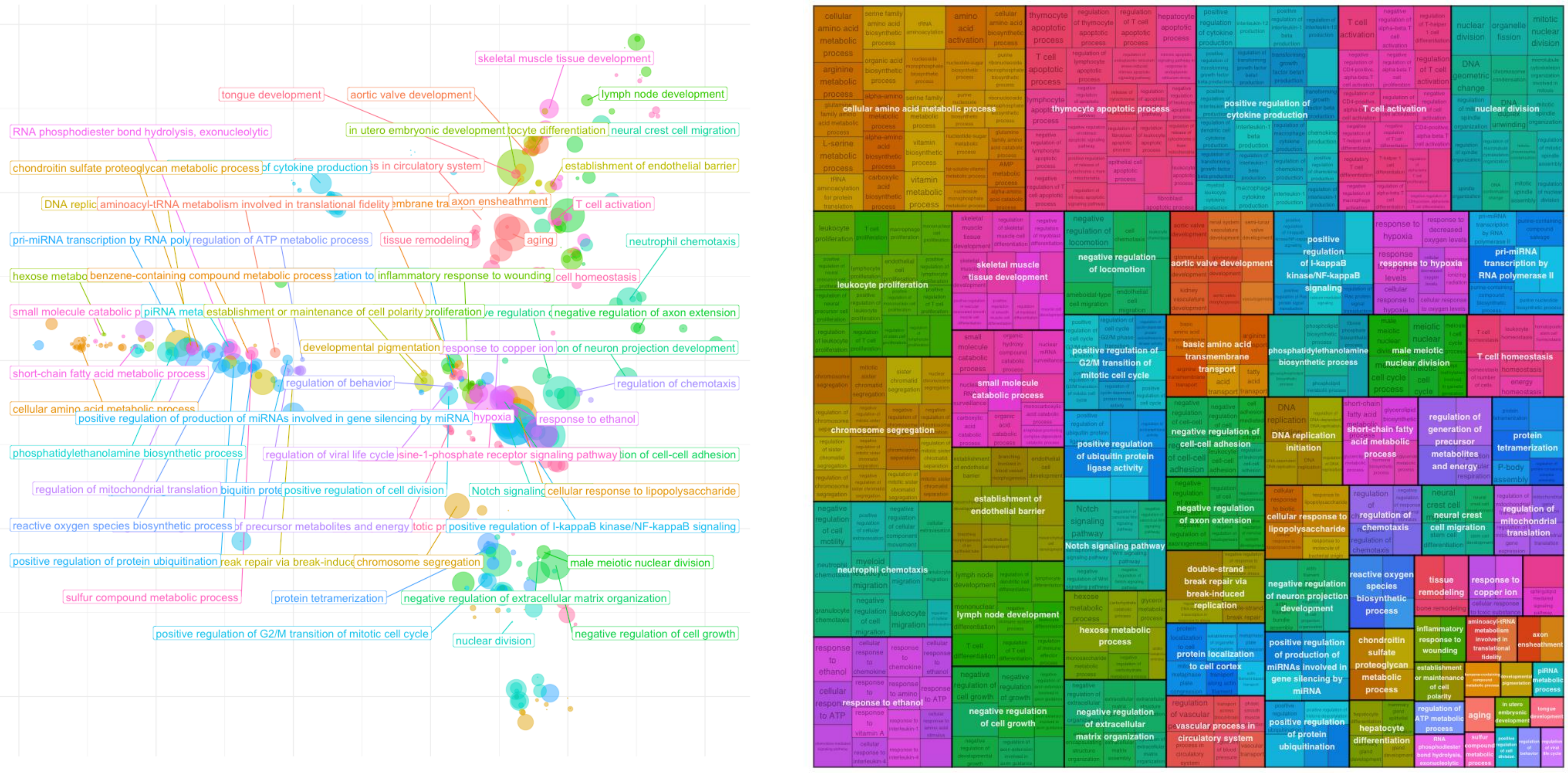

**Supplementary Figure S8b: Functional enrichment analysis of mRNA.** (A) Functional cluster and functional enrichment analysis of mRNA found E) up-regulated and B-D) down-regulated in senescent compared to early passage extracellular vesicles

**A) Functional cluster (Metascape)**

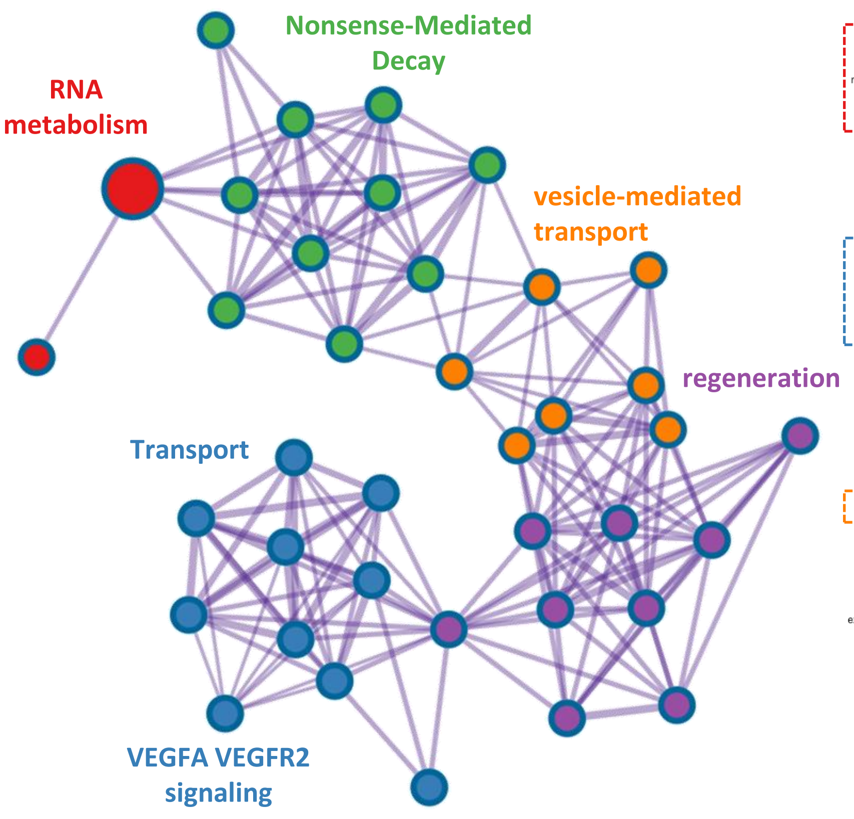


**B) Down-regulated mRNA (GO_Biological process)**

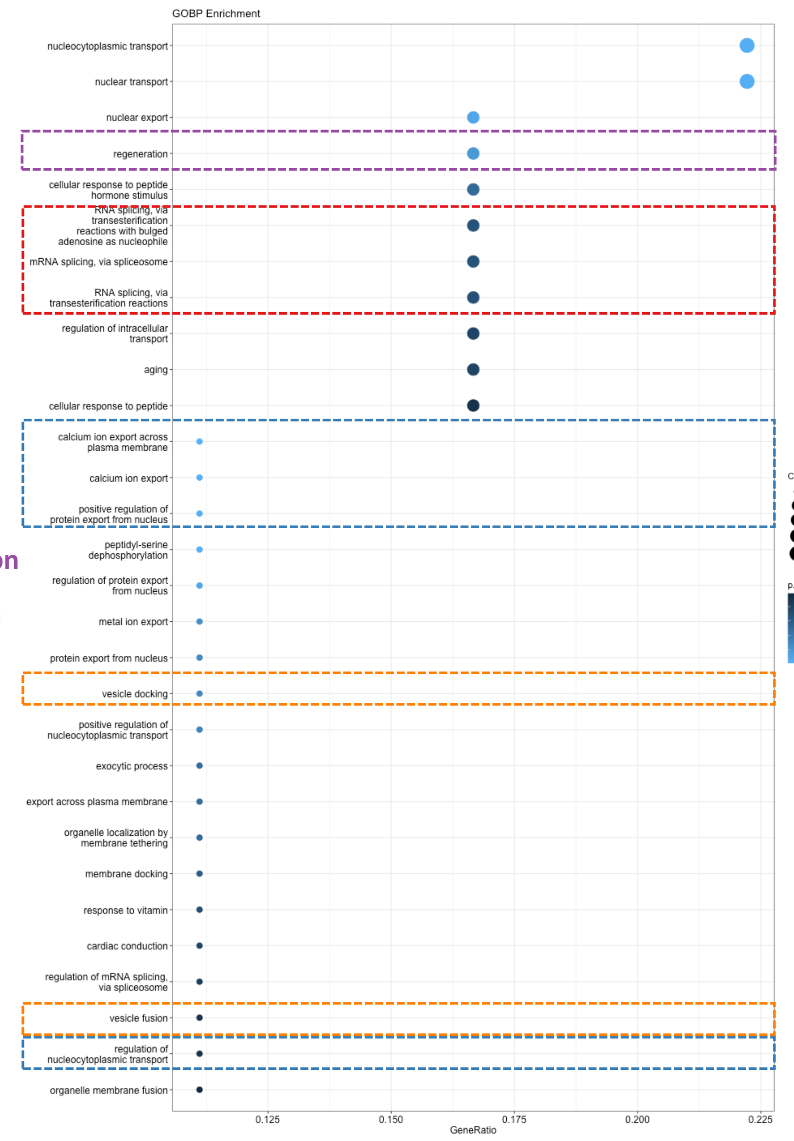


**C) Down-regulated mRNA (Reactome)**

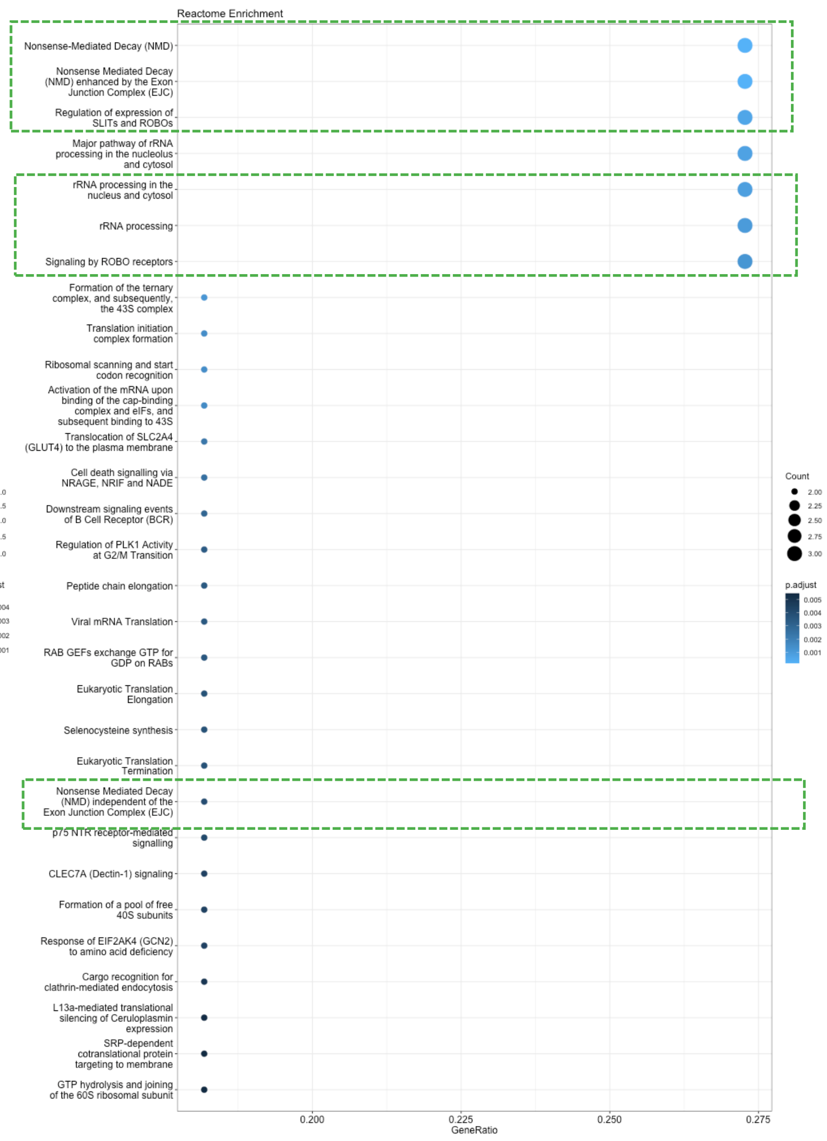


**D) Down-regulated mRNA (Bubble plot)**

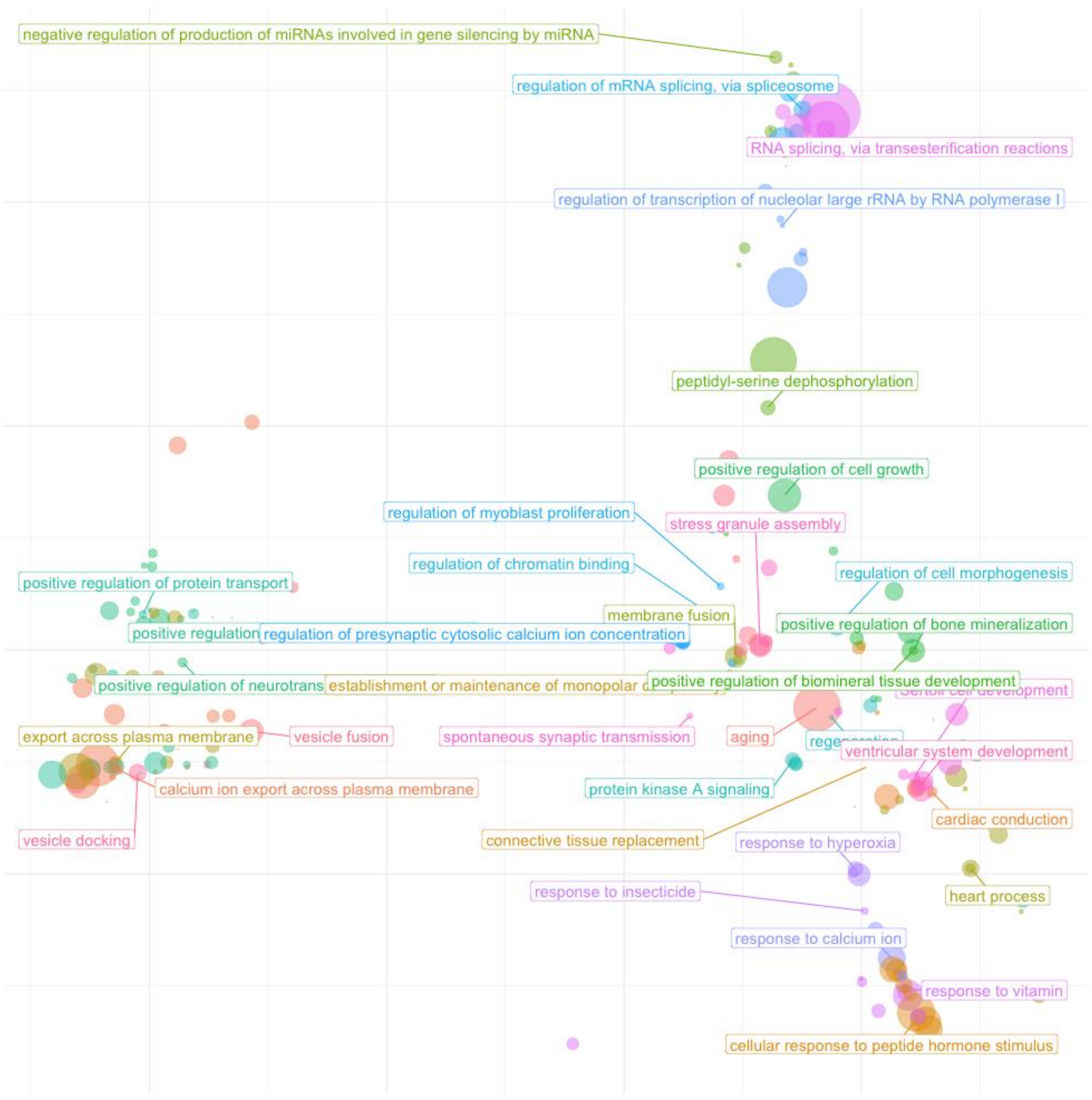


**E) Up-regulated mRNA (Bubble plot)**

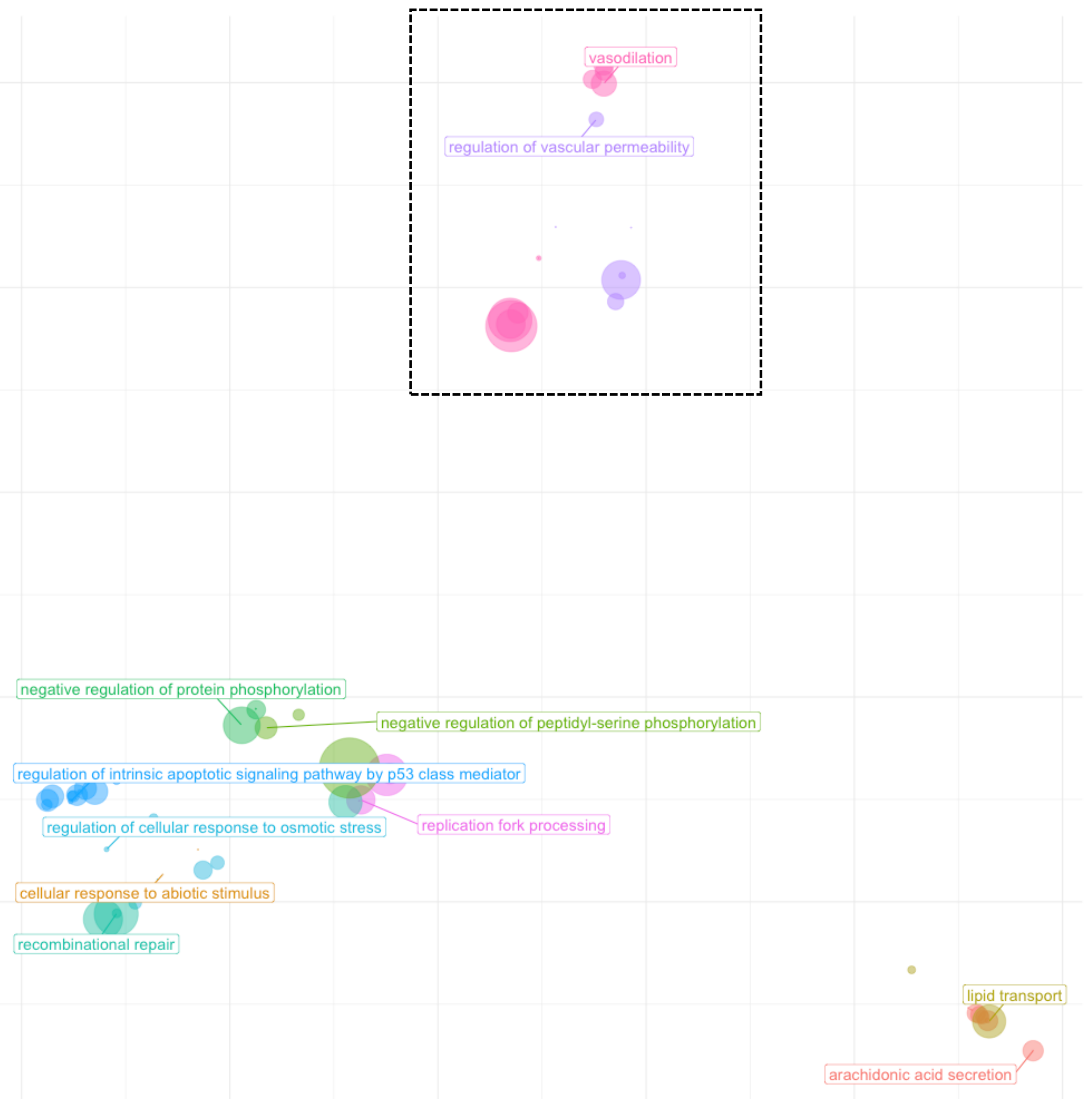

#### Supplementary Figure S8c: Functional integration of proteomics and transcriptomics analysis of senescent extracellular vesicles: functional reduction heatmaps

##### A) Up-regulated mRNA/Protein

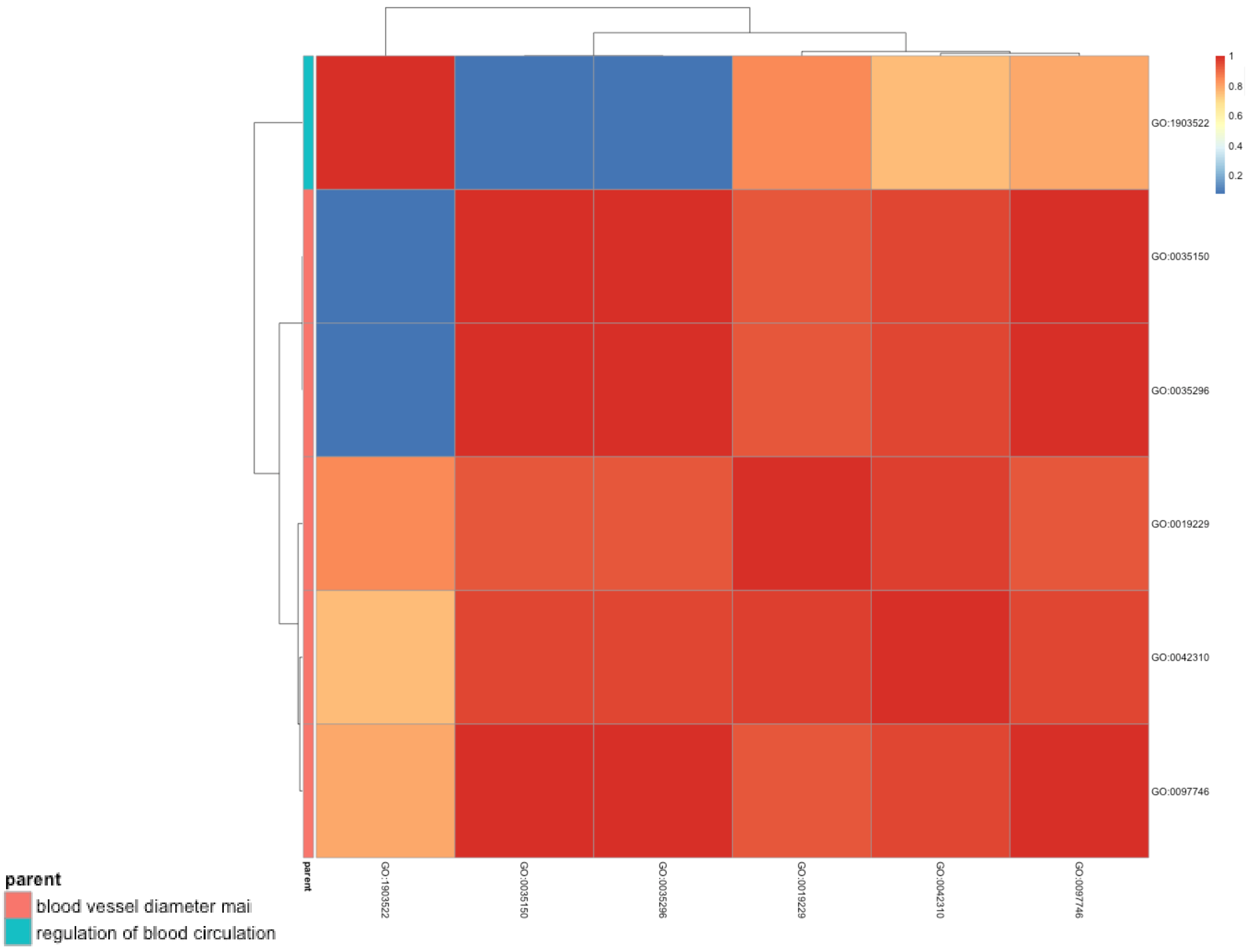


##### B) Down-regulated mRNA/Protein

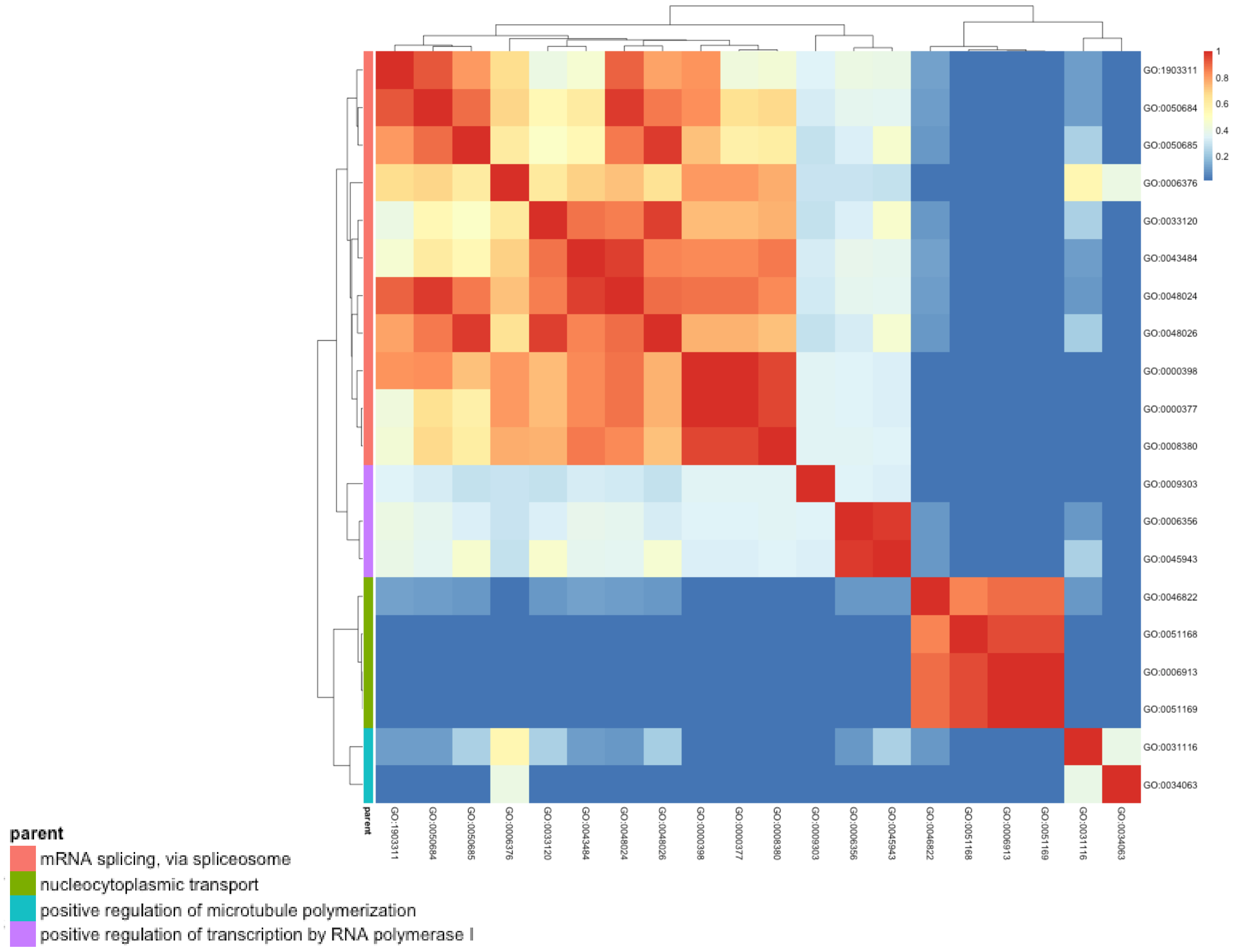

**Supplementary Figure S9a: Functional enrichment analysis of miRNA.** A) Up-regulated and B) down-regulated miRNA in senescent compared to early passage cells.

#### A) Up-regulated miRNA (Bubble plot)

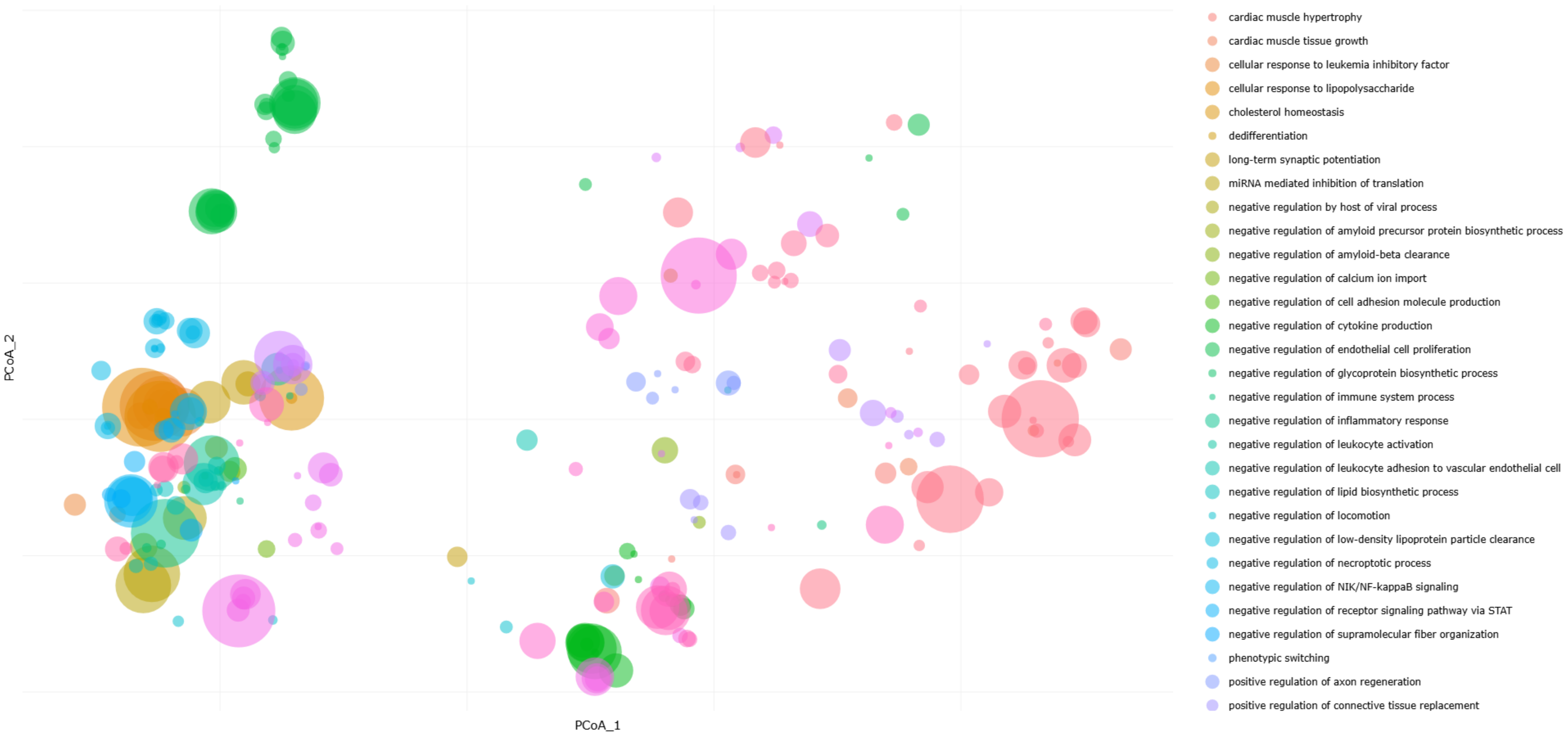


#### B) Down-regulated miRNA (Bubble plot)

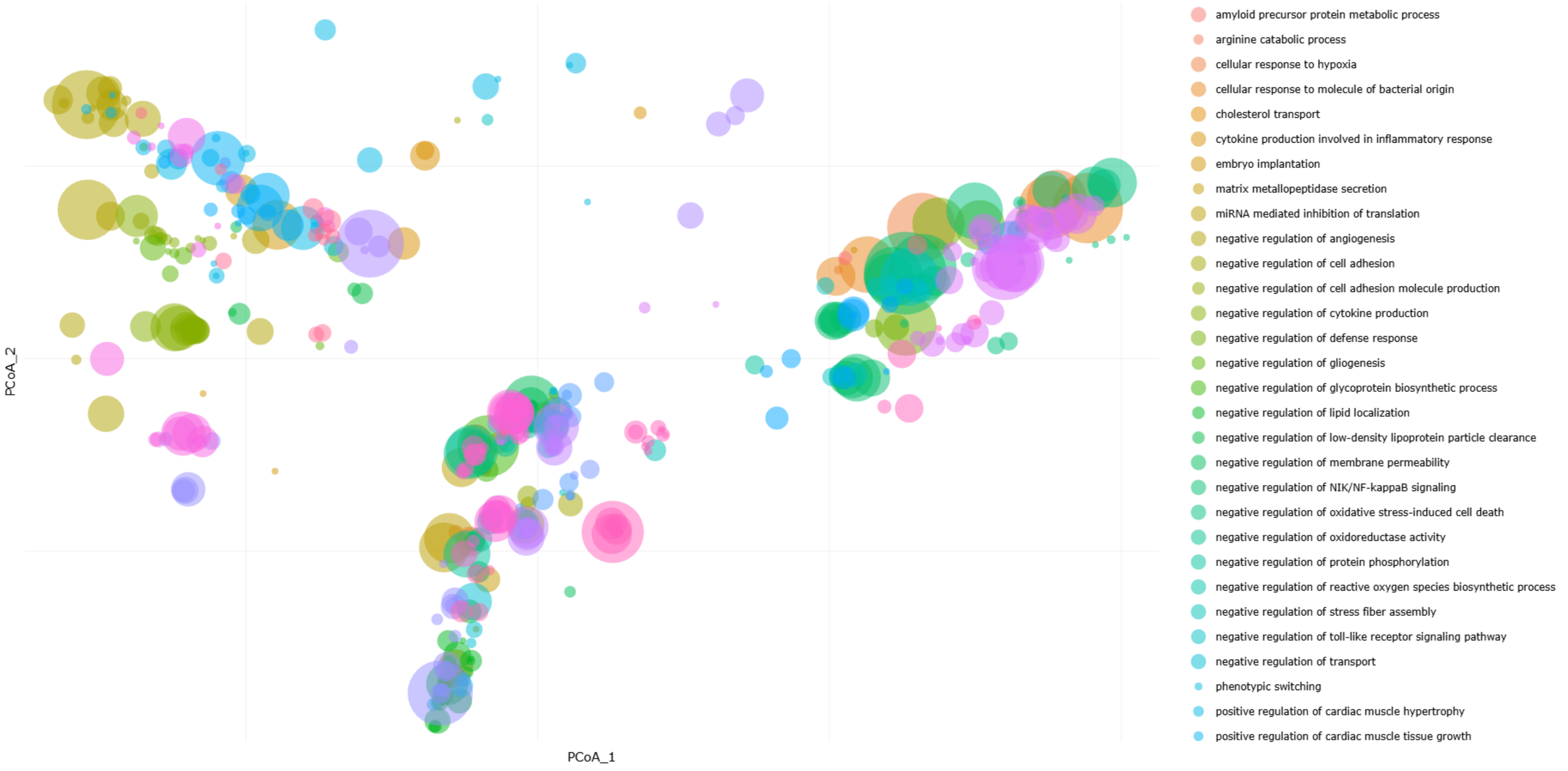

**Supplementary Figure S9b: Functional enrichment analysis of miRNA.** A) Up-regulated and B) down-regulated miRNA in senescent compared to early passage extracellular vesicles.

#### A) Up-regulated miRNA (Bubble plot)

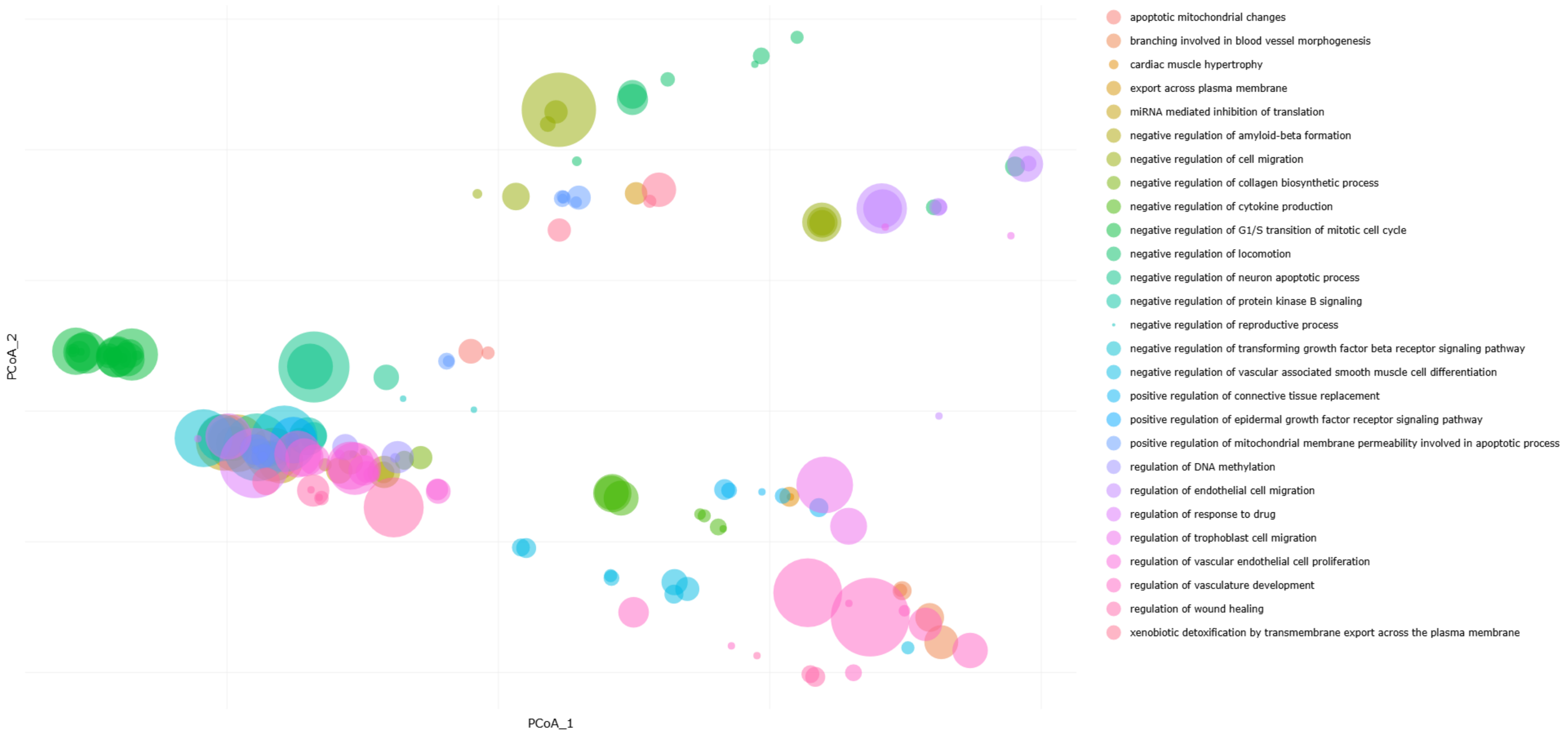


#### B) Down-regulated miRNA (Bubble plot)

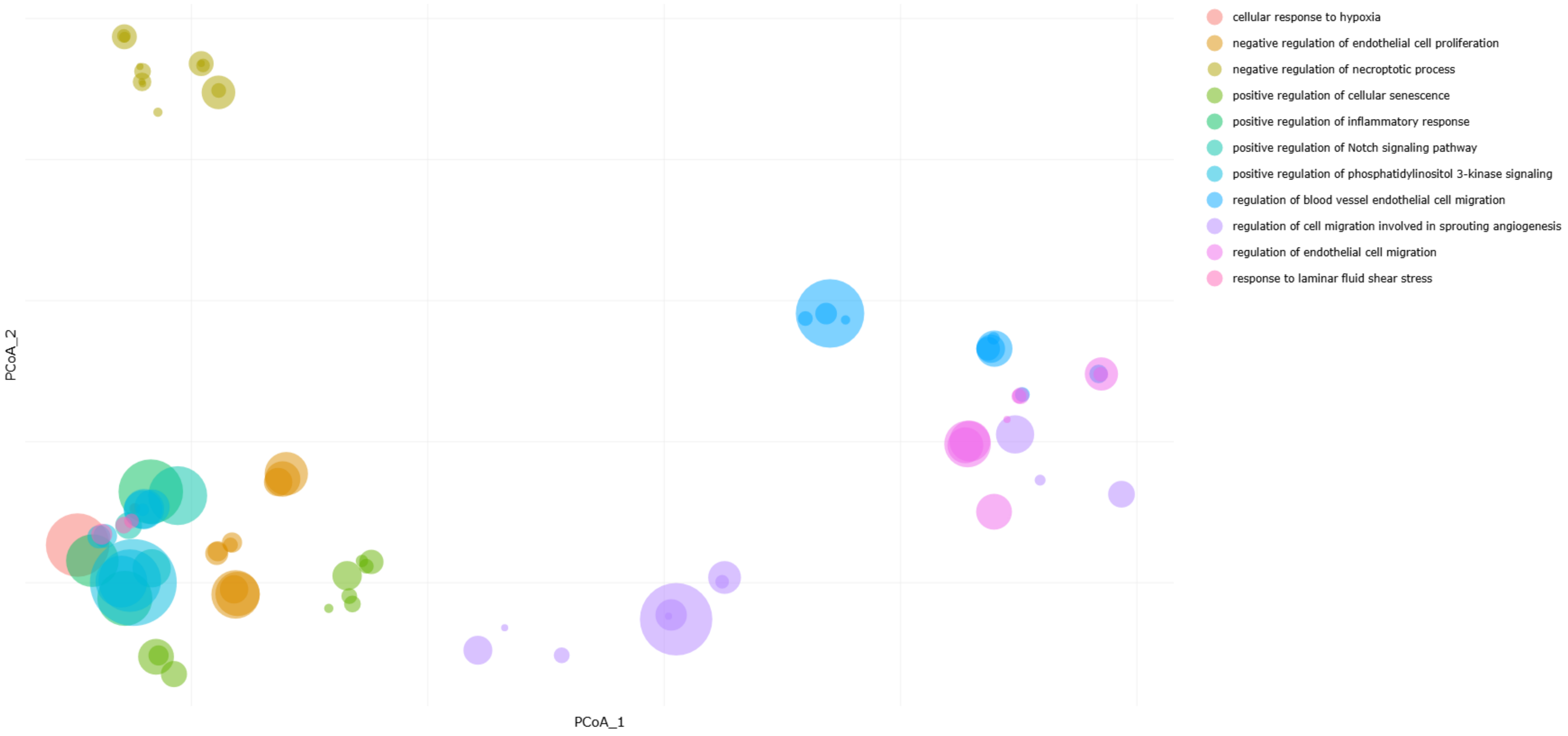

#### Supplementary Figure S10a: Functional integration of miRNAomics with proteomics and transcriptomics analysis of senescent cells: functional reduction heatmaps

##### A) Up-regulated miRNA/mRNA

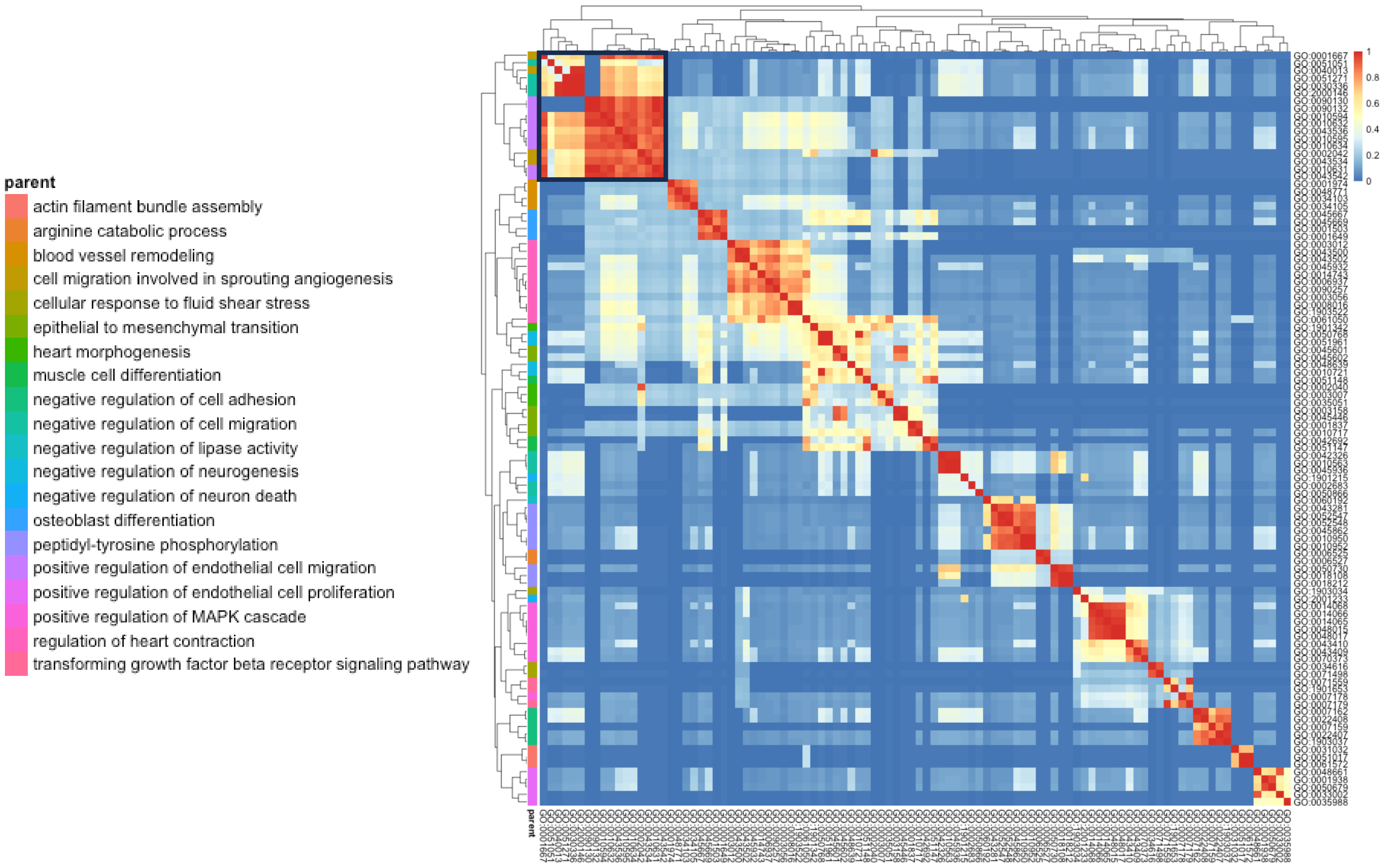


##### B) Up-regulated miRNA/mRNA/Protein

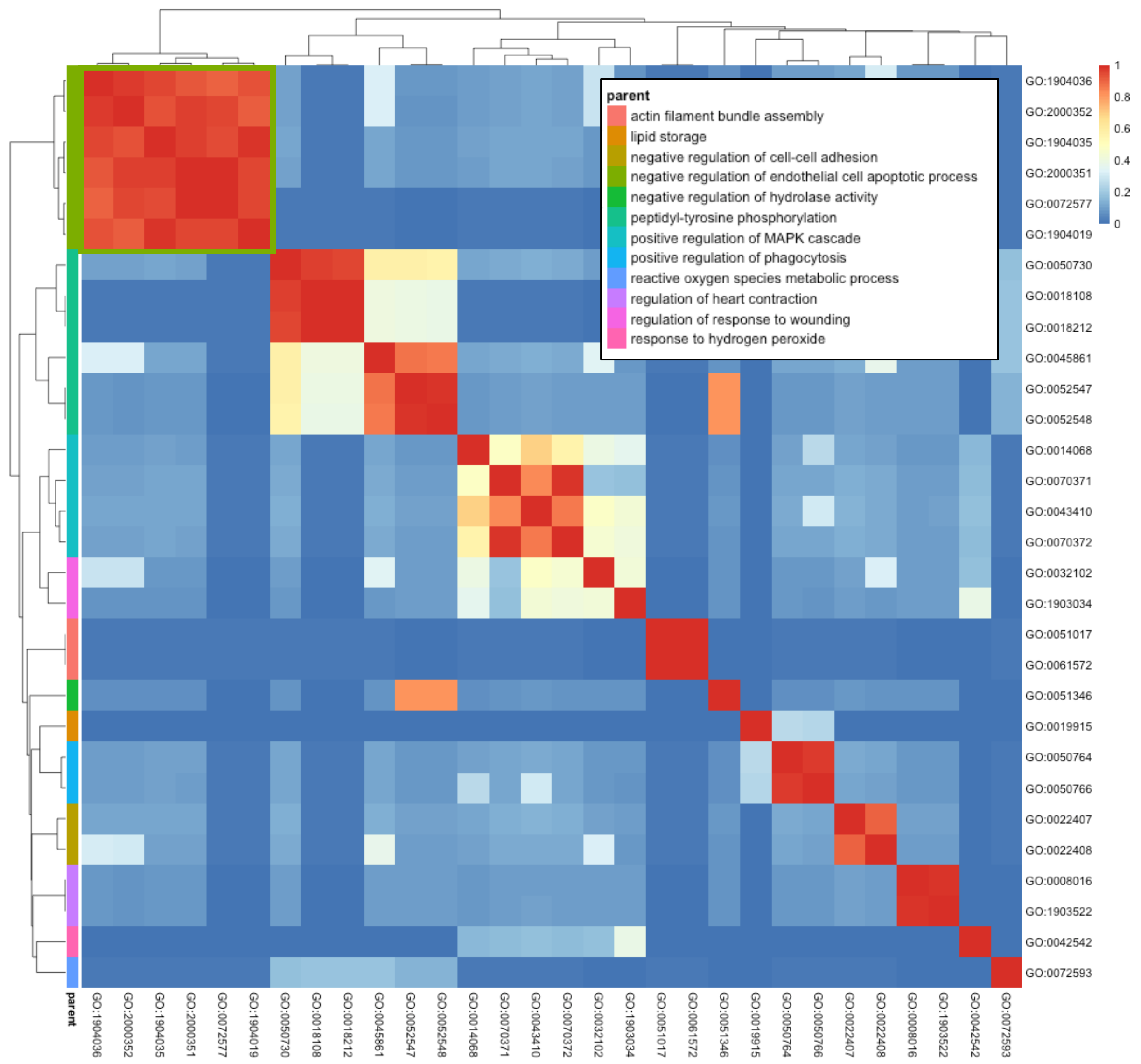


##### Up-regulated miRNA/mRNA/Protein

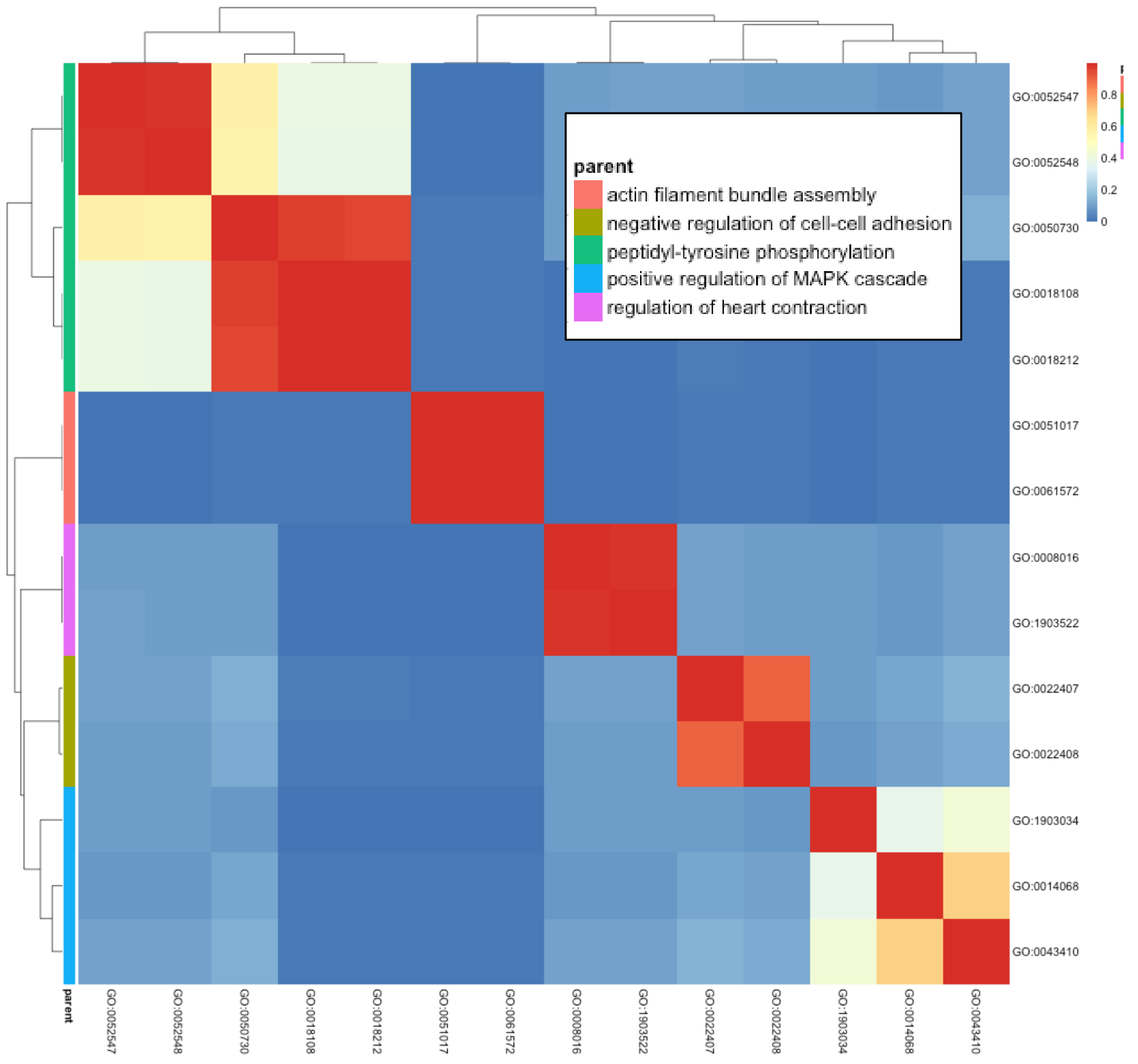

**Supplementary Figure S10b: Functional integration of miRNAomics with proteomics and transcriptomics analysis of senescent cells: functional reduction heatmaps**

#### A) Down-regulated miRNA/mRNA

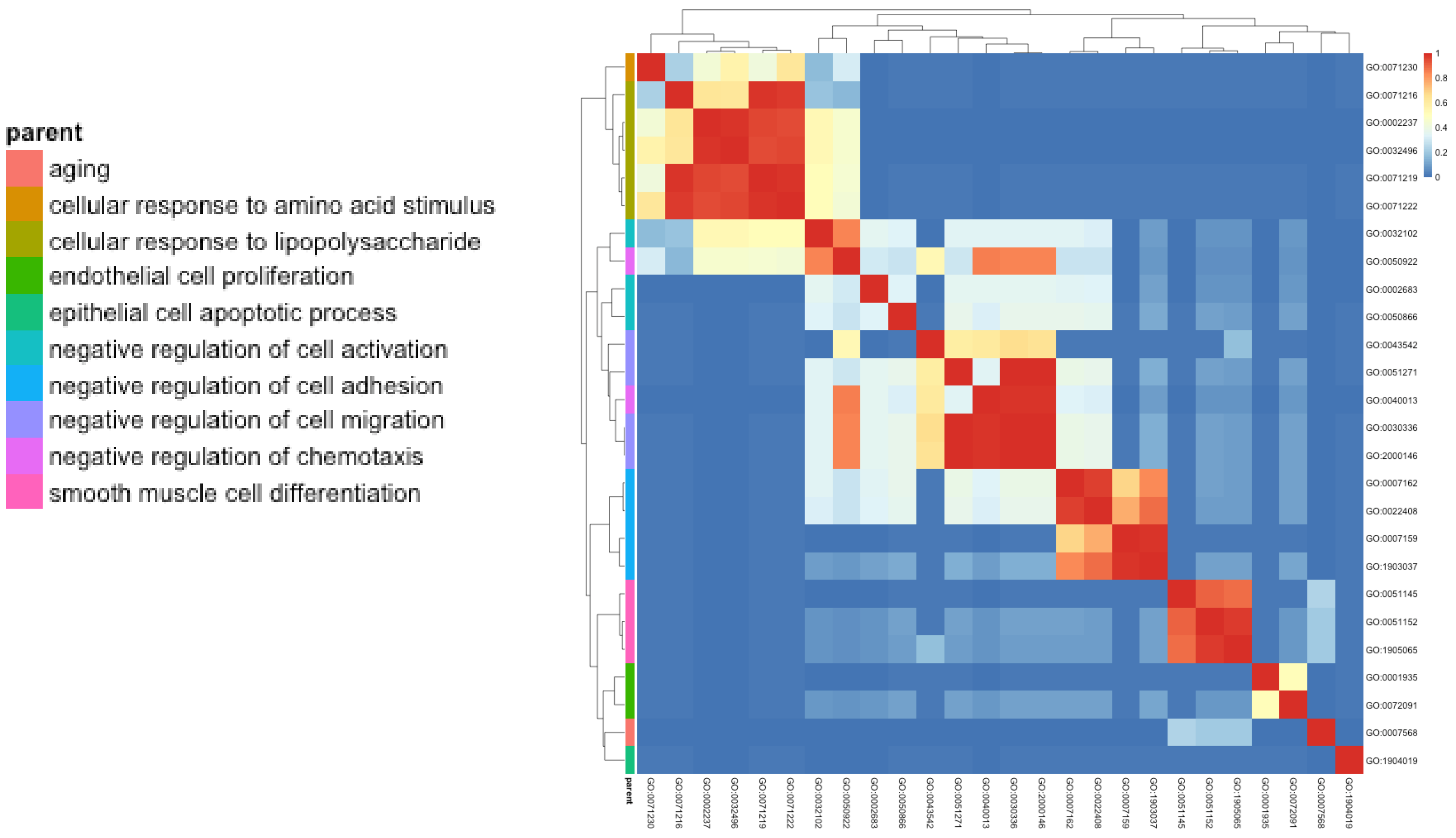


#### B) Down-regulated mRNA/Protein

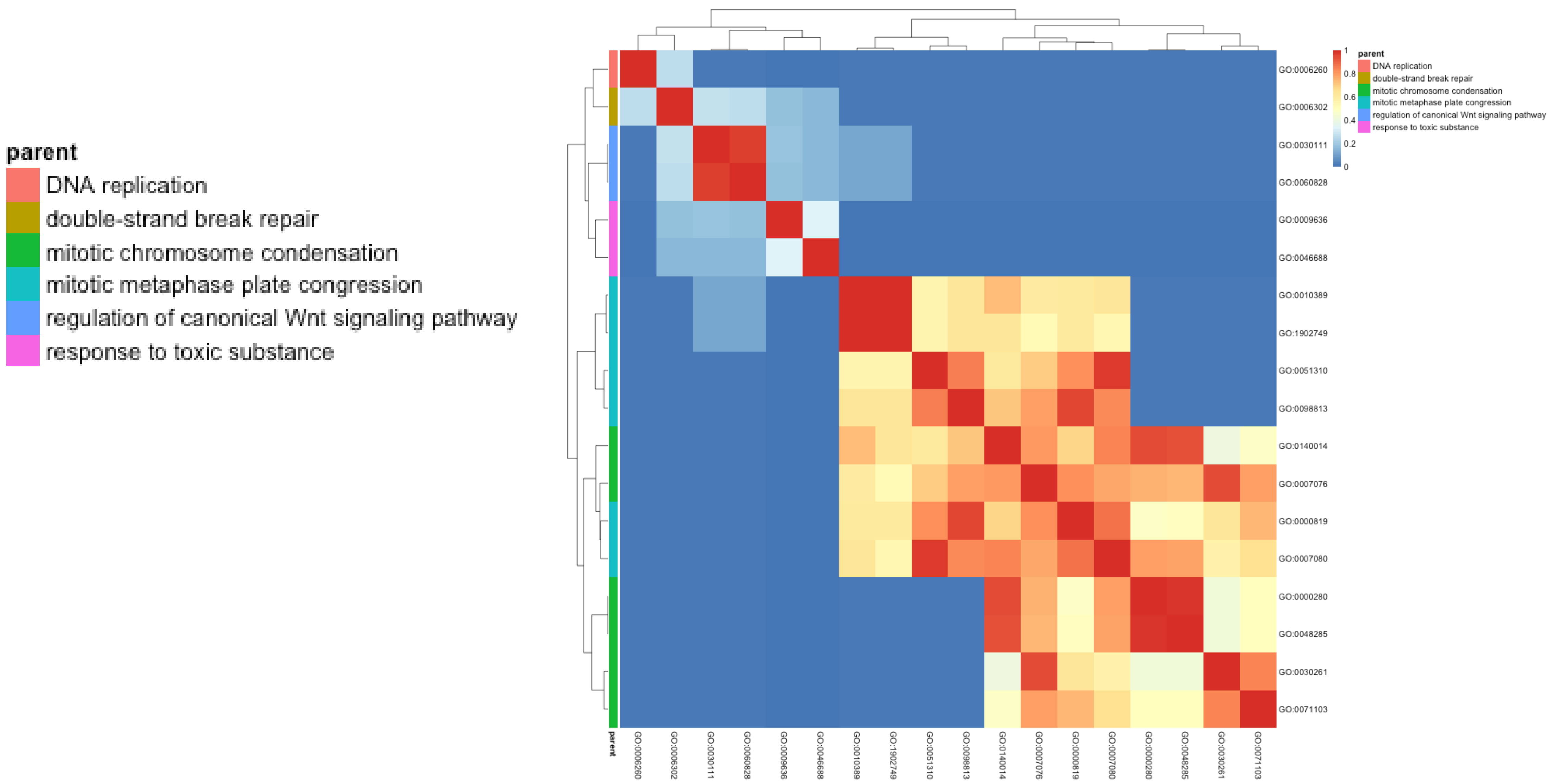

**Supplementary Figure S10c: Functional integration of miRNAomics with proteomics and transcriptomics analysis of senescent extracellular vesicles: functional reduction heatmaps**

#### A) Up-regulated miRNA/Protein

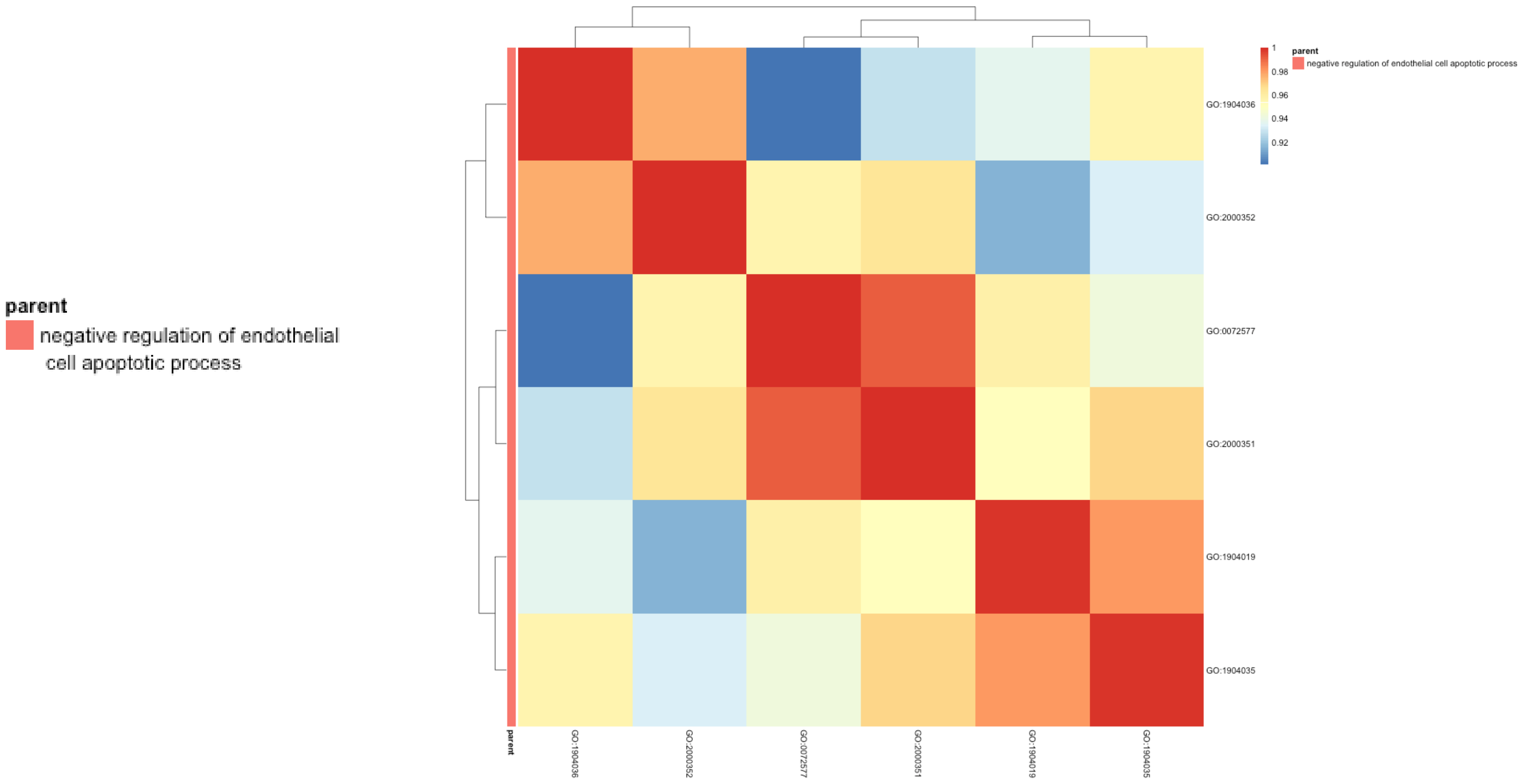


#### B) Down-regulated miRNA/mRNA

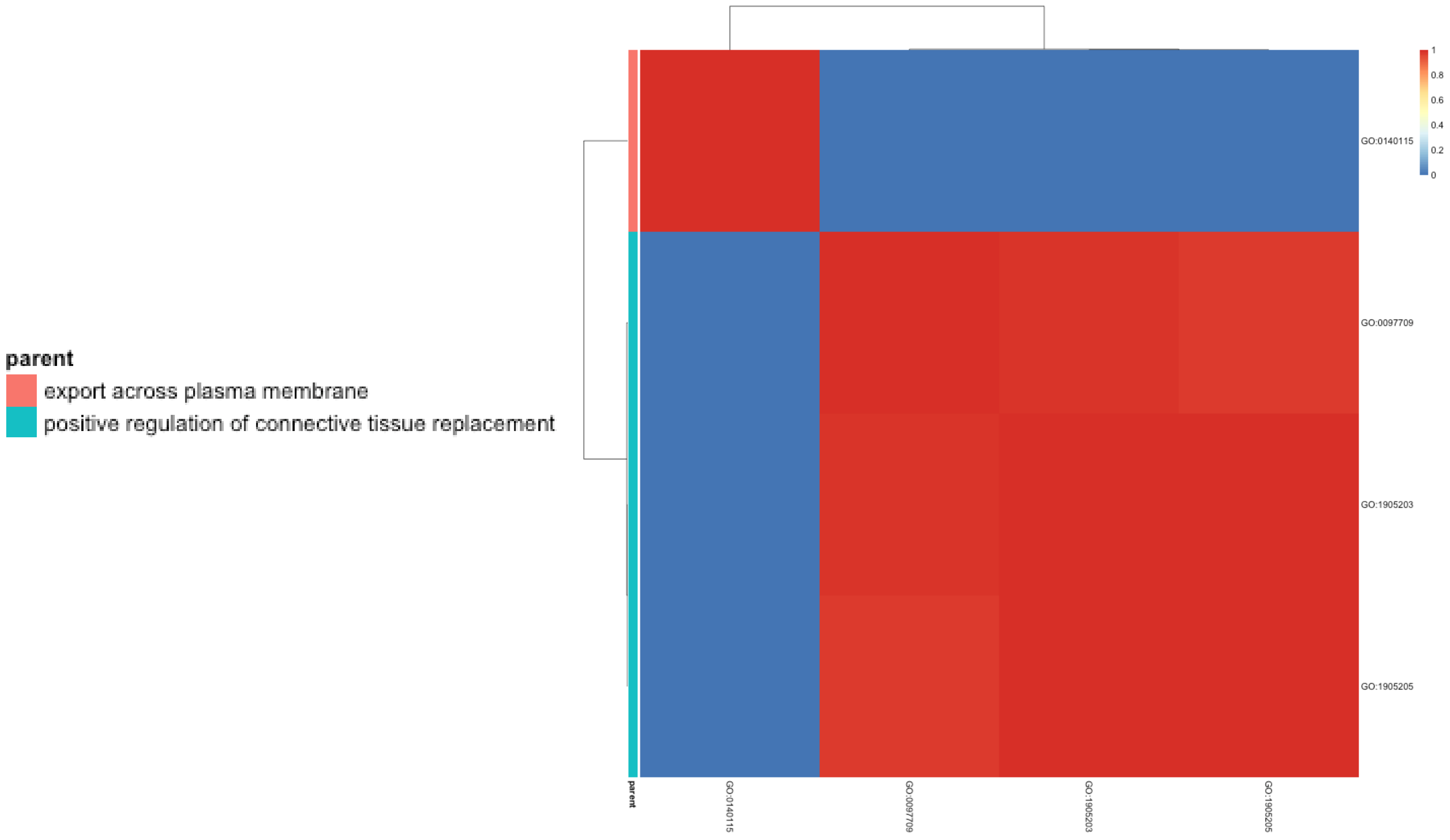

#### Supplementary Figure 10: Molecular integration of miRNAomics with proteomics and transcriptomics analysis of senescent cells: Enrichr Knowledge

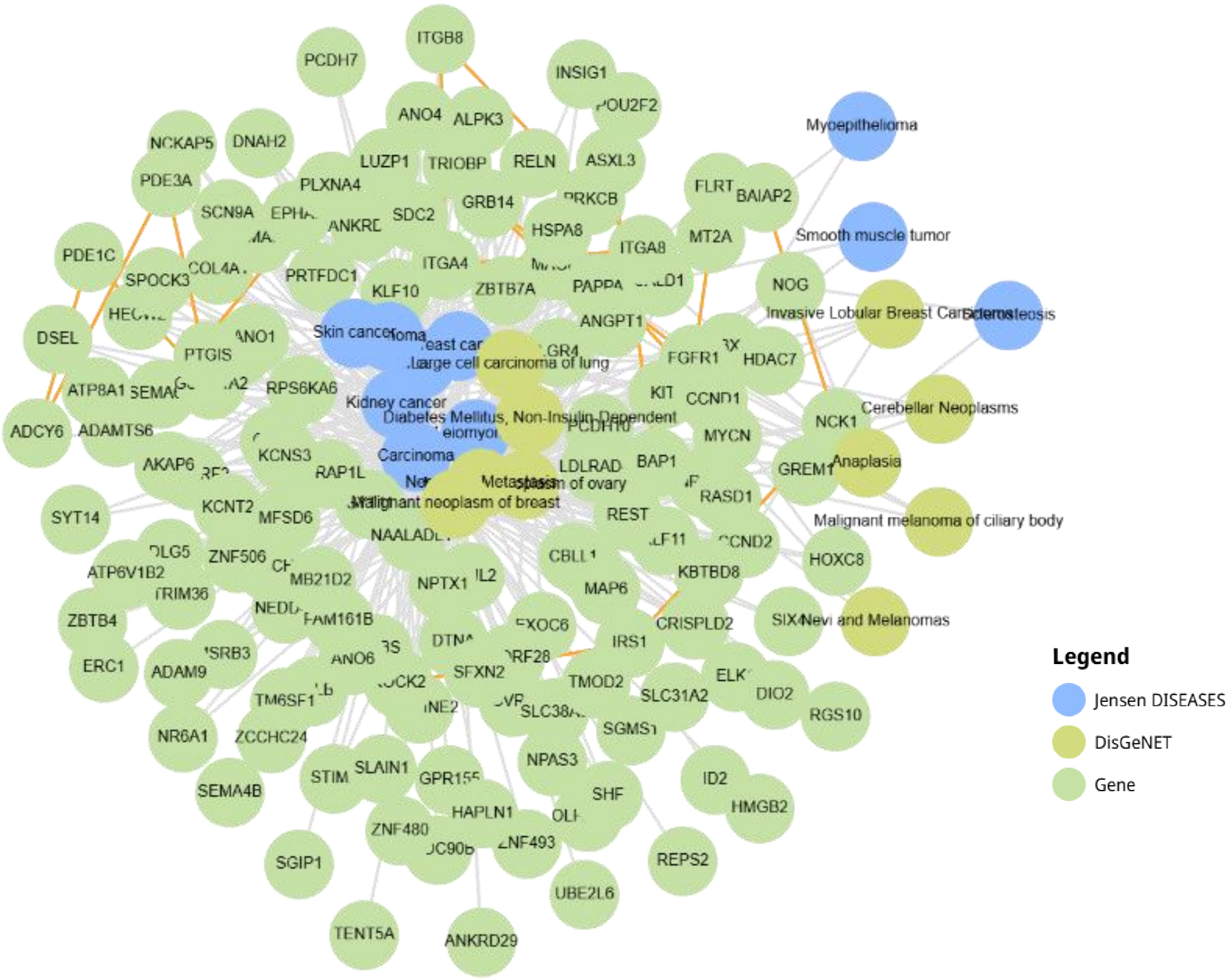
